# IDPConformerGenerator: A Flexible Software Suite for Sampling Conformational Space of Disordered Protein States

**DOI:** 10.1101/2022.05.28.493726

**Authors:** João M.C. Teixeira, Zi Hao Liu, Ashley Namini, Jie Li, Robert M. Vernon, Mickaël Krzeminski, Alaa A. Shamandy, Oufan Zhang, Mojtaba Haghighatlari, Lei Yu, Teresa Head-Gordon, Julie D. Forman-Kay

## Abstract

The power of structural information for informing biological mechanism is clear for stable folded macromolecules, but similar structure-function insight is more difficult to obtain for highly dynamic systems such as intrinsically disordered proteins (IDPs) which must be described as structural ensembles. Here we present IDPConformerGenerator, a flexible, modular open source software platform for generating large and diverse ensembles of disordered protein states that builds conformers that obey geometric, steric and other physical restraints on the input sequence. IDPConformerGenerator samples backbone phi (φ), psi (ψ), and omega (ω) torsion angles of relevant sequence fragments from loops and secondary structure elements extracted from folded protein structures in the RCSB Protein Data Bank, and builds side chains from robust Monte Carlo algorithms using expanded rotamer libraries. IDPConformerGenerator has many user-defined options enabling variable fractional sampling of secondary structures, supports Bayesian models for assessing agreement of IDP ensembles for consistency with experimental data, and introduces a machine learning approach to transform between internal to Cartesian coordinates with reduced error. IDPConformerGenerator will facilitate the characterization of disordered proteins to ultimately provide structural insights into these states that have key biological functions.

## INTRODUCTION

Disordered states of proteins, including unfolded states, intrinsically disordered regions (IDRs) of otherwise folded domains, as well as full intrinsically disordered proteins (IDPs) are increasingly recognized for the roles they play in folding kinetics^1^ aggregation propensity^2,3^, critical biological functions^4^, and pathological disease states^5^. Structural insights are needed to better understand these disordered protein states, and a variety of solution experiments have been developed to enable structural and dynamic descriptions of disordered proteins using nuclear magnetic resonance (NMR), small angle X-ray scattering (SAXS), single molecule fluorescence (SMF) and other data types^6–11^. However, solution experimental data for disordered states are averaged over a very large number of heterogeneous interconverting conformations, leading to greater challenges in structural interpretation than for folded proteins. Thus, specific computational approaches are required to bridge the gap between experiment and structural ensembles for disordered protein states.

The overall general approach begins with a large set of potential fractionally populated conformations and then either selects a subset of these and/or assigns weights to conformational sub-populations that best agree with the available, but highly averaged, experimental restraints. These two components have typically been considered separate problems and a number of methods exist for each. TraDES^12,13^, Flexible-meccano^14^, FastFloppyTail^15^ and other methods^16^ are available to generate conformer pools, based primarily on the statistical distributions found in folded protein structures from the RCSB Protein Data Bank^17,18^. TraDES builds trajectories of 3 Cα positions at a time based on the probabilities in a set of non-redundant structures from the PDB and then fills in the rest of the chain. Flexible-meccano builds chains by selecting φ/ψ torsion angles based on amino-acid specific conformational potentials derived from the PDB. FastFloppyTail also utilizes a three-residue fragment-based approach, with a bias towards the loop regions of the PDB. An approach from the group of Bernado^16^ similarly uses data from the PDB extracted as tripeptide segments. Another approach builds on Flexible-meccano but uses tripeptide conformers derived from MD simulations^19^.

Because the structural ensembles provided by these methods are agnostic to experiment, a separate step is used to select a subset of the conformations, or more generally reweight the conformers in the pool to define ensembles that best represent information about the disordered state from solution experiments. These include, among others, ENSEMBLE^20^, ASTEROIDS^20,21^, X-EISD^22,23^, BME^24^, and BW^25^/VBWSS^26^, with a number utilizing Bayesian statistics^22–24^ and/or maximum entropy^24,27^ to address inherent uncertainties in experiment and back-calculations from a disordered structural ensemble. Molecular dynamics simulations can provide conformers that are biased by the physicochemical interactions included in the force fields^28–31^, and represent Boltzmann weighted states, and have been used with NMR biases^2^ and a reweighted hierarchical chain growth algorithm^32^ to generate disordered ensembles.

Together these approaches have all been valuable for both creating and ultimately characterizing a variety of disordered proteins, however a number of challenges remain. In particular, for disordered proteins having preferential sampling of fractional secondary structure and tertiary contacts, especially for longer sequences, the starting pool sample becomes the more limiting factor for successful identification of subsets for reweighted ensembles that can fit experimental data. While a number of these tools can generate conformers biased by known secondary structure distributions, most of these tools are not flexible as to how users can generate disordered conformer ensembles, as well as evaluating them with respect to experimental data.

Here, we report the open source software platform, “IDPConformerGenerator”, for generating disordered protein conformations, utilizing a wide range of new and novel methods and models within a single software suite. IDPConformerGenerator begins with backbone builds based on torsion angle distributions of phi (φ), psi (ψ), and omega (ω) found in the RCSB Protein Data Bank (PDB) and then enables the build of side chain ensembles using Monte Carlo Side Chain Ensemble (MC-SCE) that completes the all-atom description by including hydrogens. IDPConformerGenerator has significant flexibility in user-defined options for size of peptide fragments used to build the backbone, amino acid substitutions, secondary structure biases, steric clash criteria, and energy biasing using force fields. Additional stand-alone and integrated algorithms within IDPConformerGenerator extend the fundamental internal coordinate conformer ensemble builds with state-of-the-art transformations to Cartesian coordinates using Int2Cart^33^, which yields more correct valence geometries and reduces steric clashes. Finally, the generated ensembles can be evaluated with stand-alone and integrated software modules such as the X-EISD Bayesian model for assessing agreement with many different experimental data types including NMR, SAXS, and single molecule fluorescence resonance energy transfer (smFRET)^22,23^. What makes IDPConformerGenerator distinct from other tools is its flexibility as a user-friendly toolkit to explore different computational strategies and protocols for rationally defining conformational ensembles of (intrinsically) disordered protein sequences.

We demonstrate that IDPConformerGenerator can efficiently calculate ensembles of proteins up to at least 440 residues in length with a variety of secondary structural distributions and tertiary contact patterns. Many of these have reasonable RMSDs from experimental solution data, particularly some generated with bias for fractional secondary structure based on NMR chemical shifts. These results support the utility of IDPConformerGenerator for creation of initial conformer pools that are more physically representative and more readily optimized by using experimental restraints with X-EISD^22,23^ or ENSEMBLE^20^.

## METHODS AND MODELS

### Design of IDPConformerGenerator

We set out to design a tool to efficiently generate conformers that realistically sample likely conformational space of intrinsically disordered sequences from statistical sampling of backbone torsion angles (φ, ψ, and ω) of short protein segments in the PDB that are identical or similar in sequence to the protein under investigation. This led to our choice to exploit the PDB for sampling of torsion angle space to generate more physically meaningful conformers, a choice also utilized by TraDES^12,13^, Flexible-meccano^14^, FastFloppyTail^15^ and others. Given these physically sound backbone conformations, we also provide side chain building algorithms such as MC-SCE that can generate ensembles of different rotamer states that are Boltzmann weighted and further all-atom representations by including hydrogens. The resulting sets of conformations are intended to be utilized as inputs to downstream approaches to define ensembles that best agree with experimental data, such as those that select subsets (e.g., ENSEMBLE^20^, ASTEROIDS^20,21^), re-weight conformers (e.g., BME^24^), or both (X-EISD^22,23^).

#### Building conformational ensembles

IDPConformerGenerator starts by creating a protein sequence database annotated with φ, ψ, and ω torsion angles and secondary structure per residue. We use non-redundant lists of structures such as those generated by Dunbrack PISCES database^34^. Hence, IDPConformerGenerator builds structures by extracting φ, ψ, and ω backbone torsion angles from the database, fragment-by-fragment (with fragments being peptides of variable length), using torsion angles matching the input sequence for each fragment or matching a user-defined residue tolerance (or substitution) dictionary. While other tools utilize rigid fragment sizes, IDPConformerGenerator allows users to configure the size and probability of the peptide fragments used to build the IDP chain stepwise, modulating the sampling strategy to explore. IDPConformerGenerator uses DSSP nomenclature^35,36^ to annotate residues by secondary structure elements. Because of this, users can define the secondary structural classes that IDPConformerGenerator will sample, either across the sequence or in a residue-specific manner, based on knowledge such as from NMR chemical shifts for fractional populations of secondary structures as a function of residue^37–39^. We provide several methods to sample bond geometries when converting from internal to Cartesian coordinates. The most important one is the recently published Int2Cart methodology^33^ that predicts bond lengths and angles for a set of torsion angles and residue identities. Instead of using hard-spheres to model atom volumes, IDPConformerGenerator computes the whole Lennard-Jones (LJ) potential to tolerate small clashes that can be compensated by favorable interactions, with user-defined thresholds to direct acceptance of a fragment or backtracking to rebuild. To generate full side chains, IDPConformerGenerator has integrated the Monte Carlo Side Chain Ensemble (MC-SCE) algorithm, originally developed for the more difficult case of folded proteins but which works easily for disordered states^40^.

#### Associated and Integrated tools

IDPConformerGenerator is designed as a platform to facilitate generation of disordered protein conformations, including analysis of resulting ensembles and scoring or re-weighting with respect to experimental data. Tools for analysis of structure are integrated within IDPConformerGenerator, including for secondary structure, torsion angle distributions, radius of gyration (Rg), end-to-end distances, asphericity (deviation from spherical shape of the conformers), and Cα - Cα distance and distance difference matrices. The software easily enables use of downstream tools for scoring, re-weighting or sub-setting to fit experimental data, and will serve as a future platform for integrating these tools, including the simple ENSEMBLE approach and the X-EISD Bayesian model.

### The IDPConformerGenerator software platform

To facilitate its development and use, IDPConformerGenerator is open source and extensively documented and the architecture is modular to allow easy extension with other modules and strategies (https://github.com/julie-forman-kay-lab/IDPConformerGenerator). IDPConformerGenerator is written in Python, and all its automated functionalities are available as command line commands. In addition, all IDPConformerGenerator’s functionalities are available through the Python interpreter and can be imported and used independently by more advanced users. Also, all IDPConformerGenerator pipelines are distributable across multiple CPU cores. IDPConformerGenerator’s software design facilitates a flexible approach to building conformers with numerous user parameters that enable very different realistic ensembles to be built, with the design philosophy and options discussed here. The overall workflow of IDPConformerGenerator is described in detail next, and is schematized in **Figure 1**.

**Figure 1.**
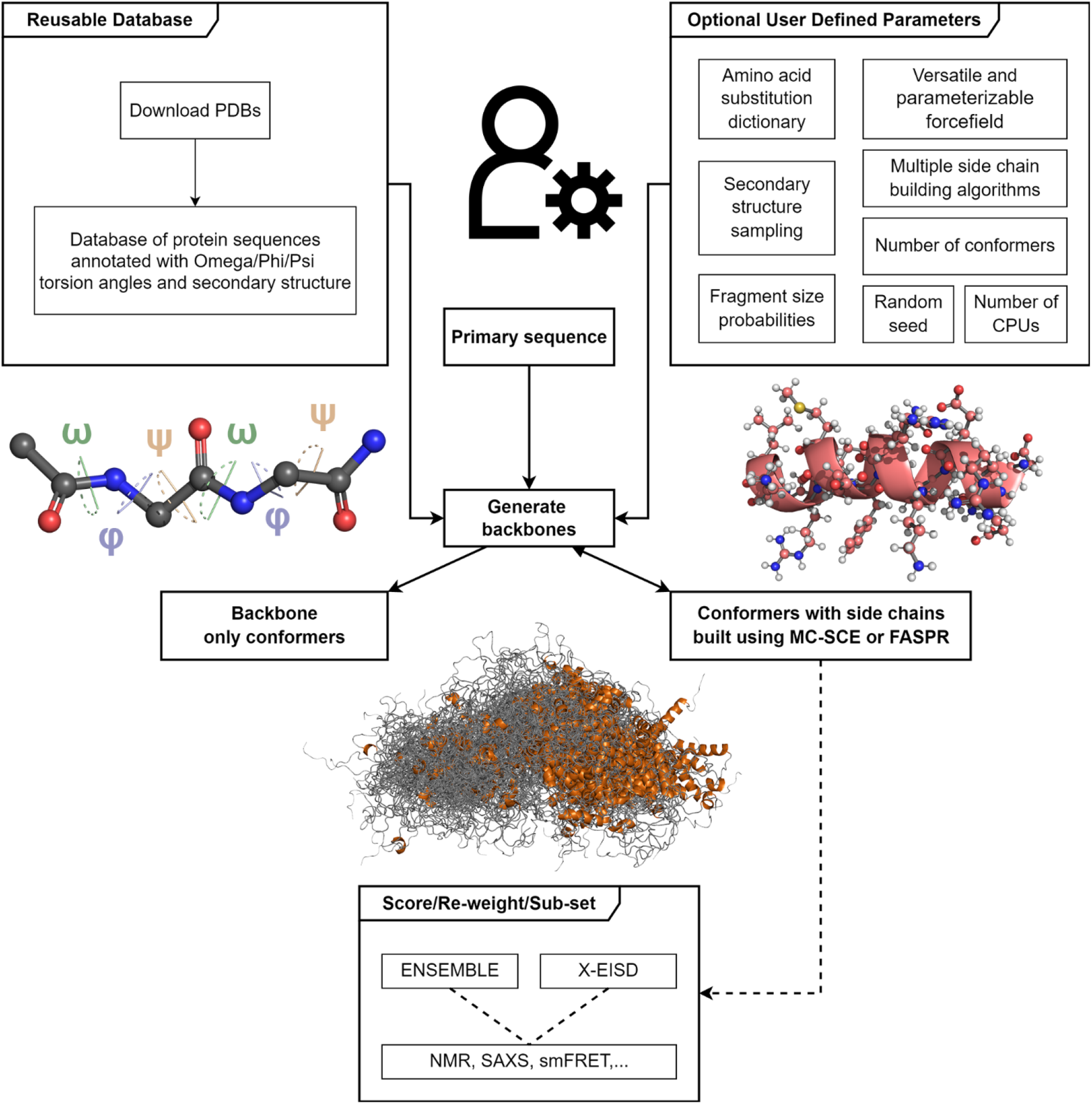
Schematic diagram of the IDPConformerGenerator approach. Generating conformers requires creation of a reusable database of backbone torsion angles and input of the primary sequence, with optional user-defined parameters including for amino acid substitutions, secondary structure sampling and fragment size probabilities. An example of a peptide of 2 residues (fragment size 2) that is used to build inhibitor-2 is shown with backbone torsion angles labeled, an helical secondary structure with all-atom side-chains of an I-2 conformer, as well as an illustrative set of 100 generated conformations for inhibitor-2. Conformers generated can then be scored or re-weighted based on experimental data.

### Backbone conformer generation

IDPConformerGenerator requires a torsion angle database from which to build backbone conformers. User can provide custom-made lists of the PDB chains to assemble the torsion angle database, thus allowing tuning for resolution and diversity. IDPConformerGenerator considers only continuous chains and selects the atoms with the highest occupancy for those with alternative locations. The IDPConformerGenerator parser works with both the older PDB format as well as newer mmCIF files. For the results reported here, we employed a fixed non-redundant database of PDB structures from the Dunbrack PISCES database^34^ (October 15th, 2020), including high-resolution structures with resolution better than or equal to 2.0 Å and R-factor of 0.25 or lower with a maximum mutual sequence identity of 90%.

Conformer backbones are built fragment-by-fragment, where fragments can be configured for different lengths (described below). Physical validation of the conformers (for example, steric clash check) is parametrizable via forcefields and threshold parameters. If a clash is found, IDPConformerGenerator will delete the last fragment and attempt a new one. If no new fragment can be added without steric clash, the building process will backtrack to delete additional fragments. This process is repeated until the whole backbone is complete. With the default parameters, some sequences can be built very fast, while others require extensive sampling times, also dependent on length (see below).

#### Including peptide bond ω torsion angle

One of the major differences between IDPConformerGenerator and previous backbone sampling tools is that IDPConformerGenerator includes peptide ω torsion angles in the sampling and building regime. The ω torsion angle is considered to be part of the torsion angle set for each residue in the order ω/φ/ψ. The decision to include ω is fundamental to our strategy to explore the IDP landscape by addition of multiple residue-sized fragments, and since ω angles can vary up to 20° in loop regions of high-resolution structures (**Figure 2**), including ω in the generator increases the accuracy of the extracted fragment. Note that this variation is not dependent on the resolution of the structure but that helices have narrower ω torsion angle distribution (**Supplementary Figure 1**). Accurate ω torsion angles are also critical for future incorporation of folded domains within otherwise disordered chains.

**Figure 2.**
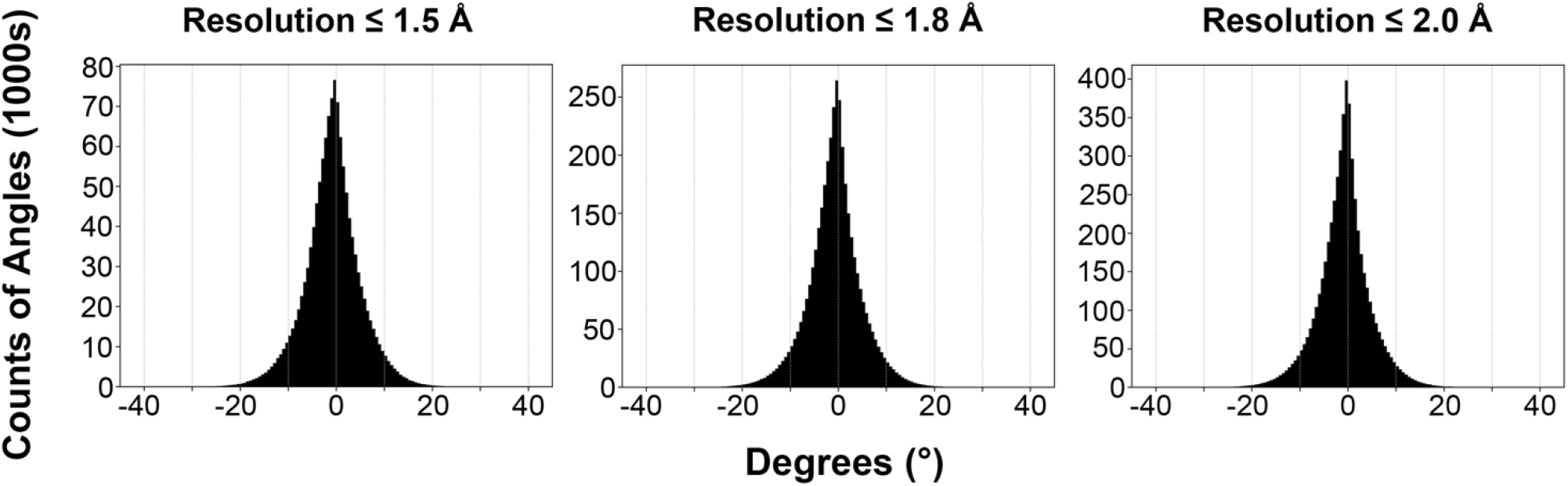
Histogram of ω (omega) dihedral angle distributions for structures found within the IDPConformerGenerator database. PDBs from the 24,003 PDBID database were used, with sets of this with resolutions better than 1.5Å (∼5,000 structures), better than 1.8Å (∼16,000 structures) and better than 2.0Å (full set). The deviations from an ω torsion angle of 180° (trans) is plotted centered around 0° to facilitate visualization of the distribution, rather than the actual ω angles. Cis peptide bonds are ignored for visualization purposes.

#### Sequence-specificity of chosen torsion angles

IDPConformerGenerator builds structures based on extracting backbone torsion angles from the torsion angle database, using torsion angles either (i) only from residues that exactly match the input sequence (when possible, see below) or (ii) from residues that match user-defined residue substitutions, i.e., isoleucine matching either isoleucine or leucine. The ability to explore a narrow or broader sequence space of the PDB with user-defined flexibility is an important feature of IDPConformerGenerator. Utilizing the exact sequence to find torsion angles in the PDB-derived database will guide IDPConformerGenerator to choose torsion angles that reflect the structural biases of that sequence. This capitalizes on the PDB-derived database as an “empirical force field” and is expected to generate local and secondary structure biases based on the sequence dependence of these structures. Enabling substitution of residues for similar residues, with a user defined substitution list, recognizes the availability limits of exact sequence matches in the PDB, and extends the potential torsion angles possible to be sampled.

IDPConformerGenerator has the ability to extract substitution lists based a table derived from the EDSSMat50 IDP-specific substitution matrix^41^ (see **Supplementary Table S1A**), with the user specifying the columns to be used to define how conservative or liberal to make the substitutions. Users can also provide specific substitution lists through the command line (with example shown) ‘-subs {“A”: “AG”}’, with substitutions described in the form of a Python dictionary where keys are the residues to replace, and values are the list of residues to include.

#### Peptide fragment sizes

IDPConformerGenerator enables users to define the size and probability of the peptide fragments used to build the IDP chain stepwise, modulating the sampling strategy to explore. By default, IDPConformerGenerator samples fragment sizes of 1, 2, 3, 4, and 5 residues with 10%, 10%, 30%, 30%, and 20% probabilities, respectively. Users can freely configure these values in any given manner. Therefore, IDPConformerGenerator can emulate previously published algorithms that build conformer chains by single residue or tripeptide additions while allowing full select countless other strategies. Shorter fragments (such as one residue at a time) disregard the sequence context of residue torsion angle frequencies, while with larger fragment sizes, information about cooperative structure (helical, turn, or extended strand-like elements) from the PDB is included in the growing disordered protein chain. In practice, fragment sizes up to pentapeptides are most valuable as it is hard to find sequence matches for longer segments, and cooperative structures found in disordered proteins generally do not extend beyond five residues (although there can be regions of longer cooperative helix). If a sequence match for the requested fragment size is not found, IDPConformerGenerator reduces the fragment size, one residue at a time, until a sequence match is found in the database, taking into account residue substitutions. Fragments are reduced until single residue additions are sampled. Regardless of the size of the fragment, if a proline residue in the input sequence follows the selected fragment, IDPConformerGenerator tries to expand the fragment sequence to include the proline as well because prolines impose severe torsion restraints on preceding residues, a strategy used by other disordered chain-generating tools.^16,42,43^

#### Secondary structural sampling

IDPConformerGenerator uses DSSP nomenclature^35,36^ and the IDPConformerGenerator-created torsion angle database can be configured to annotate residues by DSSP or any other third-party software (via the IDPConformerGenerator ‘sscalc’ interface) that classifies secondary structure elements. Because of this, the secondary structural classes of the PDB database that are sampled can be defined by users. A logical approach for generating disordered ensembles is to sample the loop regions of folded structures, which dominate most IDPs/IDRs. FastFloppyTail focuses on these regions and others have noted an increased ability to fit local structural experimental data to ensembles derived from the coil or loop regions of PDBs^44^, even though these regions can be poorly defined^45^. Using loops will bias to irregular structural elements found in the PDB that most likely represent disordered states, and also include short helical and extended elements. However, significantly populated helical elements and other secondary structural elements are found in disordered proteins states, so other DSSP codes are valuable to include. Therefore, IDPConformerGenerator allow users to sample any residues regardless of secondary structure annotation (“ANY” flag) or to specifically sample loops, helices, extended structures, poly-proline II helices or individual or combinations of any DSSP secondary structure code.

The secondary structure bias can be done uniformly across the sequence or in a residue-specific manner, using knowledge such as from NMR chemical shifts for fractional populations of secondary structures as a function of residue^37–39^. Depending on user choice, IDPConformerGenerator can prioritize building with the secondary structures of interest over the full sequence, leading to sampled fragments consisting only of residues matching a single secondary structure annotation, or can completely disregard secondary structures while building to allow the inherent secondary structure propensities observed in the PDB database to emerge. In this way, for example, ABC could be a fragment in which A and B residues are annotated as loop and C is annotated as a helix (with C being the first residue of a helix following a loop). Moreover, custom secondary structure sampling based on experimental knowledge of the fractional populations of secondary structures can be used, which can in turn override the database bias to build conformers to match known structural probabilities. For bias of secondary structure on a per-residue basis, building utilizes a custom secondary structure sampling database file containing fractional propensities for different secondary structures as a function of residue derived from NMR chemical shift data (using csssconv with δ2D^38^ or CheSPI^39^). If the chemical shift data are not available but other knowledge of sampling of helical or extended/β-strand regions exists (or if users want to explicitly define these), users can specify where significant sampling of helical or extended/β-strand regions are and sample the rest of the conformer without bias. Combining the rich ability to sample torsion angles from specific secondary structure annotations with the residue type substitutions that increase matching tolerance and specification of fragment sizes, users can sample highly restrictive or very broad conformational spaces.

#### Sidechain Building

During the backbone building process immediately after each backbone fragment is created, alanine sidechains are added onto all residues, except for glycines and prolines for which the full residue is added. These dummy alanines serve as coarse grain representations of the real sidechain. They avoid building backbone conformations that are too compact to fit sidechains without steric clashes, yet the volume of the alanine sidechain is small enough to allow the backbone to sample packed conformations, enabling exploration of sidechain packing.

However full side chains must be added, and IDPConformerGenerator adds sidechains using the Monte Carlo Side Chain Ensemble (MC-SCE) algorithm by sending backbone atom coordinates only and excluding any alanine and proline sidechains. The MC-SCE algorithm is a Monte Carlo approach for building side chain conformations on a predefined backbone structure^40^ that utilizes a convergent Rosenbluth sampling scheme and an augmented Dunbrack library for side chain rotamer sampling.^40^ The MC-SCE algorithm was originally written in Fortran but was fully rebuilt in Python to interface with IDPConformerGenerator to build side chain structures. Given a backbone structure, MC-SCE builds the side chains by aligning the backbone N, Cα and C’ atoms of the Dunbrack templates with the backbone from IDPConformerGenerator. The side chains are then rotated according to the sampled torsion angles, and this sampling procedure is the key to the Monte Carlo nature of the algorithm.

MCSCE can be used as both a stand-alone option (https://github.com/THGLab/MCSCE) and as two modes for working within IDPConformerGenerator. The simple mode provides an option for rapidly adding side chains to a backbone structure without introducing clashes, but the conformations might be energetically sub-optimal. Conversely, the exhaustive mode generates side chain conformations via user defined total number of trials for parallel execution of the building process, with the all-atom structure having the lowest energy of these returned to IDPConformerGenerator, but takes longer to run. (See Supplementary Information for more details.)

The FASPR^40^ algorithm to build side chain structures is also an integrated option. This stand-alone software for sidechain packing performs quickly for folded proteins. Note that FASPR does not include hydrogens, leading to a need to identify an optimal approach to build them afterwards. We have opted for MC-SCE as our preferred approach as it generates a complete all-atom description of conformers, including hydrogens, which is an important advantage over FASPR.

#### Internal to Cartesian Coordinate Transformations

The design of IDPConformerGenerator as a builder based on torsion angle sampling, rather than based in Cartesian coordinate space, has benefits and drawbacks. One clear benefit is that building with secondary structural biases, such as from NMR chemical shifts and backbone ^3^J-coupling data, is “native” to the builder. Building with tertiary contact biases, such as from NMR ^1^H-^1^H NOE or PRE data, is not. Importantly, energy calculations are made on Cartesian coordinates. In order to facilitate energy calculations, the conformers based on internal coordinates must be back-transformed to Cartesian coordinates.

The original approach used for most of the work reported here uses statistical sampling of bond angles for the set of matched fragments and average values for bond lengths based on the identity of the previous and next residue to the residue being built. The currently recommended approach which improves upon the aforementioned strategy uses Int2Cart developed by Li and co-workers^33^ a deep learning model that better predicts the correlations between the whole sequence, and bond lengths and bond angles for a given set of ω, φ, and ψ torsion angles to yield more accurate Cartesian coordinates. This very recent implementation can be used as both a stand-alone option (https://github.com/THGLab/int2cart) and is also integrated within IDPConformerGenerator directly.

#### Energy considerations

Instead of using hard-spheres to model atom volumes, IDPConformerGenerator computes the whole Lennard-Jones (LJ) potential to create conformers that are self-avoiding polymers. The default LJ parameters are the Amber14SB force field, which were used for the results generated here, but are also user-definable. Computing the whole LJ potential allows the building engine to tolerate small clashes that can be compensated by favorable interactions. and compensates for the fact that rigid conformers are created, i.e., no flexible minimization is performed at any stage. Severe clashes will, nonetheless, have a profound impact on the energy term, and thus the energy threshold for rejection can be defined by users, with higher values allowing exploration of broader conformational space. This feature is useful when modeling sequences with reduced representation in the database.

IDPConformerGenerator can build backbone-only or full sidechain-containing conformers. For this reason, two energy threshold parameters are implemented, one to control the tolerance for backbone atoms (”-etbb”), and another to control the energy threshold in all-atom conformers with sidechains (”-etss”). The energy thresholds to accept or reject a conformer can be calculated pairwise (atom-by-atom) or over the full structure, based on user choice. For each fragment built, the energy is computed; if below the threshold, the fragment is accepted, and otherwise, it is rejected. If sidechains are being built, once the backbone is complete, IDPConformerGenerator attempts to place the sidechains. If successful (energy term below threshold) the conformer is saved to disk. Otherwise, the backbone conformation is considered to be too restrictive to build sidechains, the whole conformer is discarded, and creation of a new conformer starts. Since the energy threshold for acceptance after sidechain addition is distinct from the threshold controlling backbone building, a user can accept all sidechain packing results by providing a large number for the sidechain energy threshold.

#### X-EISD Bayesian model

IDPConformerGenerator is designed as a platform and supports direct incorporation of calculated ensemble into downstream tools for scoring and re-weighting based on experimental data. The internal integration of these tools is envisioned in the near future. Of particular interest is X-EISD^22,23^, a method which calculates the maximum log-likelihood of a protein structural ensemble by accounting for the uncertainties of a wide range of experimental data and back-calculation models from structures. These data include NMR chemical shifts, J-couplings, residual dipolar couplings (RDCs), hydrodynamic radii, nuclear Overhauser effects (NOEs), and paramagnetic resonance enhancements (PREs), smFRET, and SAXS curves.^22,23^ We also have introduced new data types, R2 relaxation rates and S^2^ order parameters, for the selection of an IDP ensemble consistent with NMR dynamics data^46^. Given the ensembles created with IDPConformerGenerator, the X-EISD model can be used as a scoring function that helps reweight the IDP ensembles for best agreement with experimental data given the different experimental and back-calculation uncertainties.

#### Analysis tools

There are a number of commands currently integrated within IDPConformerGenerator that can enable analysis of resulting ensembles. These include analysis of the resulting fractional secondary structure. In addition, a set of complementary analysis tools were utilized to ask specific research questions regarding the ensembles (see below, available as stand-alone scripts). These include the root-mean-squared deviations (RMSDs) from experimental data restraints and ENSEMBLE and X-EISD scores; pair-wise RMSDs of atomic coordinates; measures of local structure: secondary structure, φ/ψ/ω distributions; measures of hydrodynamic properties: radius of gyration (Rg), end-to-end distances, asphericity (deviation from spherical shape of the conformers); and measures of tertiary contacts: Cα - Cα distance and distance difference matrices.

*<u>Additional user-defined parameters</u>* are available, including random seeds to control reproducibility. IDPConformerGenerator runs are deterministic, i.e., the same results can be achieved by providing the same initial database, the same input parameters and the same random seeds, on the same machine. Users can also specify the number of cores of a multi-processor computer.

## RESULTS

In order to demonstrate the utility of IDPConformerGenerator and ask questions regarding the optimal parameters for building diverse and physically meaningful ensembles, we have used a set of intrinsically disordered proteins (IDPs) of various sizes: Sic^47^, alpha-synuclein^48^, inhibitor-2^49^ and Tau^50^, as well as the unfolded state of the N-terminal SH3 domain of the Drosophila signaling protein Drk (drkN SH3)^51^.

- Sic1 is a yeast cell-cycle regulator that inhibit a cyclin-dependent kinase and is degraded following ubiquitination due to binding the ubiquitin ligase substrate-binding domain (Cdc4 WD40 domain) in a dynamic complex dependent on multi-site phosphorylation^47^. The N-terminal 92 residues of Sic1 are necessary and sufficient for binding and have been extensively characterized by NMR, SAXS and smFRET, and this fragment is therefore used here^6,52–54^.
- Human alpha-synuclein (aSyn, 140 residues) is highly abundant in the brain where it is found largely in the axon terminals of presynaptic neurons to regulate synaptic vesicle trafficking and subsequent neurotransmitter release^48^. In the presence of membrane vesicles (or other lipid environments), alpha-synuclein forms a helical structure, but in the absence of lipid it is highly disordered with minimal propensity for helical or other secondary structure. It has been studied in both states, but for testing purposes we utilize NMR and SAXS data from the disordered state^15,21, 55–60^.
- Inhibitor-2 (I-2, 159 residues) is an inhibitor of protein phosphatase 1 (PP1), forming a dynamic complex with PP1 that only orders a limited portion of I-2 upon binding, based on crystallographic data^49^. In the absence of PP1, I-2 is disordered yet has significant population of helices, based on characterization by NMR^61,62^.
- Tau (microtubule-associated protein tau) is a 758-residue IDP with numerous functional annotations, including promotion of microtubule assembly and stability and roles in establishing and maintaining polarity of axons in neurons^50^. It is an RNA-binding protein that phase separates in vitro and is found in cellular biomolecular condensates, consistent with its lower complexity sequence^63^. We use the first 441 residues as a test system since a fragment encompassing these residues has been studied using NMR^64,65^. Short Tau peptides have also been studied^15^ and we similarly utilize a Tau peptide as a test system.
- Finally, the drkN SH3 domain exists in a dynamic equilibrium between folded and unfolded states, with the unfolded state extensively studied as a model disordered protein for development of ensemble calculation methods due to the large number of experimental NMR, SAXS and smFRET restraints available and its small size (59 residues)^22,46,62,66^.

Sequences for these proteins and fragments are given in **Supplementary Table S1B**. Note that there are some peptide sequences of aSyn and Tau in the PDB database we use (**Supplementary Table S1C**), many in complex with antibodies. It may be valuable to include structures of the protein of interest or homologous proteins, such as complexes of folded proteins with the disordered protein of interest (or its fragments), to provide conformations that are likely to be sampled at some level. Alternatively, users may choose to exclude such structures to avoid potential bias. Either approach is possible because IDPConformerGenerator allows users to assemble custom-made databases of torsion angles from user-defined input PDB lists. The number of sequence matches for different fragment sizes of the drkN SH3 domain in our database for different secondary structure sampling, including exact matches or with substitutions, is given in **Supplementary Table S1D** to provide concrete examples of how torsion angles are chosen in IDPConformerGenerator. The ‘stats’ sub-client calculates the sequence matches in the database, for an input sequence and considering the input parameters of the building process. In this way, users can easily assess the number of angles available for each chunk and identify possible bottle necks were residue tolerance might be needed.

Here we characterize multiple aspects of IDPConformerGenerator: computational speed for generating ensembles, the diversity of conformer sampling, the presence or absence of secondary structure (especially helical fraction), how well these unoptimized disordered ensembles recapitulate experimental data, and comparisons to other structural ensemble generators such as TraDES and FastFloppyTail.

### Computational Timings

The speed of conformational ensemble generation is significant, particularly for larger proteins, as a large and conformationally diverse input pools are valuable for further reweighting or sub-setting. We compared speeds for generating disordered conformers using IDPConformerGenerator with different fragment sizes and different secondary structure option for all of the test systems. For these, we generated backbones for each protein, a 100 kJ backbone energy threshold and MC-SCE for side-chains. The goal was to yield 1000 successful full conformers for each protein such that timings were normalized on a per successful conformer basis. Exact timings and percentage of successful conformers from the generated backbones can be found in **Supplementary Table S2A**.

**Figure 3A** demonstrates the general trend of faster conformer generation for shorter length chains, as expected, with a non-linear dependence. Building with only helices or extended strands is usually faster than building with loops or mixtures of loops with helical or extended structures, such as with CSSS or ANY, as loops increase the likelihood of steric clashes and difficulties in sidechain packing, although helices and strands are not as representative of disordered states (**Supplementary Table S2A**). As shown in **Figure 3B**, increasing the fragment size significantly increases the speed of conformer generation for the proteins investigated, and varying the secondary structure sampling method alters the speed for different fragment sizes. In most cases, using substitutions was also found to be faster, likely due to more fragment matches in the database. Overall, conformer generation is reasonably efficient but strongly dependent on chain length, with speeds of 400-500 conformers per hour per computer node for the drkN SH3 domain unfolded state (59 residues, res), 200-275 for Sic1 (92 res), 50-100 for aSyn (140 res), 40-75 for I-2 (159 res) and about 5 for Tau (441 res), using one node on the Graham supercomputing resource. For this, and all other calculations unless otherwise specified, we used one node of the Graham resource of Compute Canada (now Digital Research Alliance of Canada) with 2x Intel E5-2683 v4 Broadwell @ 2.1GHz CPUs and with 125GB of RAM per node.

**Figure 3.**
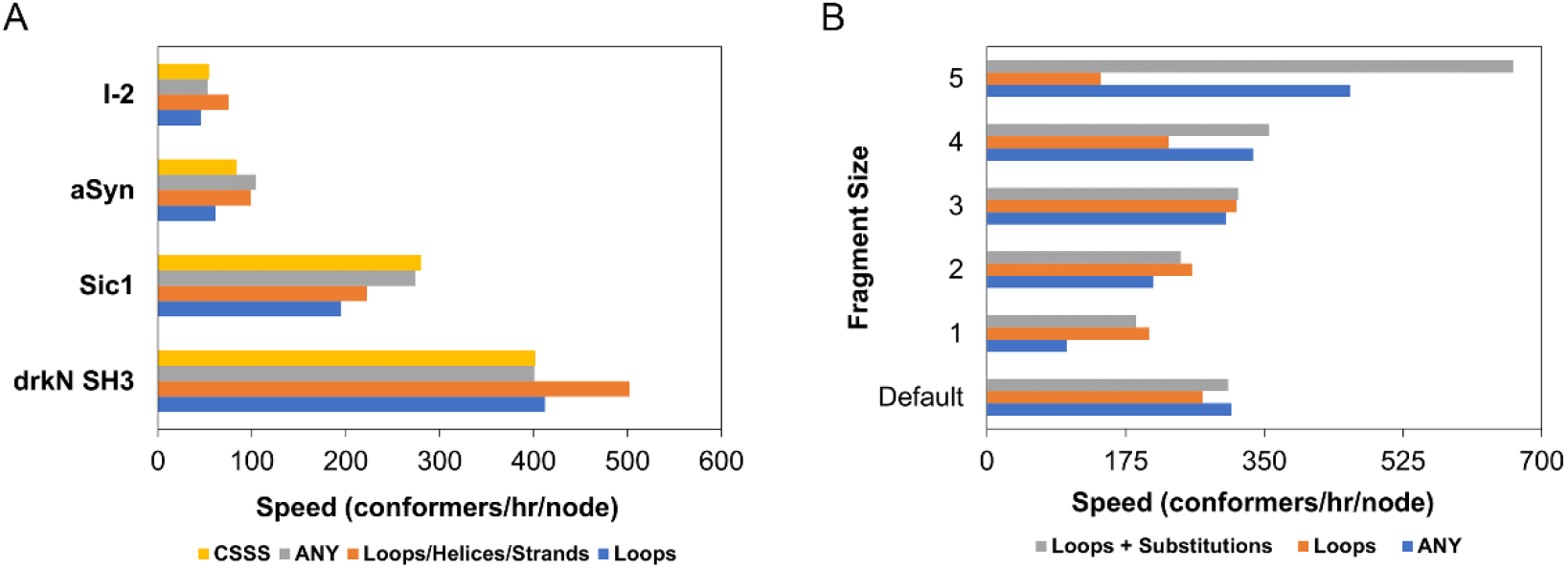
Timings for IDPConformerGenerator with MC-SCE and dependencies on fragment length, secondary structure sampling, and protein test system size. Speed is defined as number of conformers per hour. **(A)** Speeds for variable secondary structure sampling methods on IDPs with different lengths, shown for sampling with custom secondary structure propensities (CSSS, yellow), “ANY” secondary structure (grey), different combinations of loops, helices, and strands (orange), and only loops (blue). Speeds are shown for selected ensembles of the drkN SH3 domain unfolded state (59 amino acids, aa), the Sic1 N-terminal targeting region (92 aa), alpha-synuclein (aSyn, 140 aa), inhibitor 2 (I-2, 159 aa). Speeds for all conformer ensembles generated including for the Tau fragment (441 aa) are in Supplementary Table S2A. **(B)** Speeds for variable fragment sizes and secondary structure sampling methods for Sic1 are shown for sampling with only loops with substitutions (grey), only loops without substitutions (orange), and “ANY” secondary structure (blue). Default fragment length probabilities are 0.1, 0.1, 0.3, 0.3, 0.2 for fragment lengths of 1, 2, 3, 4, 5, respectively.

We also tested the intersection of the impact of sequence length and the diversity of amino acid residues in the sequence on the speed of conformer generation. Low complexity IDRs using fewer amino acids are increasingly understood to have a functional role in facilitating phase separation within biomolecular condensates or membrane-less organelles^67,68^. Of our test proteins, the Tau fragment is the longest (441 res) and is known to phase separate^63^. It is also lower in complexity than the other test proteins, with the first 300 residues annotated as having compositional bias by CAST and being low complexity by SEG, respectively^69,70^. Such low complexity sequences are not found in the folded proteins in our database and we explored if they would take longer to build. We quantified speed of conformer generation in minutes per amino acid on the multiprocessor server. Tau was segmented into three segments of 147 res to compare with I-2 (159 res) and aSyn (140 res), and five segments of ∼90 residues in order to compare with Sic1 (92 res). We found that the central 147-res segment of the Tau fragment was the fastest to build, but that there were no clear trends on the basis of complexity when comparing Tau to aSyn or I-2 (**Supplementary Figure S2**).

The sidechain addition step is much longer than backbone generation, with our preferred sidechain packing algorithm MC-SCE taking a larger fraction of the time as chain length increases. MC-SCE was initially optimized for packing sidechains onto the backbone of folded proteins. Although the success rate decreases with longer backbone lengths, we found that the settings in MC-SCE could be optimized for IDPs by reducing the number of attempts/trials spent on packing sidechains onto backbones from 128 to 32. For Tau, using 32 trials increased the speed per conformer by 3 to 4.4 times depending on secondary structure sampling (**Supplementary Table S2B**). Another observation based on these benchmarks is that the success rate increases with an increased number of backbone conformers available as input to MC-SCE.

For methodological purposes, we also asked what the optimal energy flags for speed of calculation of conformers that do not have significant steric clashes in order to facilitate rapid building of structural ensembles. We built sets of 1000 backbone conformers of I-2 with loops or other secondary structure sampling with either 100 or 250 kJ pairwise energy thresholds and used MC-SCE for sidechains, with average times of 68 and 38 minutes, respectively. The ∼78.9% increase in time for the 100 kJ threshold only led to an ∼10% gain in clash free conformers. We observed similar results for aSyn, with average times of 45 and 25 minutes, respectively, and an ∼80.0% increase in time for the 100 kJ threshold and only an ∼16% gain in clash free conformers. Thus, increasing the energy threshold can speed up the full conformer generation time for proteins at least as long as I-2. (**Supplementary Table S2C**).

### Sampling depth

Next, we interrogated the depth of the torsion angle space in ensembles built from torsions derived from the PDB dataset. When building proteins with a specific sequence, particularly for fragment sizes of 3, 4 and 5, the finite size of the PDB-derived database leads to minimal torsion angle options as only sequence matches of the defined fragment size can be used to build. This leads to what we call torsion angle bottlenecks for specific residues. **Figure 4** shows the φ distributions for Sic1 generated using loops with various fragment sizes, demonstrating decreasing numbers of distinct torsion angles as fragment size increases. For fragment sizes of 5 between residues 20-30 essentially one set of backbone torsion angles was used over all these 1000 structures. **Supplementary Figure S3** shows histograms of how many segments of the drkN SH3 domain sequence for various fragment sizes are present in the database, demonstrating the minimal data for fragment sizes of 6 and 7, with values also provided in **Supplementary Table 1D**. In order to avoid such torsion angle bottlenecks, mixtures of fragment sizes are optimal when requesting exact sequence matches. The default of probabilistic sampling of fragment sizes of 1, 2, 3, 4, and 5 in the ratios of 1:2:3:3:1 enables contribution from larger fragment sizes with cooperative structural elements while minimizing torsion angle bottlenecks, as seen in the bottom rows of **Figure 4**. Using substitutions can help avoid bottlenecks, with the right panels of Figure 3 showing greater torsion angle coverage than the left. Increasing the number of DSSP codes utilized can also be beneficial (**Supplementary Figure S4)**, as using only helices or only strands yields limited torsion angle sampling (and is not realistic for disordered chains). Being agnostic to secondary structure annotation is another approach, as seen for the difference between using loops only or all possible annotations for the drkN SH3 domain sequence (**Supplementary Figure S5**).

**Figure 4.**
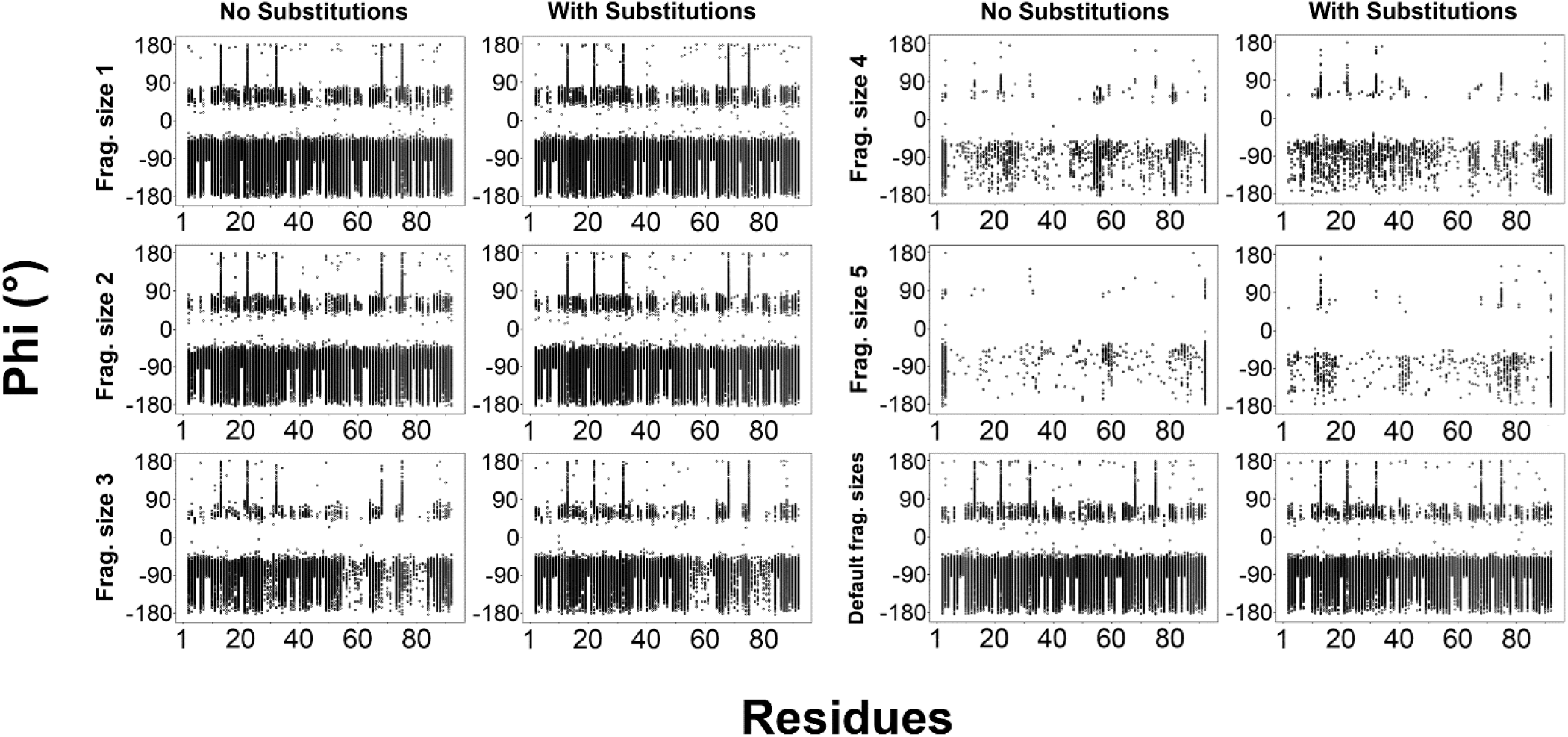
Phi (φ) torsion angles in Sic1 ensembles sampled using different fragment sizes, with and without substitutions. Calculations were for 1000-conformer ensembles generated by sampling loops only, with fragment sizes of 1, 2, 3, 4, 5, and default fragment size probabilities. The left and right columns are for the Sic1 sequence without or with substitutions, respectively. Substitutions are derived from columns 5, 3, and 2 of the EDSSMat50 amino acid substitution matrix. Plot generated with the ‘--plots’ flag in ‘idpconfgen torsions’ CLI.

We then looked for the optimal parameters (fragment size, secondary structure flags) for increasing diversity of calculated structures, as measured by average pair-wise RMSDs, hydrodynamic parameters, and asphericity. Rg, end-to-end distance (Ree) and asphericity are all measures of the shape of a conformation, with smaller Rg and Ree values and asphericity approaching 0 implying more spherical, compact chains, while large values reflect irregular, less compact shapes. As expected, we note that it is critical to incorporate loop regions to build diverse structural ensembles of disordered protein states, since with only helical or extended DSSP flags, long helices or strands are built, not representative of disordered conformations (**Figure 5**, **Supplementary Figure S6 and Supplementary Table S3**). This is seen by the much higher Ree and asphericity values, such as for those built with helices only having asphericity values of >0.8 and with strands only having Ree values 1.5 to 2 times as large as for those built with loops. To further increase the diversity of calculated structures, the “ANY” secondary structure flag is optimal, as it will use the natural secondary structure propensities of the entire PDB database and not limit to user-defined secondary structures that restrict sampling of conformational space (see below).

**Figure 5.**
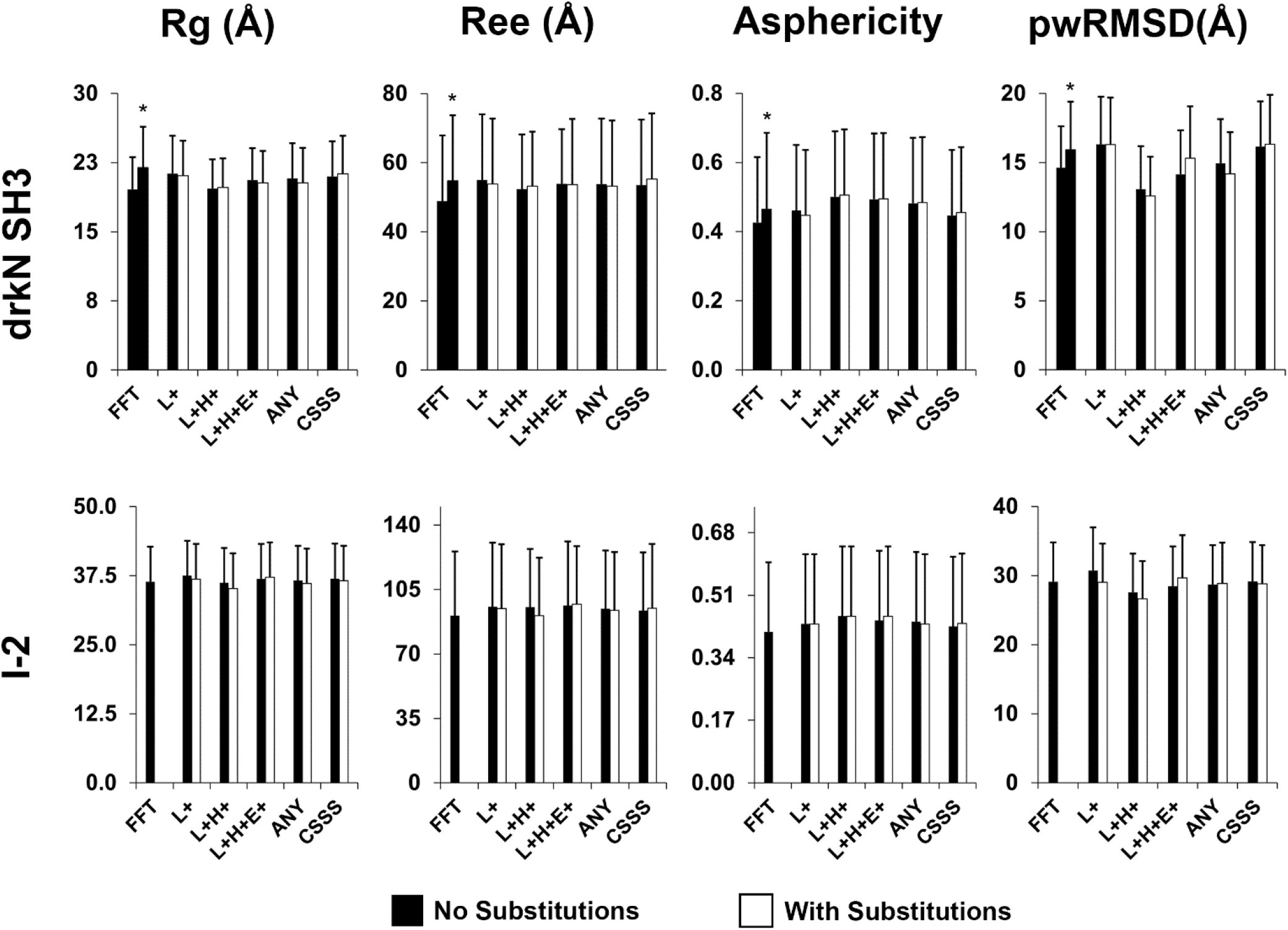
Diversity analysis of conformational ensembles of the drkN SH3 domain unfolded state and I-2. Radius of gyration (R_g_), end-to-end distance (R_ee_), asphericity (A) and pairwise root-mean-squared-deviations of atomic positions (pwRMSDs), are shown as a function of secondary structure sampling parameters for 1000-conformer ensembles generated with different secondary structure sampling, including loops (L+), loops and helices (L+H+), loops, helices and extended strands (L+H+E+), all torsion angles agnostic to secondary structure (ANY) and biased by δ2D chemical shifts (CSSS), or with FastFloppyTail (FFT), for the drkN SH3 unfolded state (row 1) and I-2 (row 2). Standard deviations for R_g_, R_ee_, A and pwRMSD are also shown as bars. Supplementary Figure S5 shows similar data for other protein systems. * is for the standard protocol which for this case treats the protein as a mixture of ordered and disordered, while the other is for a modified protocol in which the protein is considered to be fully disordered.

Plotting pairwise RMSDs as a distribution (**Figure 6**) demonstrates that the ensembles are smoothly sampled, with no significant clusters of similar structures, consistent with our goal of generating diverse conformers. Varying secondary structure sampling approaches can also increase the variety of conformational space explored, as the custom secondary structure sampling shifts the RMSD histogram to larger values. As seen in plots of pairwise RMSD distance matrices (**Supplementary Figure S7**), no regions of lower pairwise RMSDs are seen, indicating that the generated conformers have large variability in Cα backbones. Pairwise RMSD is correlated to protein length, with RMSD values ranging from above 5Å to 30Å for the shorter disordered drkN SH3 domain unfolded state (59 res) and from 15Å to above 50Å for I-2 (159 res), indicating significant heterogeneity in conformational sampling.

**Figure 6.**
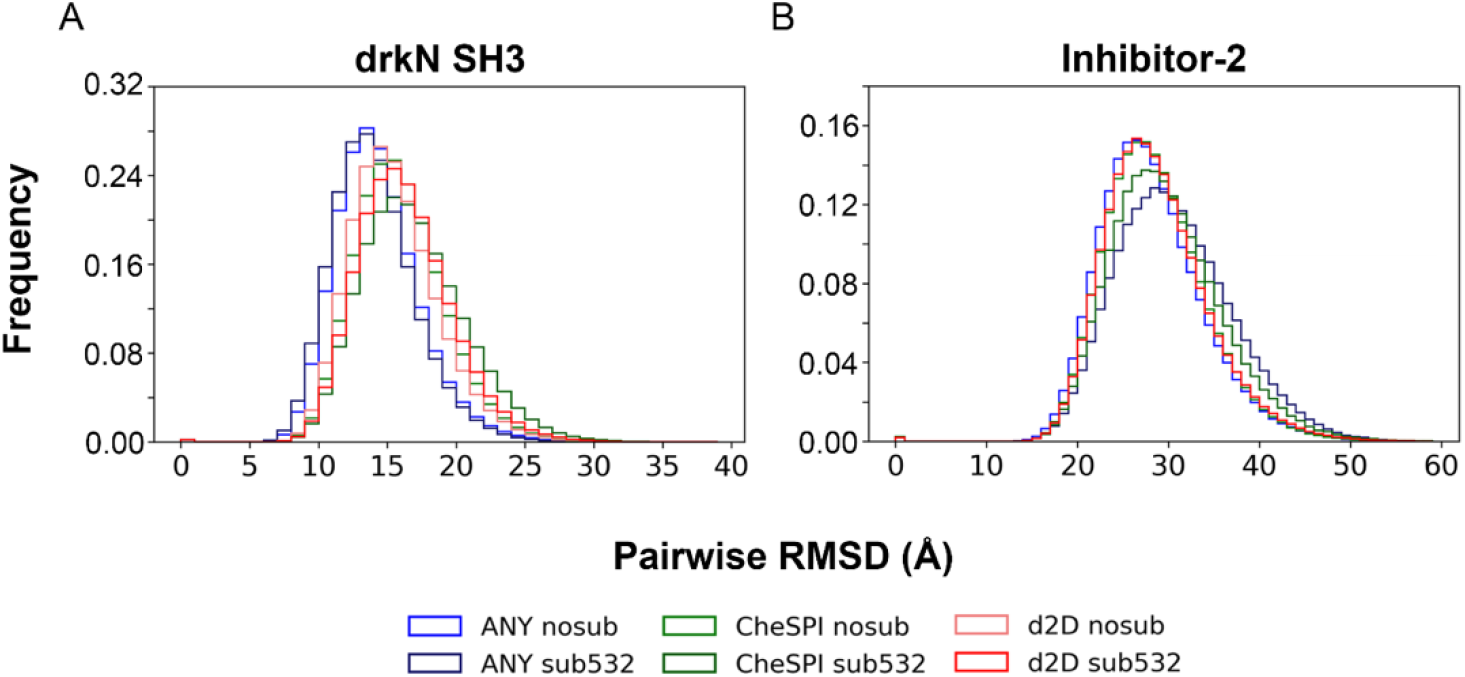
Pairwise RMSD distributions for ensembles of the (A) drkN SH3 domain unfolded state and (B) I-2. Calculations were for different ensembles of 1000 conformers each, plotted with bin sizes of 5Å. “ANY” indicates sampling the database without biasing secondary structures, “nosub” indicates no substitutions, “sub532” indicates amino acid substitutions from columns 5, 3, and 2 of the EDSSMat50 amino acid substitution matrix, and “CheSPI” or “δ2D” indicates custom secondary structure sampling (CSSS) pools biased by CheSPI or δ2D estimations of secondary structure propensities.

We were also interested to see if IDPConformerGenerator is able to effectively capture local structural changes with amino acid sequence changes. We used a 16-mer peptide from the Tau K18 fragment previously studied by Stelzl and coworkers; reweighted hierarchical chain growth was used to generate Tau ensembles recapitulating structural details that were identified by NMR to have microtuble binding capacity and that are lost upon mutation of position P301^32^. To investigate the conformation diversity explored by IDPConformerGenerator and the variation in conformations for single-site mutations, we generated sets of 10,000 conformers for wild-type (WT), P301L, P301S, and P301T for the Tau fragment sequence: DNIKHVP^301^GGGSVQIVY. We sampled considering only sequence matching, disregarding secondary structure annotations, and allowed no residue substitutions for sequence matching.

One of the structural parameters explored in the Stelzl study is the distance between V300 O and G303 N. **Figure 7A** shows distributions for this O-N distance for the different variants. In agreement with the Stelzl study, we observe a considerable fraction of conformers for the WT with distance below 4Å, reflecting a turn structure and likely hydrogen bond, while for mutants these occurrences are much rarer. Each mutant reveals different patterns of O-N distances, showing that IDPConformerGenerator can capture local conformational diversity from single point mutations and that these will be incorporated into the larger disordered chain. **Figure 7B** shows the torsion angles for residue 301 in the variants. Here, we also observe very different profiles. Note the presence of conformers with a cis-prolyl peptide bond for P301 reflecting the natural tendency of cis-Pro in the context of this sequence but absent in the mutants lacking proline. These results demonstrate that IDPConformerGenerator can effectively sample realistic local conformations in a sequence-specific manner, consistent with its design. Another approach to sampling particular turn types or other structures that is available with IDPConformerGenerator is to utilize a amino-acid substitution dictionary to incorporate residues with known propensities for these structures.

**Figure 7:**
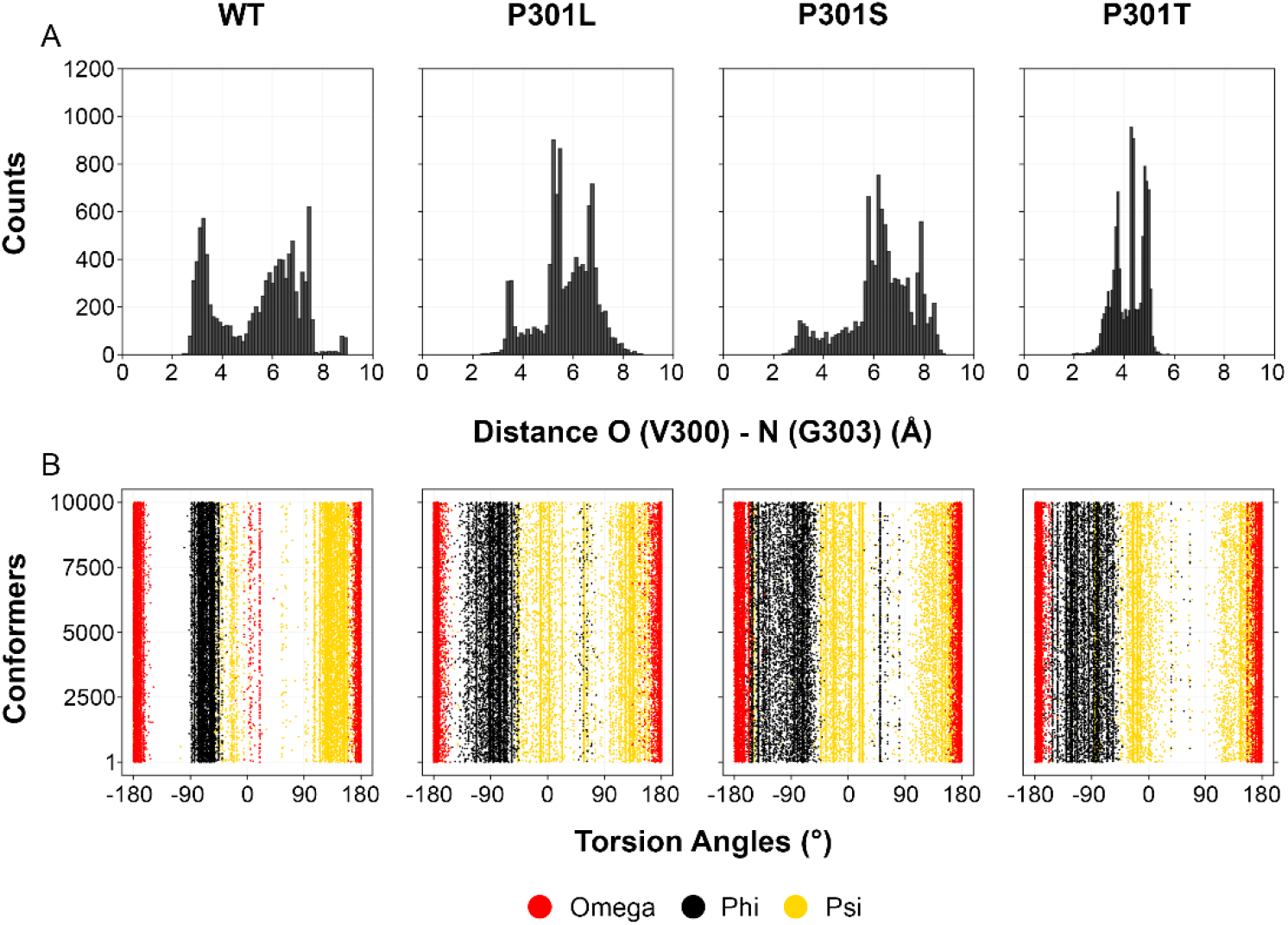
Local structural variations between the Tau K18 16-mer WT and different mutants. (A) Distribution of the distance between V300 O and G303 N atoms in 10,000-conformer ensembles generated with no substitutions. (B) Torsion angle distributions for position 301 of the different conformers in these WT and mutant ensembles, with ω representing the torsion angle N-terminal to the φ, as is our convention (typically denoted as ω of the preceding residue).

### Fractional secondary structure

An obvious question regards the impact of the secondary structure flags on the ultimate fractional secondary structures in the ensembles built. We generated 1200-conformer ensembles of Sic1 using different combinations of secondary structure sampling with loops (activated by default), helices, and strands. IDPConformerGenerator can pool together DSSP codes T (hydrogen-bonded turn), S (bend), B (β-bridge), P (PPII helix), I (pi-helix) and ’ ’ (blank, loops/irregular) as loop (L), H (α-helix) and G (3-10 helix) as helix (H), and only E (extended strand, participating in β-ladder) as strand (E). For all our calculations, we utilized this pooled set of DSSP codes. We generated Sic1 ensembles due to its lack of inherent significant biases in secondary structural propensity^52,53^ (**Figure 8**). Restricting to loops only or loops and strands, similar sampling is observed with greater sampling of the β region of the Ramachandran diagram, but since there are no hydrogen-bonded interactions, DSSP catalogs these as loops. There is also significant sampling of the α region and some low amounts of cooperative helix observed. With strands only, sampling of the β region of the Ramachandran diagram is dominant, with no strands defined by DSSP, again due to lack of hydrogen bonds. Restricting sampling to helices, however, leads to dominant sampling of the α region of the Ramachandran plot and, as expected, these structures show up as α-helix as defined in the DSSP. With loops and helices there are also significant cooperative helices generated. Similar results were seen for the drkN SH3 unfolded state (**Supplementary Figure S8**).

**Figure 8.**
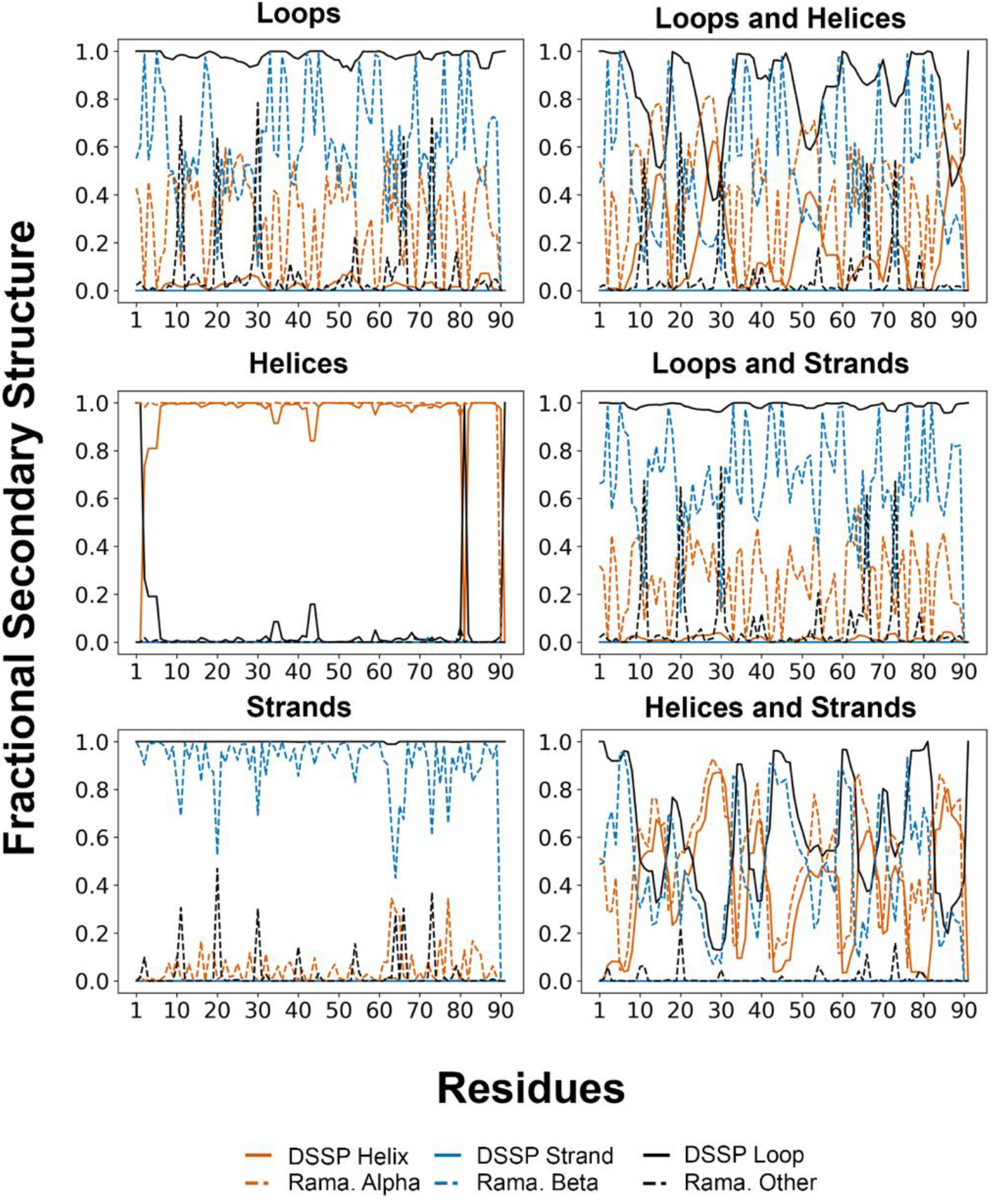
Fractional secondary structure in Sic1 ensembles. Analyses were performed on 1200-conformer pools of Sic1 generated with different combinations of secondary structure sampling consisting of loops, helix, and extended strands. Orange indicates α-helix detected by DSSP (solid) and the α-region on the Ramachandran (Rama.) diagram (dashed). Blue indicates extended strand for DSSP (solid) and β-region on the Ramachandran diagram (dashed). Black indicates coil/loop for DSSP (solid) and other regions on the Ramachandran diagram (dashed).

Sampling with all three secondary structure options in combination (loop, helix, strand) is not the same as sampling with the ANY flag (‘--dany’), as the latter samples based solely on the sequence matching patterns disregarding secondary structure annotations, thus reflecting the inherent structure propensities of the input sequence fragments as present in the database. The explicit listing of secondary structure codes limits to sampling fragments with the same secondary structure code for all residues, while with the ANY flag this is not a requirement, so we wondered if there would be significant differences in emerging structural patterns. Analysis of ensembles of the drkN SH3 domain unfolded state demonstrates no significant visual differences in torsion angle distributions, but the ANY pools do have greater psi ranges between residues 22 and 29 and there are greater helical propensities for the LHE pool compared to the ANY pool (**Figure 9**). Although the general trends of secondary structure propensities seem similar between the LHE and ANY pools, small differences demonstrate that these options generate different conformer pools. We recommend the ANY flag to build IDP conformer ensembles with sampling of torsion angles based on frequencies observed in the PDB. To maximize sampling of torsion angle space, we recommend sampling with both ANY and LHE to minimize torsion angle bottlenecks.

**Figure 9.**
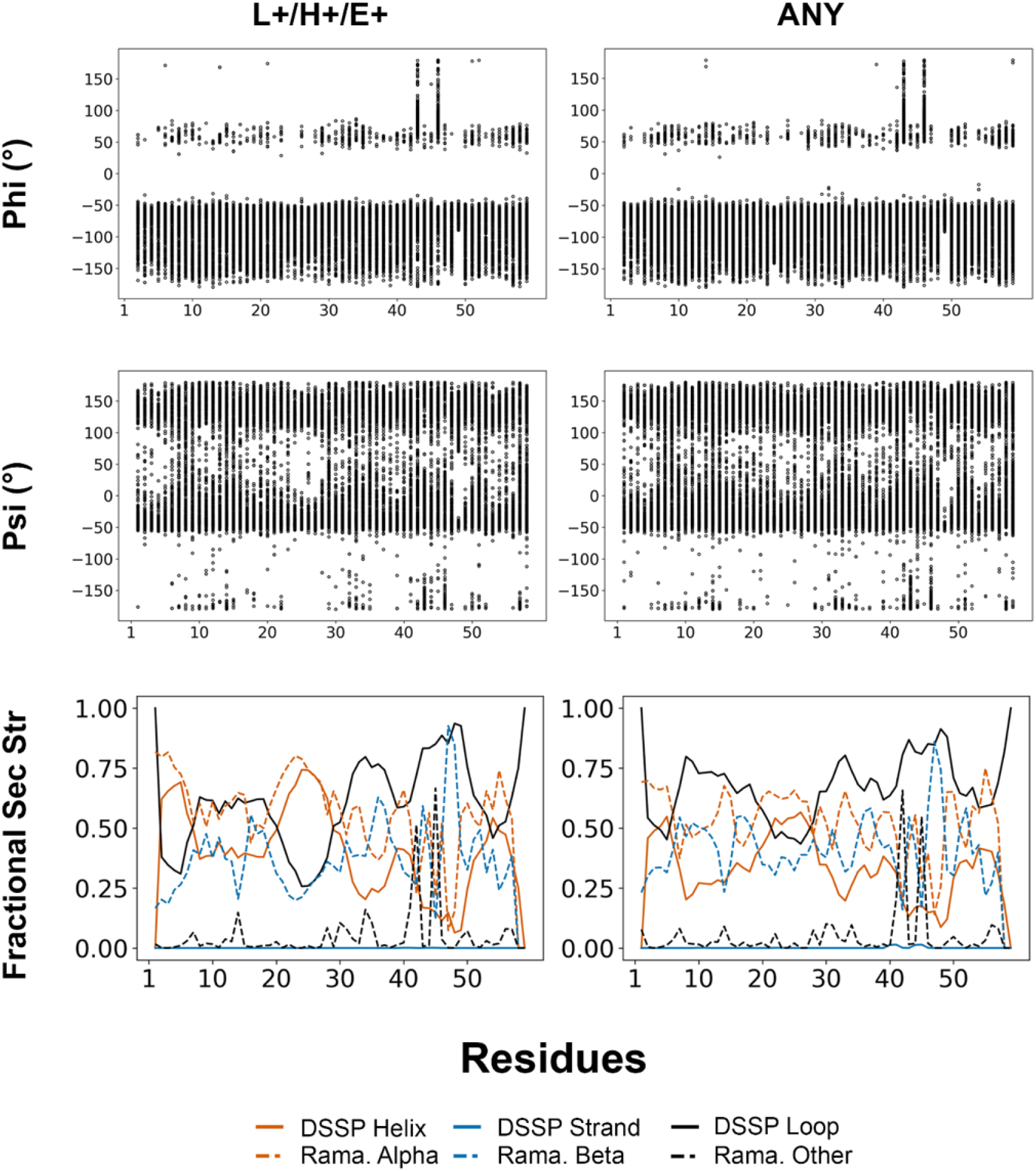
Comparison of torsion angle sampling for L+/H+/E+ and ANY. Ensembles of 1000 conformers each of drkN SH3 domain unfolded state were generated with sampling a combination of loops (L), helix (H), and extended strands (E) or sampling without biasing secondary structure with the ANY flag. Phi and psi (φ and ψ) torsion angle distributions for each conformer pool are shown as a scatter plot in the first two rows. The third row depicts fractional secondary structure based on DSSP (dark solid lines) or the Ramachandran (Rama.) diagram (dashed lines), with orange indicating α-helix for DSSP and α-region of the Ramachandran diagram, blue indicating extended strand for DSSP and β-region of the Ramachandran diagram, and black indicating coil/loop for DSSP and other regions of the Ramachandran diagram.

Importantly, we were interested in whether our design of IDPConformerGenerator to exploit the secondary structural propensities found in the PDB would match experimentally measured propensities from NMR chemical shifts. Two sets of 3000 conformers each of the drkN SH3 domain unfolded state were generated using a backbone energy threshold = 100 kJ, with the “ANY” flag and with the CSSS flag to do custom secondary structure sampling based on δ2D^38^ calculations from NMR chemical shift data^71^ on a per-residue basis. As seen in **Figure 10** (left), there are natural secondary structure propensities for α-helix particularly for residues 16-29) based on δ2D predictions for secondary structure propensities^38^. At residues 58 and 59, the predicted probabilities of secondary structure are set to 1/3 as no chemical shift data are available. Although extended β-strand regions have also been predicted with δ2D, DSSP defines extended strands based on both torsion angle ranges (the same ones as used for segmenting the Ramachandran space) and hydrogen bonds, and there are minimal cases of tertiary contacts satisfying β hydrogen bonds in these disordered ensembles. However, sampling in the β-region of the Ramachandran diagram is plentiful. (Note that it may be valuable to utilize a different definition of strand pairing besides DSSP that is more permissive for local backbone geometries to characterize potential β-strands.) This ensemble shows that helical structure is oversampled relative to what is found experimentally. However, the regions where significant α-helical secondary structure is sampled does overlap the observed secondary structure propensities identified by δ2D. With custom secondary structure sampling, in contrast, there is very good agreement for the α-helical and coil/loop structure on a per-residue basis to that suggested by δ2D (**Figure 10**, left). Sampling in the β-region is consistent, with no observed β-strand H-bonding structure seen using DSSP. Overall, biasing the sampling for torsion angles in the PDB based on secondary structure yields an ensemble with an overestimate of helical structure compared to the δ2D estimates, while directing the sampling by NMR data, as expected, yields an ensemble in much closer agreement to these data.

**Figure 10.**
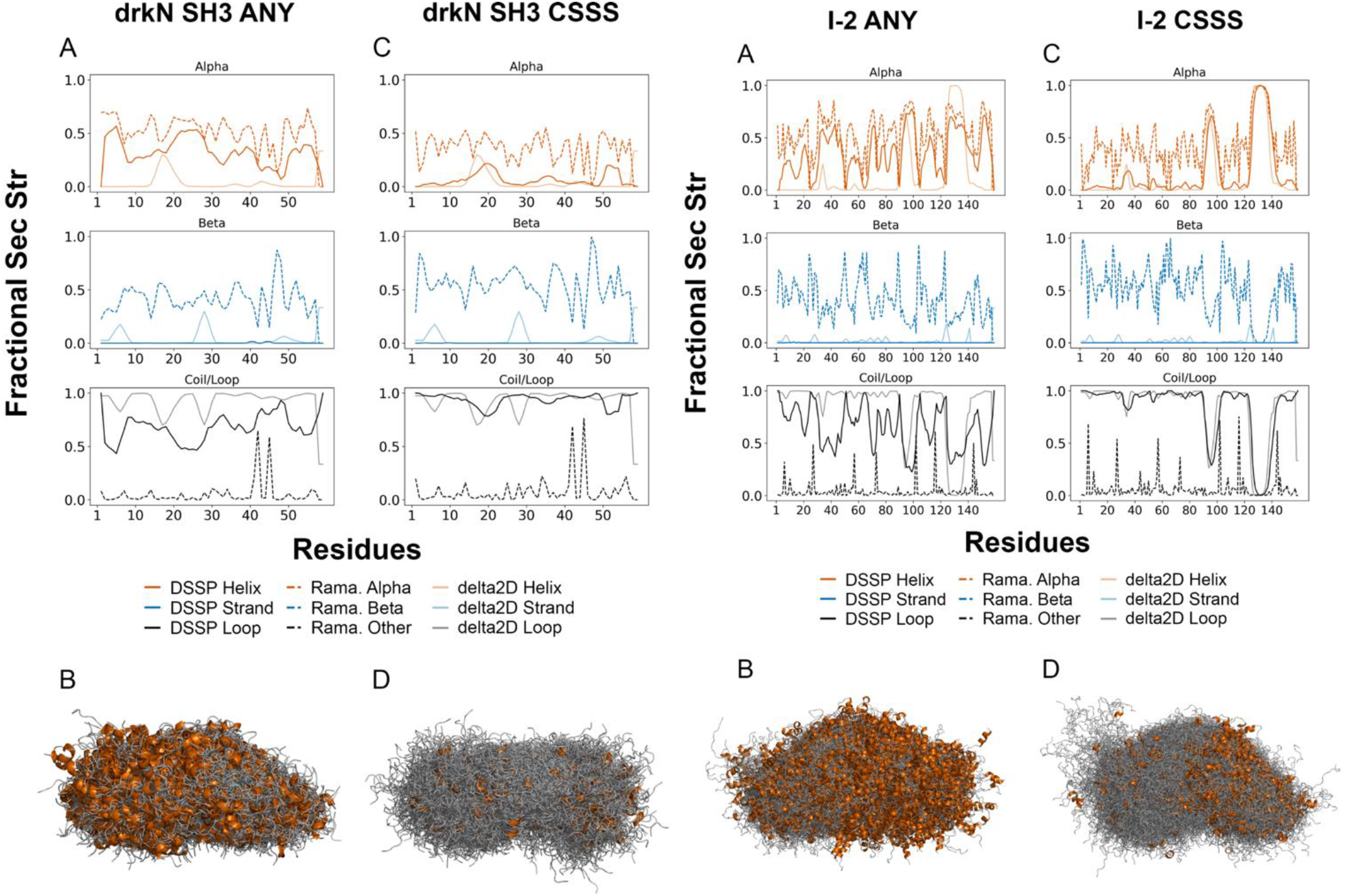
Custom secondary structure sampling. **(left)** For the drkN SH3 domain unfolded state, two sets of 3000 conformers each were generated and **(right)** for inhibitor-2 (I-2), two sets of 1500 conformers each were generated, with **(A, B)** the “ANY” flag or with **(C, D)** the CSSS flag to do custom secondary structure sampling based on δ2D calculations from NMR chemical shift data. **(A, C)** Plots of fractional secondary structure based on DSSP (dark solid lines), the Ramachandran (Rama.) diagram (dashed lines) or δ2D (light solid lines). Orange indicates α-helix for δ2D and DSSP, and α-region on the Ramachandran diagram. Blue indicates extended strand for δ2D and DSSP, and β-region on the Ramachandran diagram. Black indicates coil/loop for δ2D and DSSP, and other regions on the Ramachandran diagram. **(B, D)** Aligned conformers of the ensembles using PyMOL.

Similar results were found for I-2, which has significantly populated helices around residues 85-99 and 121-145, based on NMR chemical shifts^71^ with δ2D assignments. These peaks match with sampling from the α-region of the Ramachandran plot in the ANY ensemble (**Figure 10**, right) but there is significant helical structure throughout. Biasing by the NMR data, we can generate an ensemble with near-exact agreements of the secondary structure propensities calculated by δ2D to the secondary structure propensities of the conformer ensemble calculated by DSSP (**Figure 10**, right). While sampling torsion angles in the PDB using the ANY flag may provide some insight to the natural propensities for α or β regions of the Ramachandran diagram, biasing the sampling based on experimental NMR data can yield conformer pools that are more likely to be representative of the disordered protein. Additional plots of residue-specific fractional secondary structures for calculated ensembles are provided in **Supplementary Figures S9-S12.**

### Comparison with experimental data

Beyond the chemical shift-derived secondary structure, we were interested in the ability of the generated ensembles to match experimental data. While the goal is to build diverse conformer pools that have the potential to fit experiment following a sub-setting or re-weighting procedure, such as with ENSEMBLE^20^ or X-EISD^22,23^, an initial match to experiment clearly demonstrates this potential. Using RMSD from experimental data (**Figure 11 and Supplementary Figure S13**) and ENSEMBLE and X-EISD scores as metrics (**Supplementary Table S4**), we found that IDPConformerGenerator ensembles are not in close agreement with the experimental data, as expected, but that the deviation is not large for many restraints types, such as SAXS, chemical shifts, 3-bond ^1^HN-^1^Ha J-couplings and RDCs. While there are larger deviations for those representing tertiary contacts, PREs and ^1^H-^1^H NOEs, it is difficult to directly compare the RMSD values for the different experimental data types as the range of data varies considerably. An RMSD of ∼4Å for PRE, a measurement that goes out to 20 or 30 Å, may be closer than an RMSD of ∼0.8 ppm Cα chemical shifts, a measurement that varies less than 2 ppm. CSSS generally provides ensembles in better agreement with Cα and Cβ chemical shift restraints, as expected, particularly for proteins with known significant sampling of secondary structure, such as I-2.

**Figure 11.**
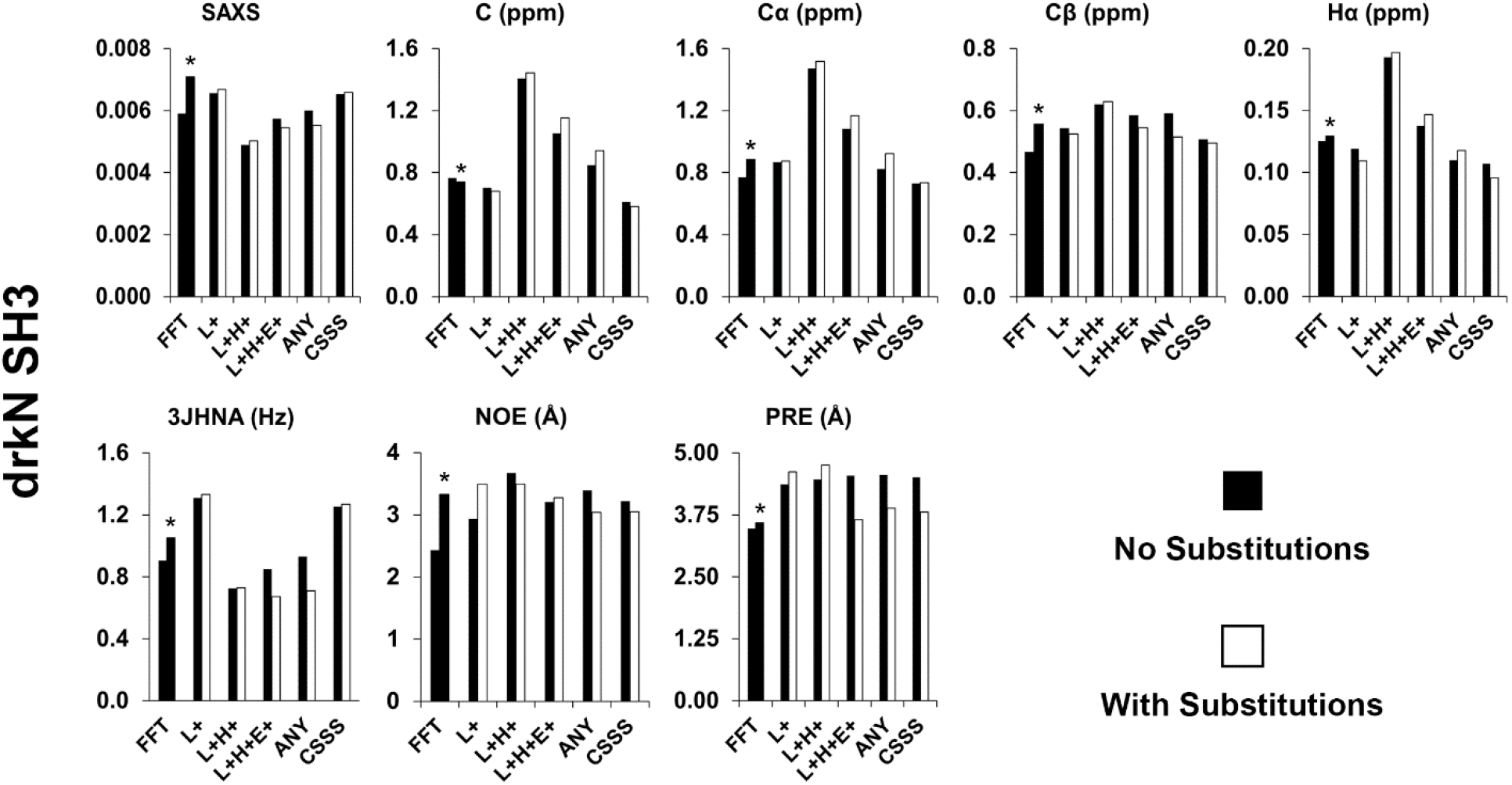
Root-mean squared deviations (RMSDs) of back-calculated values from conformational ensembles to experimental data for the drkN SH3 domain unfolded state. Analyses of 1000-conformer ensembles generated using various secondary structure sampling and using FastFloppyTail (FFT). RMSDs are given for SAXS, chemical shifts (Carbonyl, Cα, Cβ, Hα), PRE, ^3^J_HN-HA_, and NOE if available. Sources of experimental data are provided in Methods. * is for the standard protocol which for this case treats the protein as a mixture of ordered and disordered, while the other is for a modified protocol in which the protein is considered to be fully disordered.

A significant measure of the ability to match experimental restraints is effective sampling of various tertiary contacts. Comparison of the Cα - Cα distance matrices for fragment sizes of 1, 3, and 5, as well as the default fragment size sampling, for Sic1 show that there is a relatively smooth sampling of longer distances (**Figure 12**, top row). Significant differences are observed between ensembles generated with the three different fragment sizes, as seen in the difference distance matrices between ensembles, demonstrating that mixtures of fragment sizes are valuable for sampling a diverse set of tertiary contacts (**Figure 12**, bottom row). In addition, there are significant differences in tertiary contacts for Sic1 ensembles generated with loops and with “ANY” (**Supplementary Figure S14**). Together, these results provide evidence that using a large combined input pool of conformations created with varying fragment sizes, secondary structure sampling and other parameters would enable re-weighting or subsetting to fit distance restraint and other data types.

**Figure 12.**
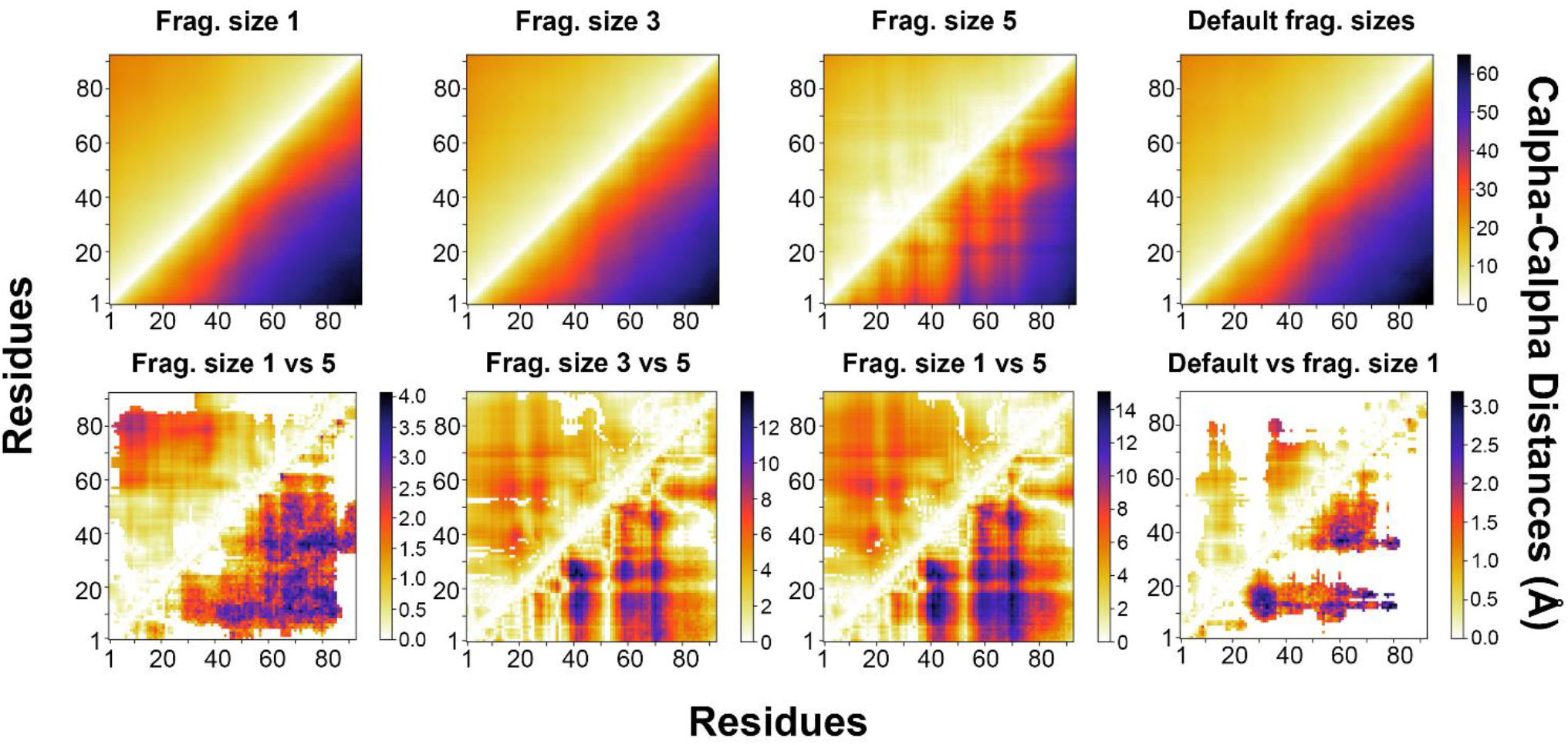
Analysis of tertiary contacts for Sic1 ensembles. (top row) Cα - Cα distance matrixes (lower) with deviations (upper) for 1000-conformer ensembles of Sic1 generated with the loops only flag for secondary structure, with substitutions from columns 5, 3, and 2 of the EDSSMat50 amino acid substitution matrix and with variable fragment lengths. (bottom row) Significant differences between Cα - Cα distance matrixes (lower) and deviations (upper), (P < 0.05 from a Mann-Whitney U test).

### Comparison to other disordered chain generating tools

One of the early motivations for building IDPConformerGenerator is the significant number of steric clashes found in TraDES^12,13^ conformers. The definition of a “clash” depends on whether only the repulsive portion of the LJ potential is considered or the whole LJ potential is used, allowing closer distances which are compensated by favorable interactions. We generated a set of 1000 Sic1 conformers using default TraDES parameters and used Chimera to check for steric clashes^72^ using a stringent criterion based largely on distance (similar to the repulsive portion of the L-J). With criteria of no backbone clashes and <=5 sidechain clashes, this TraDES ensemble had no conformers meeting the criteria. In contrast, 1000-conformer IDPConformerGenerator ensembles of Sic1 built using custom secondary structure sampling and default fragment sizes had 324/1000 conformers meeting these criteria. A similar drkN SH3 domain unfolded state pool had 395/1000 conformers meeting these criteria. Chimera’s clash definition is more stringent than the one we utilize in IDPConformerGenerator and MC-SCE, which allows close contacts if compensated by favorable L-J energy, thus all these IDPConformerGenerator conformers are arguably physically realistic conformations. IPDConformerGenerator does generate many more conformers with fewer steric clashes as defined by Chimera than does TraDES.

We also calculated a set of FastFloppyTail^15^ ensembles to enable comparison with IDPConformerGenerator. While there are different parameters for running FastFloppyTail, it is not user customizable in terms of sampling specific secondary structures or various fragment lengths, with the exception of using 3-mer and/or 9-mer fragment libraries. We therefore used the recommended protocol for each system, with 3-mer fragment libraries, and with an additional run to force the drkN SH3 domain to be disordered throughout (see Supplementary Information for details). We measured the speed, diversity, sampling of secondary structure and match to experimental data (**Figures 5, 11, Supplementary Figures S6, S9-S13, and Supplementary Tables S2, S3, S4**). For the quantitative speed comparison, we only considered the time of the calculation following set up with the initial files. For IDPConformerGenerator, initial set up includes the generation of the initial torsion angle database, which we only needed to do once for all the systems, and providing the specific protein sequence with input parameter files. Generating the initial torsion angle database took 37 minutes on a desktop computer using 63 of the 64 cores, 20-30 minutes for downloading, while processing and generating the database were fast. The time is very dependent on the internet connection speed and number of cores since the process is embarrassingly parallel. For FastFloppyTail, there is no ability to define multiple processors, and there are a number of steps and files required before the conformer generation process, including the prediction of disordered regions and secondary structure and creating a fragment library, which is required for each protein. The predictions require multiple external websites or programs. For the test systems, we could utilize the premade files on the FastFloppyTail website but for other proteins this would not be the case. For the drkN SH3 domain, Sic1, aSyn and I-2, the fragment libraries took between 11 and 16 minutes each to calculate on the HPC system we used for ensemble generation, while for Tau it was about 45 minutes (**Supplementary Table S2**). Another issue with FastFloppyTail is that disordered proteins that are not predicted to be disordered can yield challenges in setup. In particular, the unfolded state of the drkN SH3 domain has a sequence that is not predicted to be disordered, leading to our testing both the recommended algorithm and one defining it as disordered (see Supplementary Information). Alpha-synuclein is known to fold into a long α-helix in the presence of lipid or micelles and different predictive algorithms have variable success in predicting its disordered state^73^, and the authors of FastFloppyTail noted a need to find a predictor which correctly identified its disordered state^15^.

The results demonstrate that IDPConformerGenerator is faster than FastFloppyTail for chains shorter than 200 residues with default parameters. There is generally minimal difference in the diversity of the ensembles between the two tools, although asphericity values are higher for IDPConformerGenerator and FastFloppyTail ensembles are often more compact. The secondary structure sampling for FastFloppyTail ensembles often falls between IDPConformerGenerator ANY and CSSS biased by NMR chemical shifts, with FastFloppyTail having higher populated secondary structure than suggested by experiment. Matches to experimental data are more variable, with IDPConformerGenerator run using different secondary structural sampling approaches giving lowest RMSD values for different data types for different systems, often lower than the FastFloppyTail ensemble, particularly for the CSSS with chemical shifts, although not always. A clear distinction is that IDPConformerGenerator enables users to flexibly define different approaches for generation of conformers, including for diversifying the resulting ensembles.

## DISCUSSION AND CONCLUSIONS

A range of theoretical and computational approaches for generating disordered ensembles exist^10,74^, each of which has strengths and unique features based on the design philosophy. Testing of IDPConformerGenerator on our set of model disordered proteins demonstrates that this tool is extremely flexible and can function as a platform to enable generating various initial conformational pools built with different biases and parameters, valuable for addressing a range of scientific needs. It is computationally efficient depending on sequence length and parameters, and can enable sampling using the frequencies of secondary structures within the PDB database provided or the experimental secondary structural propensities from NMR experiments. The resulting ensembles are not far from fitting experimental data, including those for local structure, tertiary contacts and overall hydrodynamic properties. Future work will explore the optimal parameters for sampling structures to facilitate identification of subsets or re-weighting to best fit these tertiary contact restraints. However, the current results strongly suggest that using an input pool with a combination of ensembles generated with different approaches, including with and without substitutions, varied fragment sizes and combinations, and varied secondary structure sampling including bias with NMR chemical shift-derived probabilities, can effectively sample a range of conformational space to facilitate fitting experimental data with subsets or re-weighting.

Scientific software is often created by scientists and not software engineers, leading to tools that are not as user-friendly, generalizable, easy to maintain or thoroughly documented as desired. A larger goal of developing IDPConformerGenerator was to design it to be easy-to-use so that it would be widely used, not only to generate disordered protein ensembles as starting pools for subsetting or re-weighting but also to enable it to function as a platform for adding existing functionality or future approaches to define ensembles that best fit experimental data, for computational experiments testing various ideas and for analyses of resulting ensembles. There are a number of straight-forward extensions of IDPConformerGenerator planned, including the ability to build disordered regions around a folded domain, an important functionality (and one that motivated the creation of FastFloppyTail). Due to IDPConformerGenerator’s modularity, analysis tools can easily be built utilizing current functions. Currently available functions include those to analyze ensembles for fractional secondary structure and torsion angle distributions and to analyze the database for the number of available sequence matches and for identifying structures with select keywords. Further additions, such as the analysis of tertiary contacts, could be implemented with ease and are planned for a future release. We envision that IDPConformerGenerator will be the basis for an expanding platform of tools to facilitate structural characterization of IDPs and IDRs consistent with solution experimental data. Ultimately, the resulting ensembles should provide physical insights into how these dynamics states regulate and carry out their critical biological functions and how disease variants in IDPs/IDRs lead to pathology.

## Supporting information

Analysis scripts

## SUPPORTING INFORMATION DESCRIPTION

The following files are available free of charge.

Supplementary Information in the file IDPConfGen_Supplementary_Information.pdf (PDF)

Supplementary Tables in the file IDPConfGen_Supplementary_Tables.xlsx (Excel)

Analysis Scripts in the file analysis_scripts_IDPConfGen.zip (zip file with Python .py files)

## ACKNOWLEDGEMENTS

J.D.F.-K. and T.H.-G. acknowledge funding from the National Institute of Health under Grant 5R01GM127627-04. J.D.F.-K. also acknowledges support from the Natural Sciences and Engineering Research Council of Canada (2016-06718) and from the Canada Research Chairs Program. We acknowledge Sean Reichheld and Simon Sharpe for beta testing. We thank P. Andrew Chong and Alan Moses for critical reading of the manuscript.

## ABBREVIATIONS

IDPs: intrinsically disordered proteins
IDRs: intrinsically disordered regions
NMR: nuclear magnetic resonance spectroscopy
SAXS: small-angle X-ray scattering
SMF: single molecule fluorescence
PDB: Protein Data Bank
MC-SCE: Monte Carlo Side Chain Ensemble
smFRET: single molecule fluorescence resonance energy transfer
CSSS: custom secondary structure sampling
Rg: ra-dius of gyration
Ree: end-to-end distance
FFT: FastFloppyTail
RMSD: root-mean-squared deviation
aSyn, alpha-synuclein: I-2, inhibitor-2

---

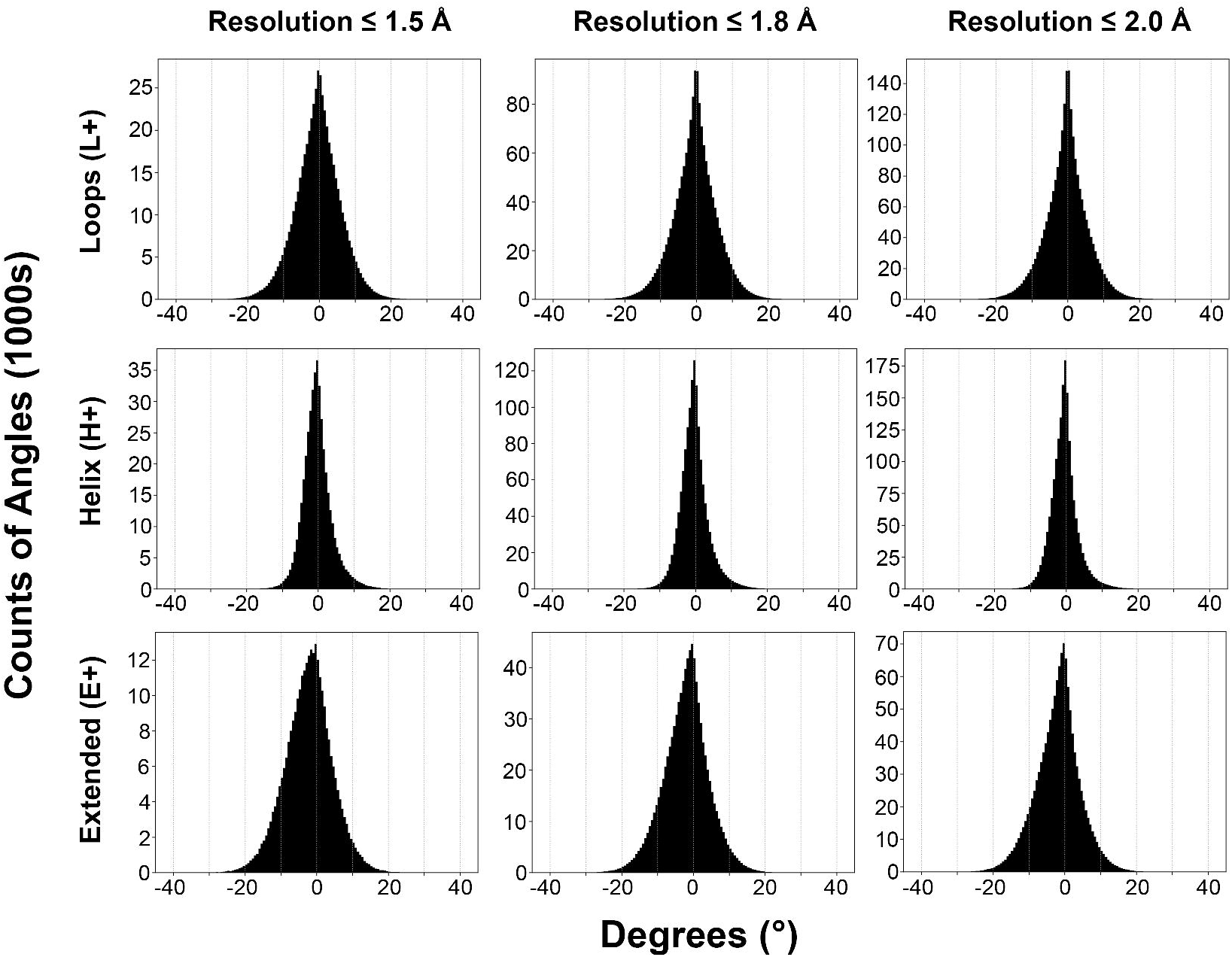

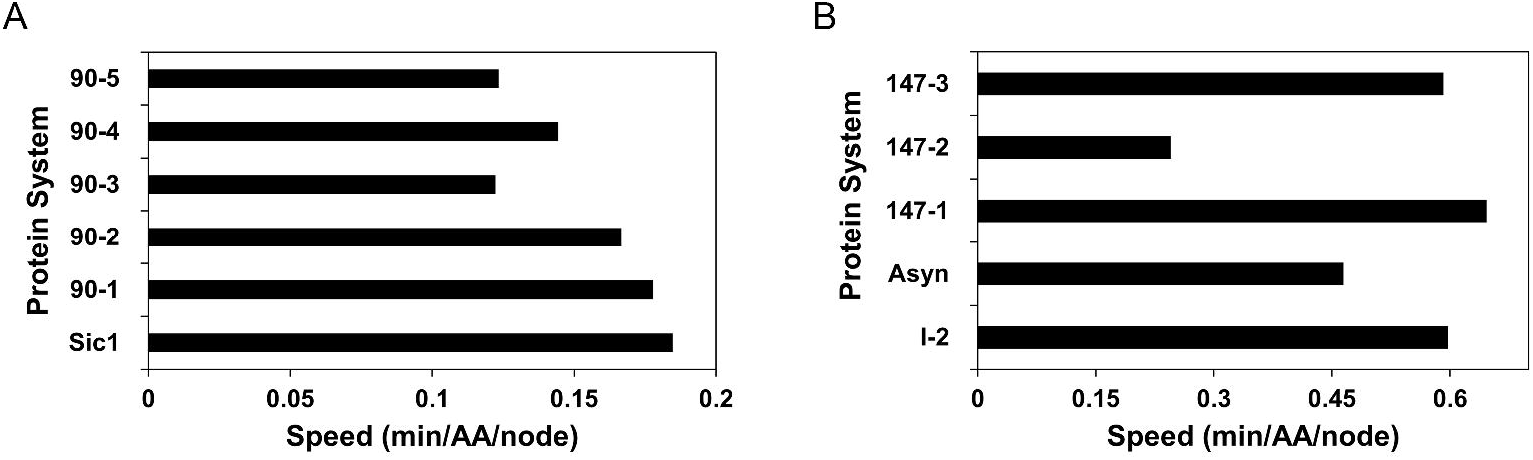

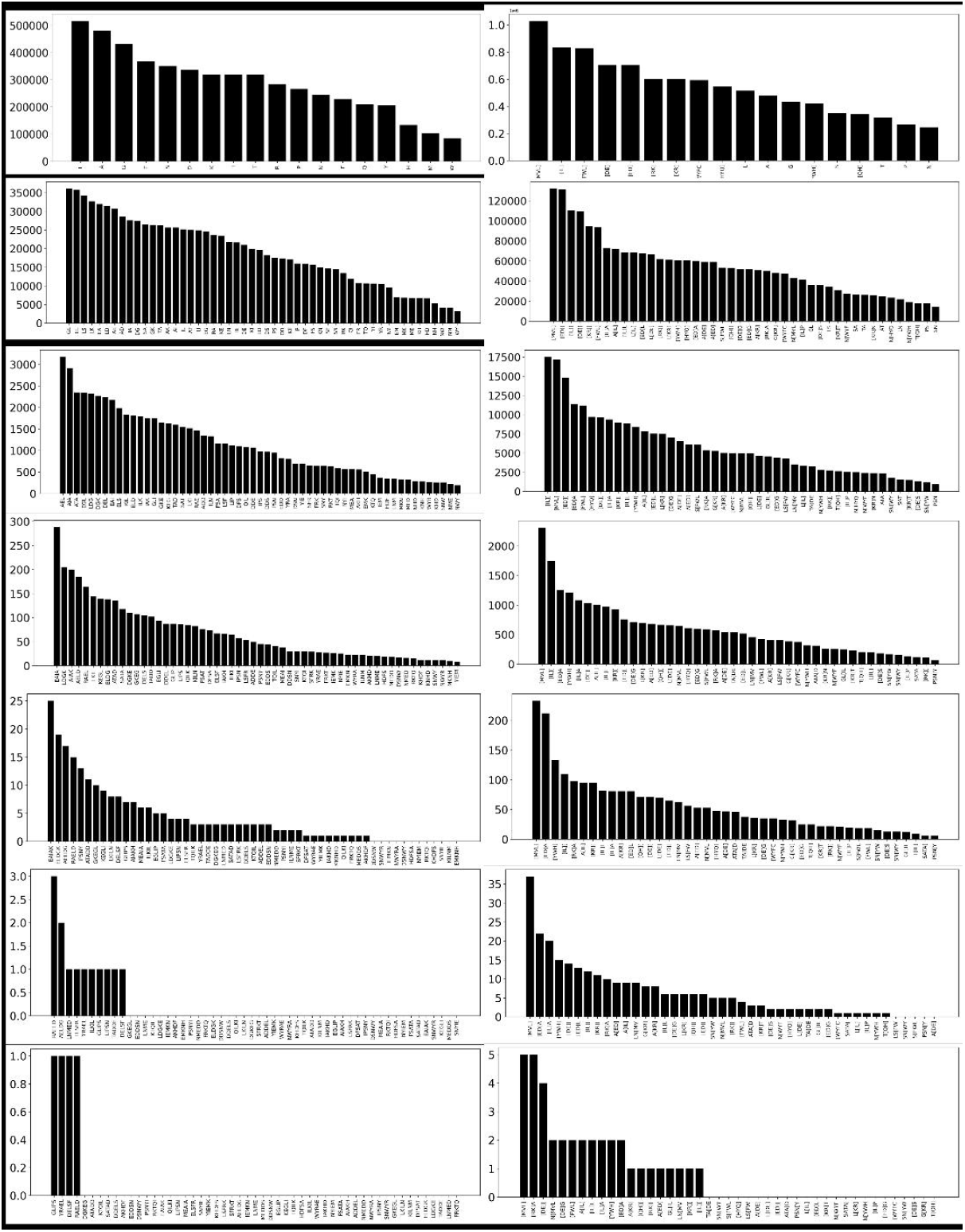

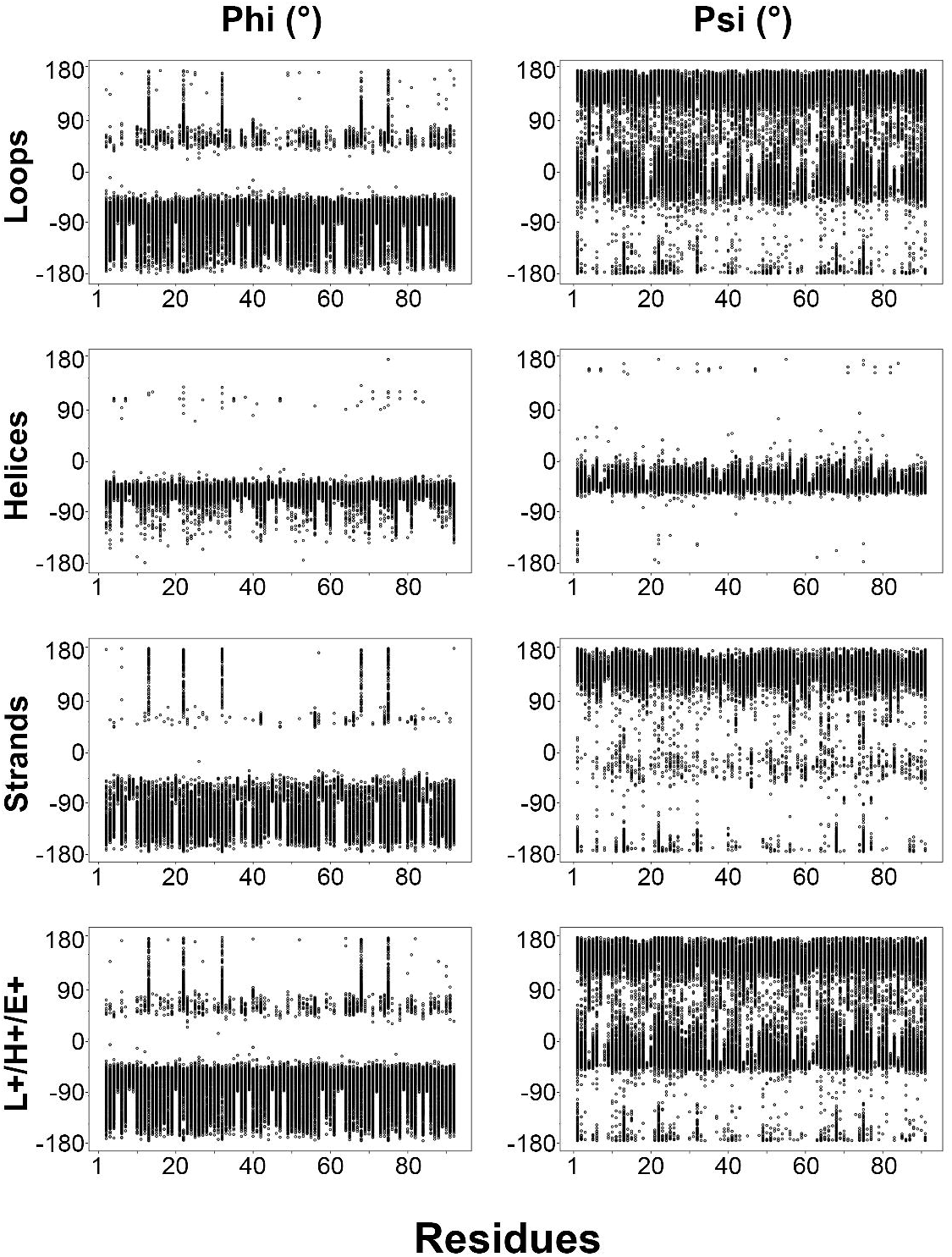

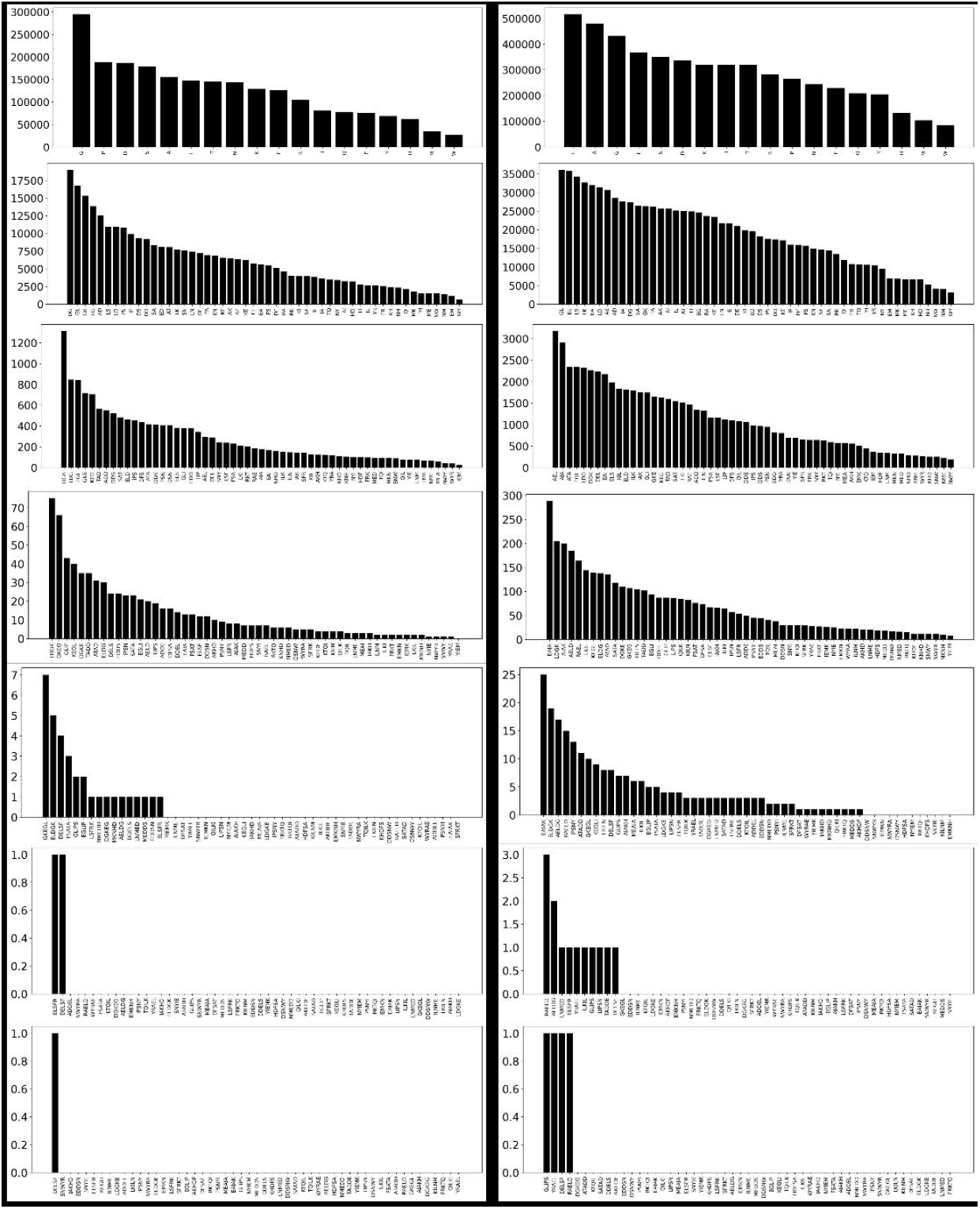

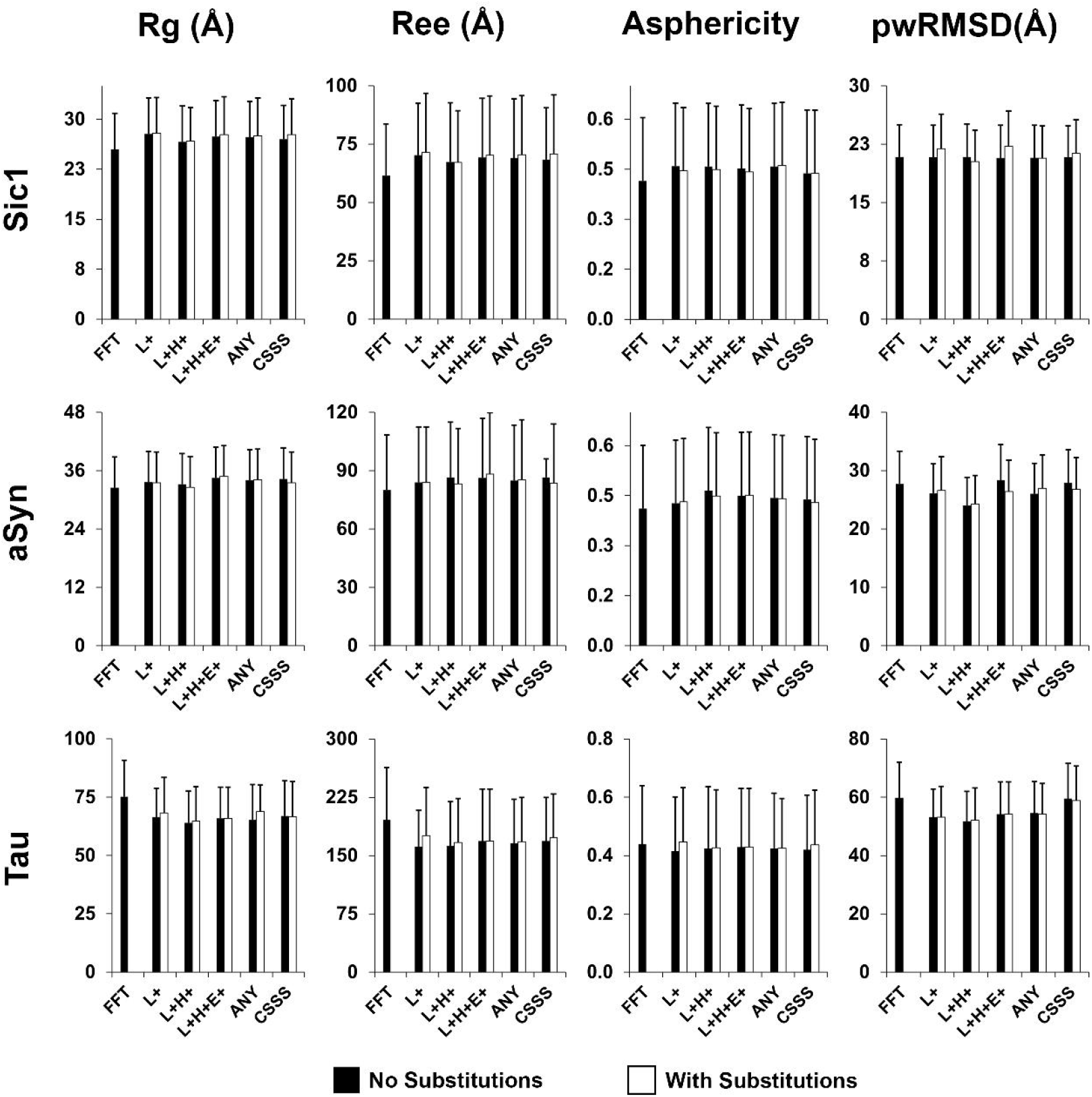

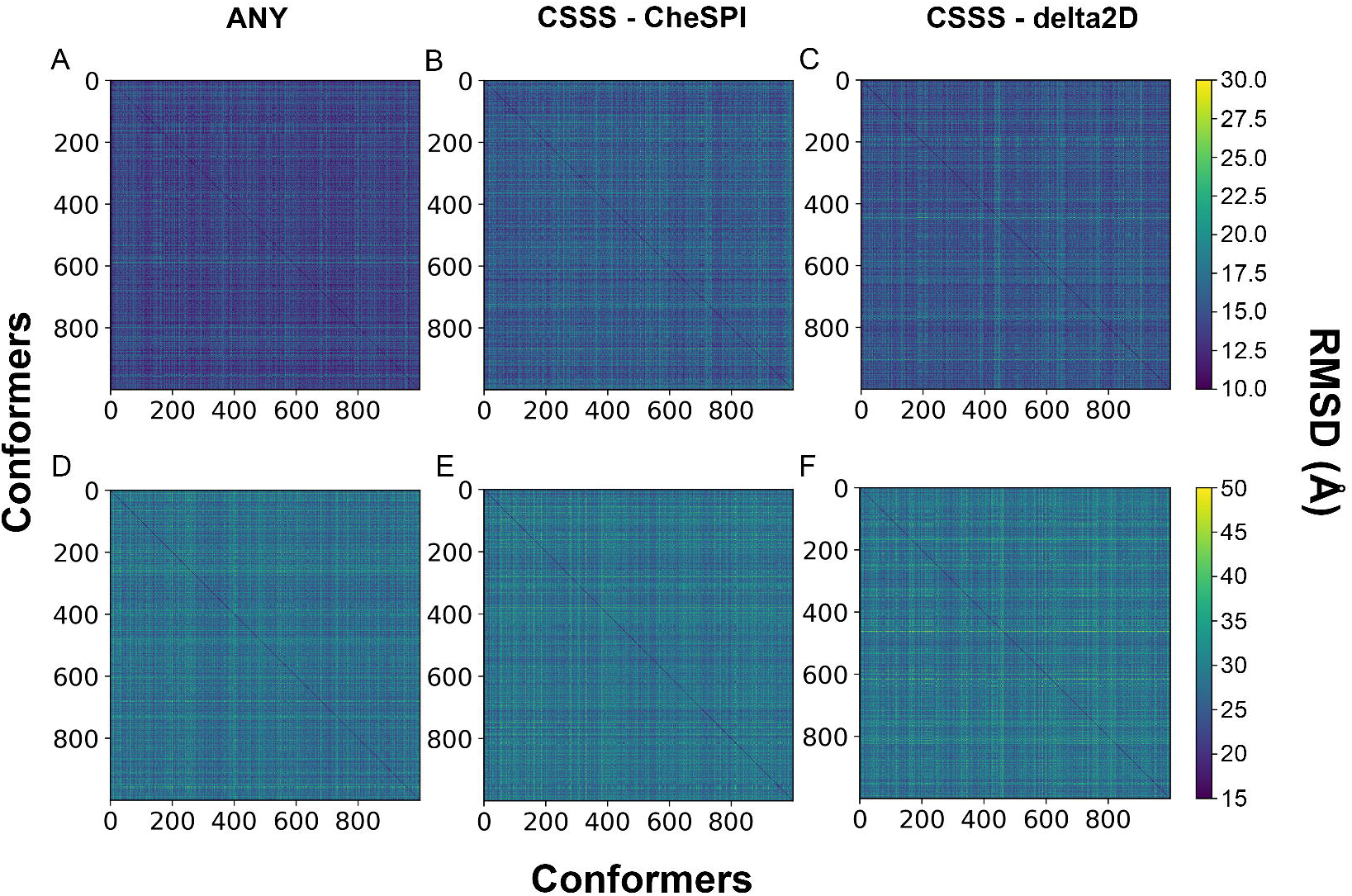

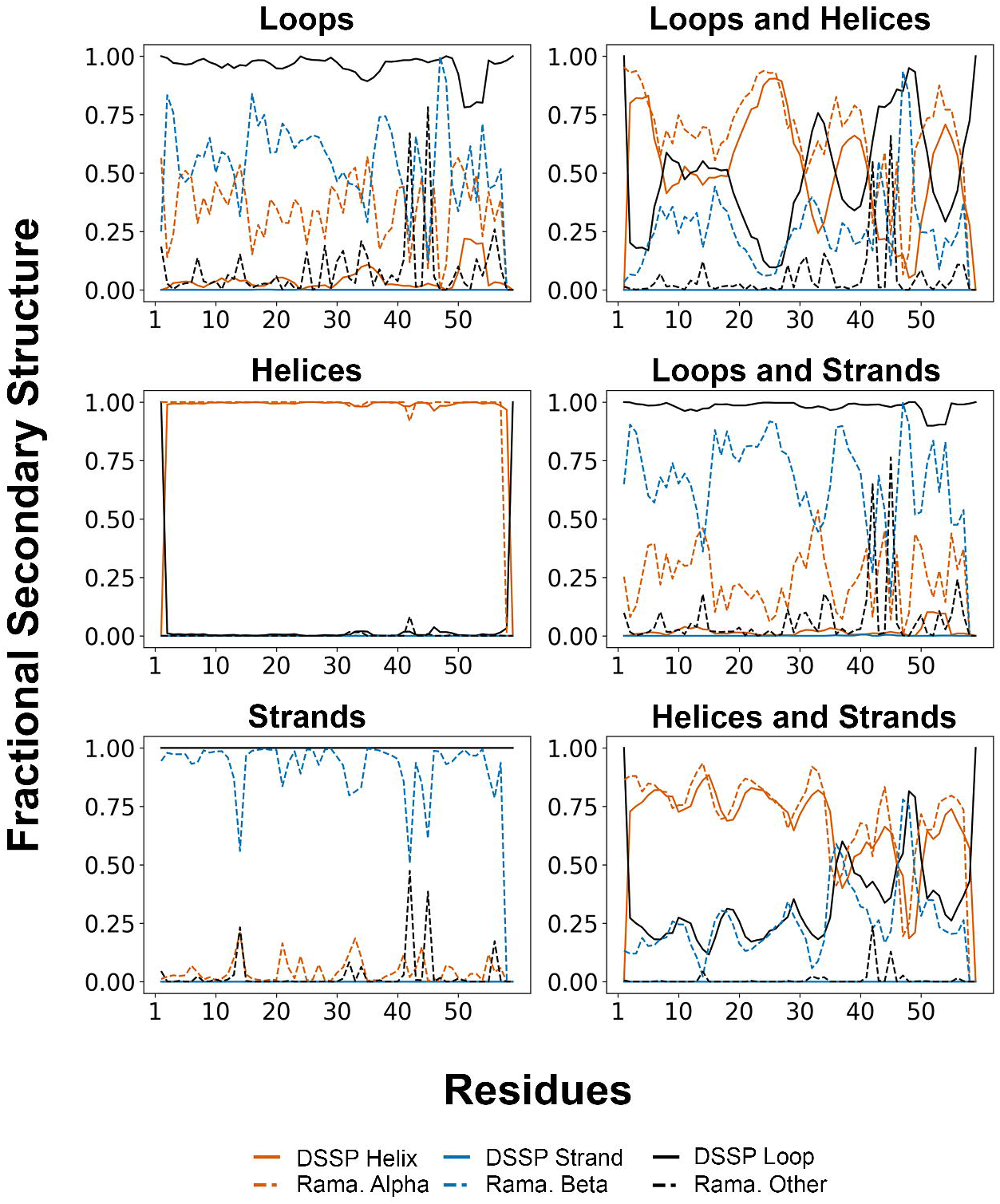

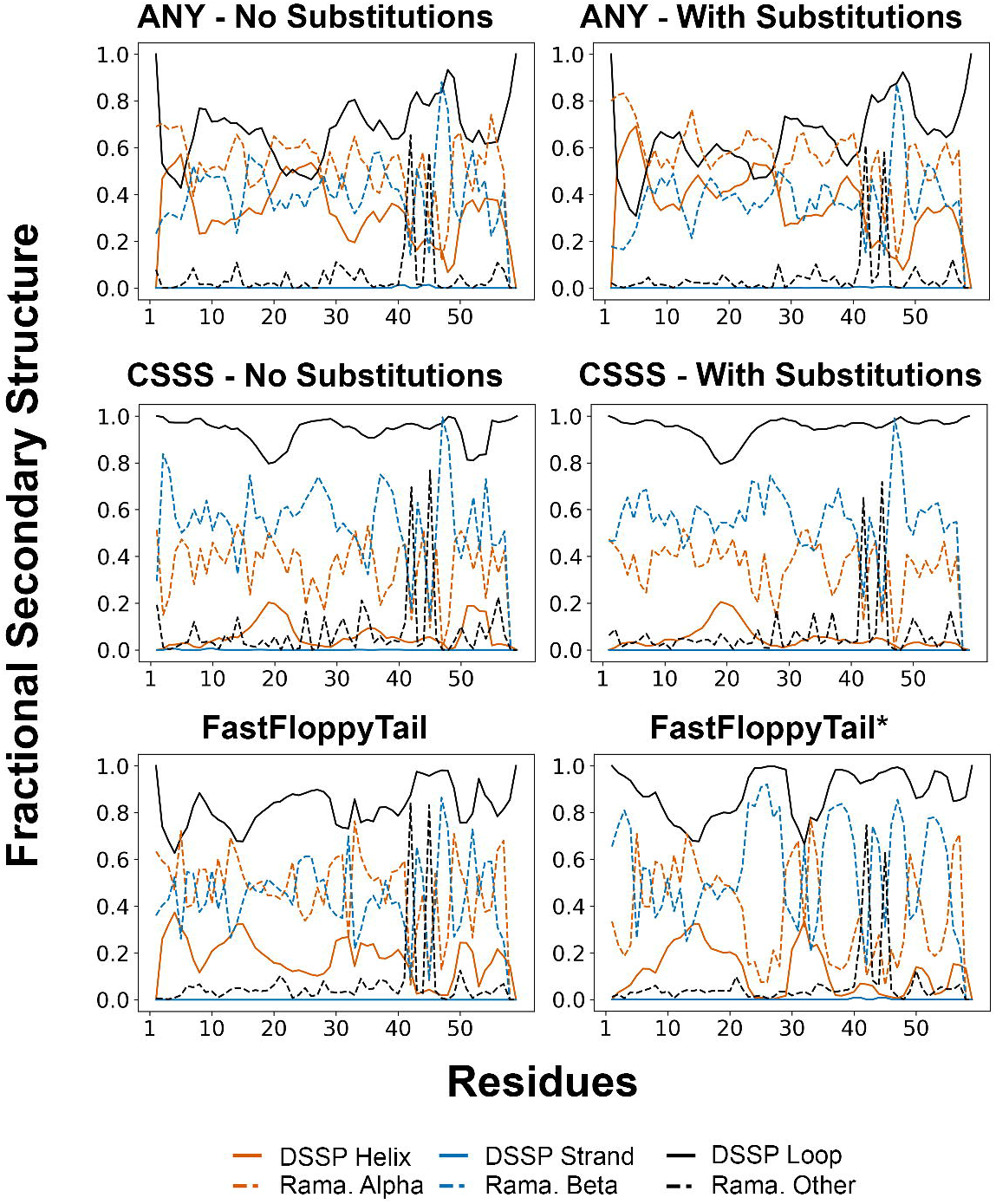

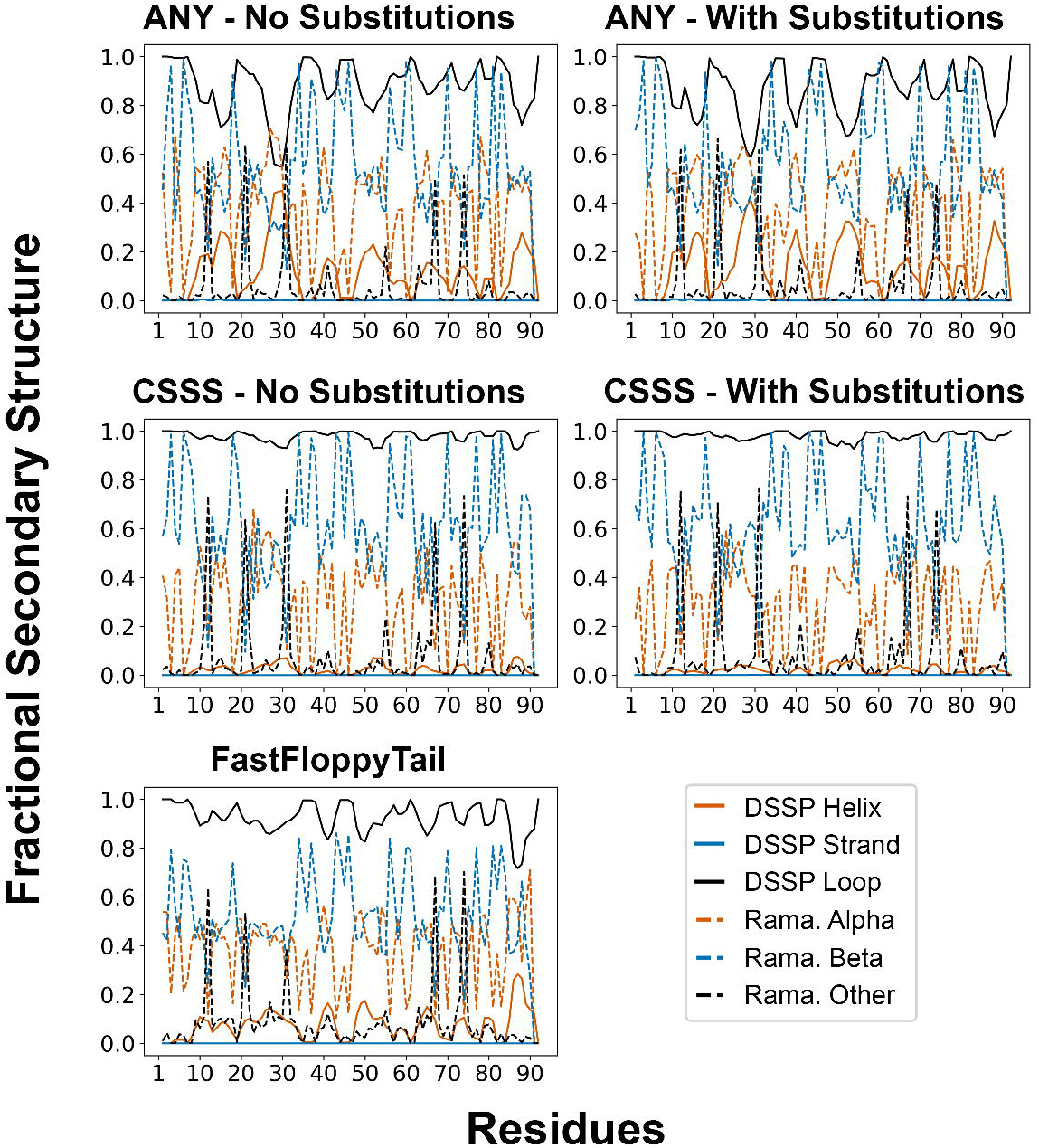

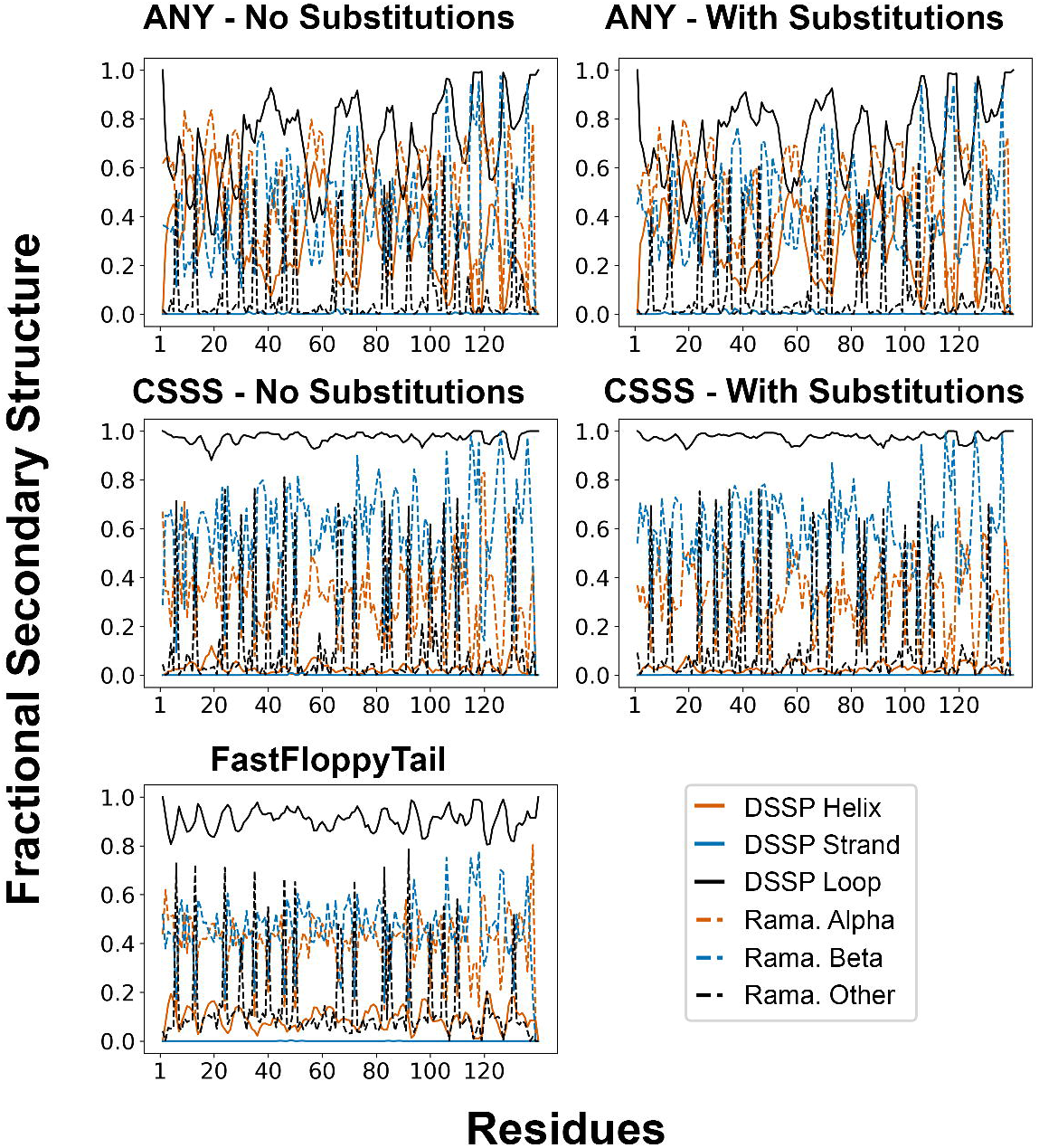

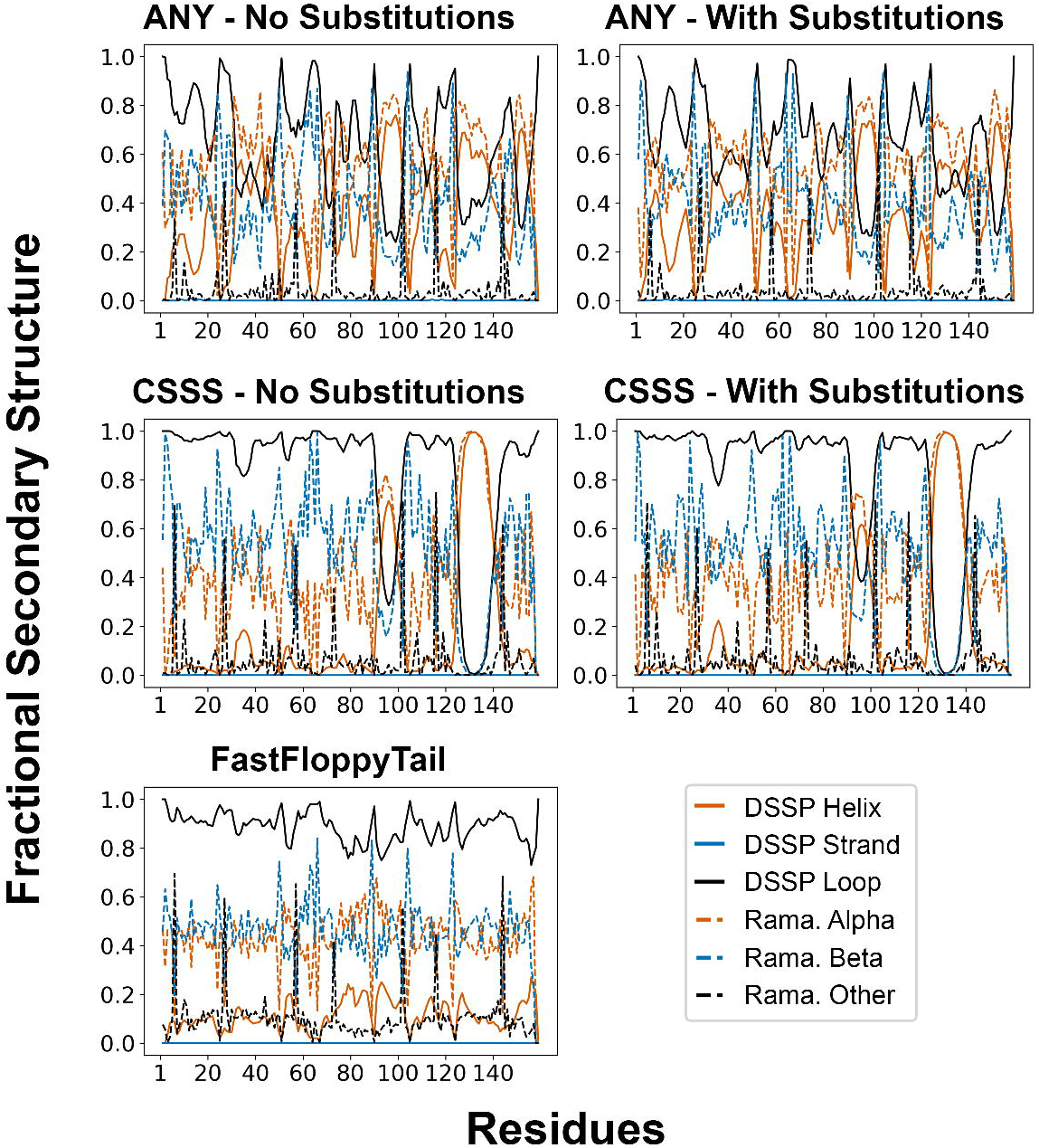

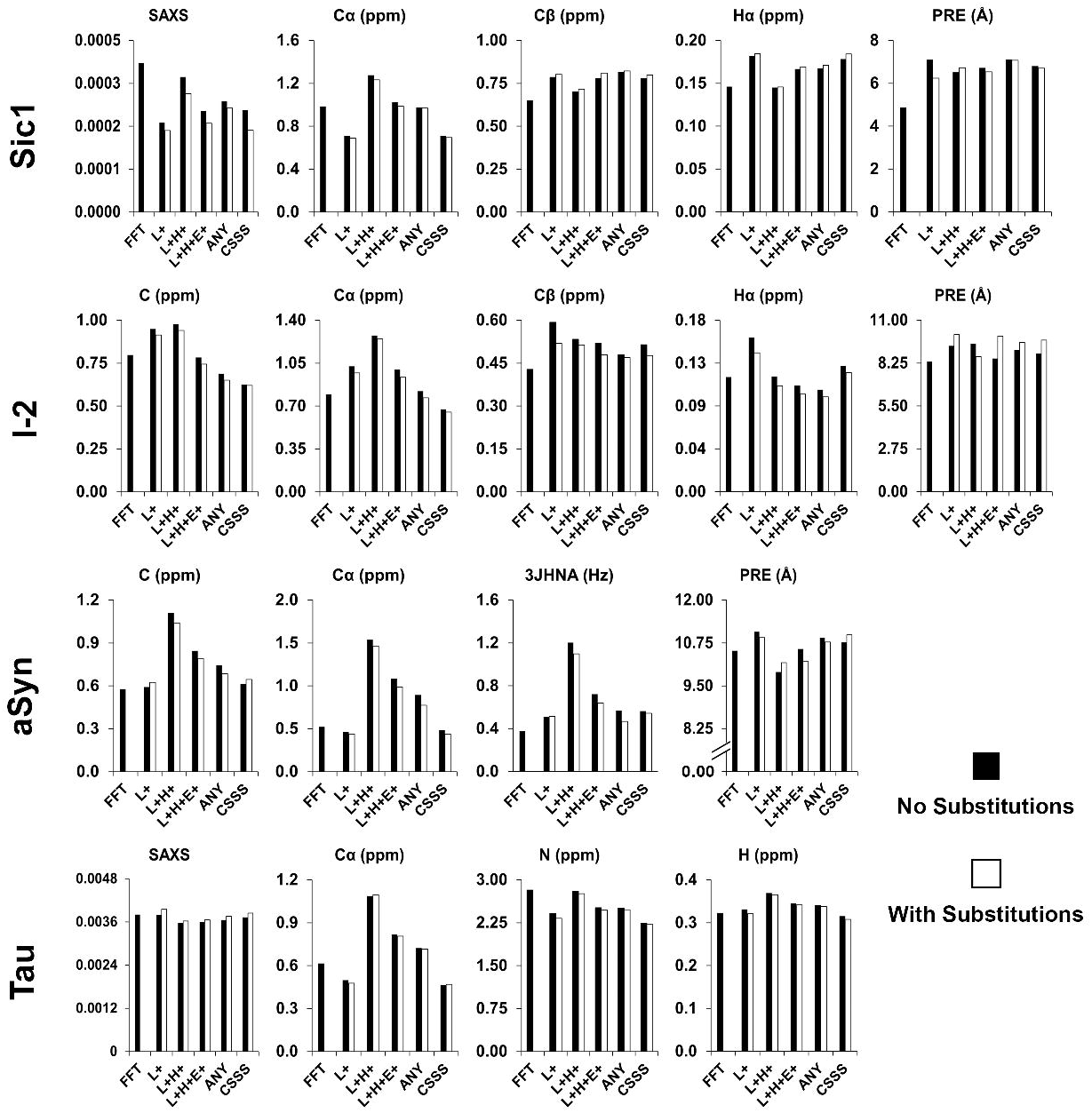

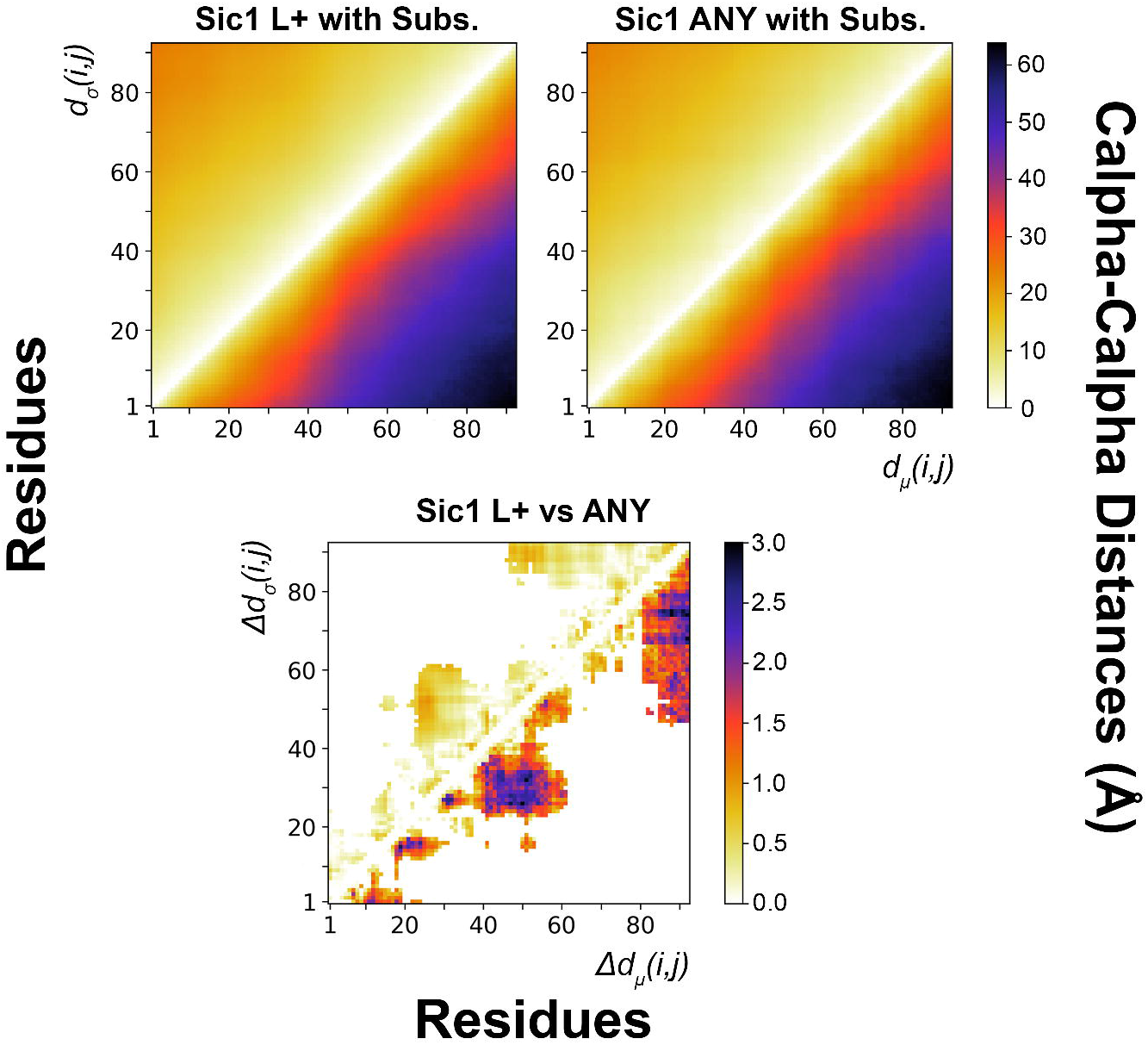

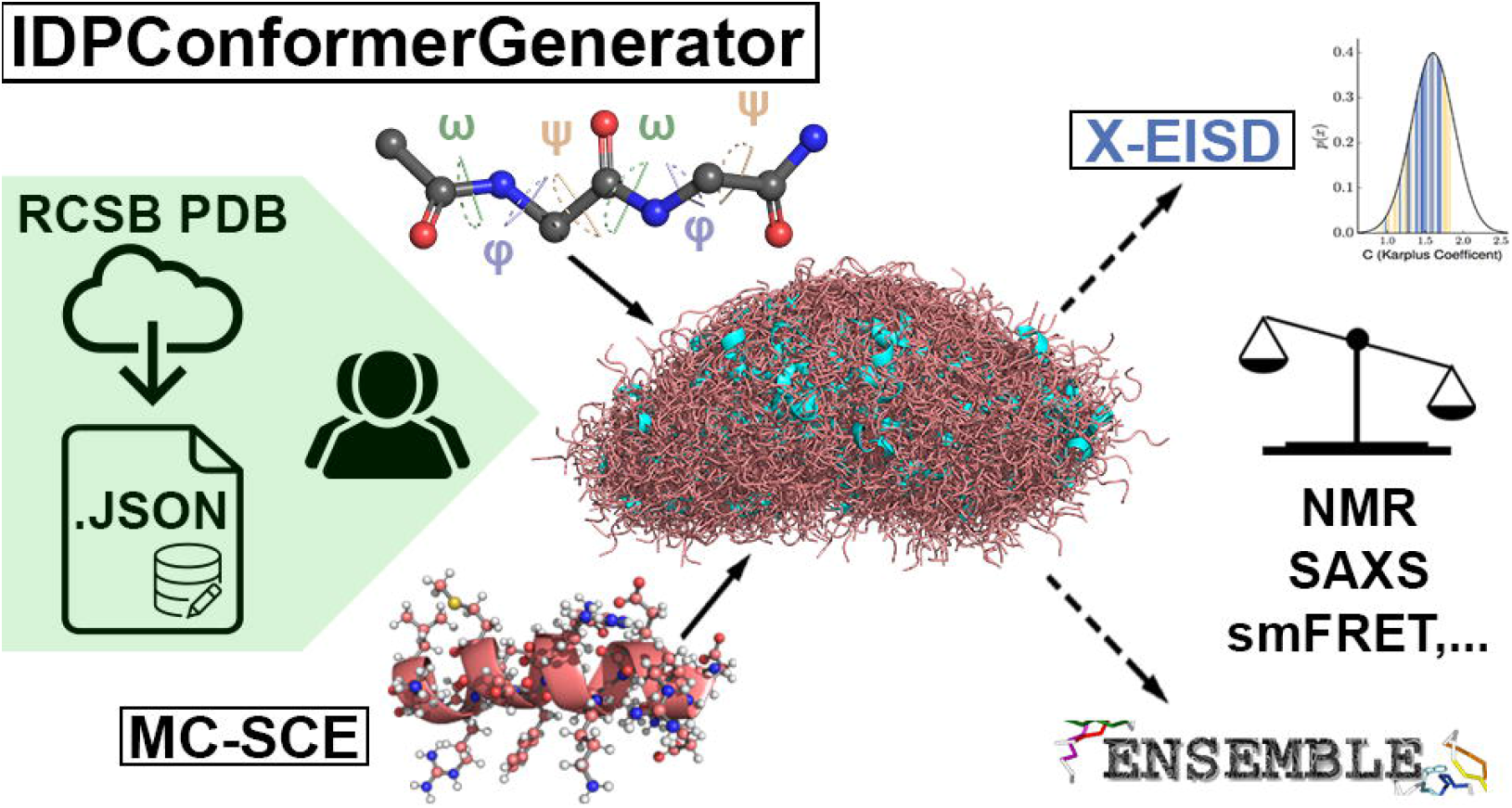

