## Supplementary material for "IDPConformerGenerator: A Flexible Software Suite for Sampling Conformational Space of Disordered Protein States": Analysis scripts: Calculate_Diversity.py

```
#!/usr/bin/env python3
# -*- coding: utf-8 -*-
"""
Created on Sat Jul 24 17:44:38 2021
Script to calculate the radius of gyration, end-end distance, asphericity, and relative standard devations 
and standard errors of the mean of these values for ensembles of PDBs using MDAnalysis 
packages. Script calculates the mean pairwise RMSD and its standard deviation using MDAnalysis packages.

References: 

https://www.mdanalysis.org/MDAnalysisTutorial/trajectories.html
https://userguide.mdanalysis.org/stable/examples/quickstart.html
https://docs.mdanalysis.org/dev/documentation_pages/core/groups.html
https://www.mdanalysis.org/pages/basic_example/
https://userguide.mdanalysis.org/stable/examples/analysis/alignment_and_rms/pairwise_rmsd.html

@author: Ashley Namini
"""
import MDAnalysis as md
from MDAnalysis.analysis import diffusionmap, align 
import glob
import numpy as np 


#path to folder of .pdb files
PATH2PDBs =  '/home/user/Documents/Conformers/asyn/ANY_sub'
files = glob.glob(PATH2PDBs+'/*.pdb')
u = md.Universe(files[0],files)


##### Global Parameters #####

bb = u.select_atoms('backbone')
nterm = u.select_atoms('protein and name N')[0]  
cterm = u.select_atoms('protein and name C')[-1]

rg = [] #list containing the Radius of gyration of each conformation
Ree = [] #list containing the End-end distance of each conformation
A = [] #list containing the asphericity of each conformation

for ts in u.trajectory:
   r = cterm.position - nterm.position
   d = np.linalg.norm(r)
   Ree.append(d)
   rg.append(u.atoms.radius_of_gyration())
   A.append(bb.asphericity())

#convert lists to numpy arrays
rg = np.array(rg)
Ree = np.array(Ree)
A = np.array(A)


mean_Rg = np.mean(rg) #ensemble average of Radius of gyration

mean_A = np.mean(A) #ensemble average of Asphericity

std_Rg = np.std(rg,ddof = 1) #sample standard deviation of Rg distribution

std_Ree = np.std(Ree,ddof = 1) #sample standard deviation of Ree distribution

std_A = np.std(A,ddof = 1) #sample standard deviation of A distribution

sem_Rg = std_Rg/np.sqrt(len(files)) #standard deviation of the mean of Rg distribution

sem_Ree = std_Ree/np.sqrt(len(files)) #standard deviation of the mean of Ree distribution

sem_A = std_A/np.sqrt(len(files)) #standard deviation of the mean of A distribution

rms_Ree = np.sqrt(np.mean(np.square(Ree,Ree))) #Root-mean square average of Ree distribution

rsd_Rg = std_Rg/mean_Rg #relativate standard deviation of Rg distribution

rsd_Ree = std_Ree/rms_Ree #relativate standard deviation of Ree distribution

rsd_A = std_A/mean_A #relativate standard deviation of A distribution


print('rms Ree: {}'.format(rms_Ree))
print('Mean Rg: {}'.format(mean_Rg))
print('Mean A: {}'.format(mean_A))
print('                        ')
print('SEM Ree: {}'.format(sem_Ree))
print('SEM Rg: {}'.format(sem_Rg))
print('SEM A: {}'.format(sem_A))
print('                        ')
print('RSD Ree: {}'.format(rsd_Ree))
print('RSD Rg: {}'.format(rsd_Rg))
print('RSD A: {}'.format(rsd_A))

print('                        ')

###### Pairwise RMSD ###### 

aligner = align.AlignTraj(u, u, select='name CA',
                          in_memory=True).run()

matrix = diffusionmap.DistanceMatrix(u, select='name CA').run()

pairwise_RMSD_matrix = matrix.dist_matrix

upper_triang_RMSD_matrix = np.triu(pairwise_RMSD_matrix)

upper_triang_RMSD_matrix[upper_triang_RMSD_matrix == 0] = np.nan

#this line outputs the mean pairwise RMSD
print('Pairwise RMSD: {}'.format(np.nanmean(upper_triang_RMSD_matrix)))

#this line outputs the sample standard deviation 
print(f'Pairwise RMSD STD: {np.nanstd(upper_triang_RMSD_matrix,ddof = 1)}')
```
