## Supplementary material for "IDPConformerGenerator: A Flexible Software Suite for Sampling Conformational Space of Disordered Protein States": Analysis scripts: Calculate_RMSDs.py

```
#!/usr/bin/env python3
# -*- coding: utf-8 -*-
"""
Created on Wed Apr  6 20:06:35 2022

This script calculates the RMSD for corresponding data types used in ENSEMBLE calculation. For example,
if ran ENSEMBLE using the SAXS and PRE module then this script will output the RMSD for the SAXS and PRE data
for that ENSEMBLE run. 

Requires: 
    1. constraint files are tab delimited or white space delimited
    2. Get_RMSDs.py, cdp.py and Calculate_RMSDS.py are in same folder
    3. Must have used ENSEMBLE using one CSP file and NB_GEN_NEW and NB_GEN_START are 0 in the .par file 

@author: Ashley Namini
"""
import os
from Get_RMSDs import SAXS_RMSD,CS_RMSD,PRE_RMSD,JC_RMSD,RDC_RMSD,NOE_RMSD

#Path to .par file used for ENSEMBLE calculation
ENSEMBLE_PAR_FILE = '/home/user/Documents/Drk_JOB1.par'

#Number of conformers in ensemble
n_conf = 1000

file = open(ENSEMBLE_PAR_FILE,'r')
Lines = file.readlines()

file.close()

bc_data_folder = ''
bc_data_folder_path = ''
csp_file_name = ''

bc_data_PRE = ''
bc_data_SAXS = ''
bc_data_J = ''
bc_data_NOE = ''
bc_data_MRDC = ''
bc_data_SHIFTX = ''

FLAG_PRE = 0
FLAG_J = 0
FLAG_NOE = 0
FLAG_SAXS = 0
FLAG_MRDC = 0
FLAG_SHIFTX = 0

SHIFTX_CON = ''
SAXS_CON = ''
MRDC_CON = ''
NOE_CON = ''
PRE_CON = ''
J_CON = ''

for i in range(len(Lines)):
    txt = Lines[i]
    if 'CSP_FILE_1' in txt:
        bc_data_folder_path += os.path.split(txt.split()[1])[0]
        csp_file_name += os.path.split(txt.split()[1])[1]
    if 'STORE_DATA' in txt:
        bc_data_folder += txt.split()[1]
    if 'FLAG_PRE' in txt:
        if int(txt.split()[1]) == 1:
            FLAG_PRE = 1
    if 'FLAG_J' in txt:
         if int(txt.split()[1]) == 1:
             FLAG_J = 1
    if 'FLAG_NOE' in txt:
         if int(txt.split()[1]) == 1:
             FLAG_NOE = 1
    if 'FLAG_SAXS' in txt:
         if int(txt.split()[1]) == 1:
             FLAG_SAXS = 1
    if 'FLAG_MRDC' in txt:
         if int(txt.split()[1]) == 1:
             FLAG_MRDC = 1
    if 'FLAG_SHIFTX' in txt:
         if int(txt.split()[1]) == 1:
             FLAG_SHIFTX = 1
             
bc_data_path = bc_data_folder_path + '/'+ bc_data_folder
   
if FLAG_PRE == 1:
    cdp_file_PRE = csp_file_name + '.PRE'
    os.system(f'python cdp.py {bc_data_path + "/"+cdp_file_PRE}')

    for i in range(len(Lines)):
        txt = Lines[i]
        if 'PRE_CONSTRAINT_FILE_NAME' in txt.split():
            PRE_CON += txt.split()[1]
            print(PRE_CON)
    bc_data_PRE = bc_data_path + '/'+cdp_file_PRE + '.HR'
    
if FLAG_NOE == 1:
    cdp_file_NOE = csp_file_name + '.NOE'
    
    os.system(f'python cdp.py {bc_data_path + "/"+cdp_file_NOE}')

    for i in range(len(Lines)):
        txt = Lines[i]
        if 'NOE_CONSTRAINT_FILE_NAME' in txt:
            NOE_CON += txt.split()[1]
            print(NOE_CON)
    bc_data_NOE = bc_data_path + '/'+cdp_file_NOE + '.HR'
        

if FLAG_J == 1:
    cdp_file_J = csp_file_name + '.J'
    
    os.system(f'python cdp.py {bc_data_path + "/"+cdp_file_J}')

    for i in range(len(Lines)):
        txt = Lines[i]
        if 'J_CONSTRAINT_FILE_NAME' in txt:
            J_CON += txt.split()[1]
    bc_data_J = bc_data_path + '/'+cdp_file_J + '.HR'

if FLAG_SAXS == 1:
    cdp_file_SAXS = csp_file_name + '.SAXS'
    
    os.system(f'python cdp.py {bc_data_path + "/"+cdp_file_SAXS}')

    for i in range(len(Lines)):
        txt = Lines[i]
        if 'SAXS_CONSTRAINT_FILE_NAME' in txt:
            SAXS_CON += txt.split()[1]
    bc_data_SAXS = bc_data_path + '/'+cdp_file_SAXS + '.HR'

if FLAG_MRDC == 1:
    cdp_file_MRDC = csp_file_name + '.MRDC'
    
    os.system(f'python cdp.py {bc_data_path + "/"+cdp_file_MRDC}')

    for i in range(len(Lines)):
        txt = Lines[i]
        if 'MRDC_CONSTRAINT_FILE_NAME' in txt:
            MRDC_CON += txt.split()[1]
    bc_data_MRDC = bc_data_path + '/'+cdp_file_MRDC + '.HR'

if FLAG_SHIFTX == 1:
    cdp_file_SHIFTX = csp_file_name + '.SHIFTX'
    
    os.system(f'python cdp.py {bc_data_path + "/"+cdp_file_SHIFTX}')

    for i in range(len(Lines)):
        txt = Lines[i]
        if 'SHIFTX_CONSTRAINT_FILE_NAME' in txt:
            SHIFTX_CON += txt.split()[1]
    bc_data_SHIFTX = bc_data_path + '/'+cdp_file_SHIFTX + '.HR'

if FLAG_PRE == 1:
    PRE_RMSD(bc_data_PRE,PRE_CON,n_conf)
if FLAG_SHIFTX == 1:
    CS_RMSD(bc_data_SHIFTX,SHIFTX_CON,n_conf)
if FLAG_SAXS == 1:
    SAXS_RMSD(bc_data_SAXS,SAXS_CON,n_conf)
if FLAG_J == 1:
    JC_RMSD(bc_data_J,J_CON,n_conf)
if FLAG_MRDC == 1:
    RDC_RMSD(bc_data_MRDC,MRDC_CON,n_conf)
if FLAG_NOE == 1:
    NOE_RMSD(bc_data_NOE,NOE_CON,n_conf)
```
