## Supplementary material for "IDPConformerGenerator: A Flexible Software Suite for Sampling Conformational Space of Disordered Protein States": Analysis scripts: CDP_Translator.py

```
#!/usr/bin/env python3
# -*- coding: utf-8 -*-
"""
Created on Wed Jul 21 12:48:45 2021

@author: Mickael Krzeminski
"""
import struct, sys, os
from pathlib import Path


__CHAR_SIZE__   = struct.calcsize('=c')
__SHORT_SIZE__  = struct.calcsize('=H')
__INT_SIZE__ = struct.calcsize('=i')
__LONG_SIZE__   = struct.calcsize('=L')
__DOUBLE_SIZE__ = struct.calcsize('=d')
__CARETURN__ = os.linesep


# Some help to explain how the script works
__USAGE__ = """

This program reads a binary CDP file as described in ENSEMBLE into a human readable format, which is transcripted in an output file that has the same name as the input file followed by ".HR".
	 Usage: %s file1 [file2] [file3] [...]

"""
%sys.argv[0]

if (len(sys.argv) < 2 or sys.argv[0] == "-h"):
		print (__USAGE__)
		sys.exit()


# Reading the specified files
filesToRead = sys.argv
filesToRead.pop(0)		# removing the first element
if (filesToRead[0] == __file__): filesToRead.pop(0)

for fileToRead in filesToRead:
	print ("%cReading %s..."%(__CARETURN__, fileToRead), end=" ")
	outFile = "%s.HR"%fileToRead

	# Checking the existence of the file and that it is a file
	if (not os.path.isfile(fileToRead)):
		print ("The file does not exist!")
		continue

	# Getting file size
	fileSize = 	Path(fileToRead).stat().st_size
	print("Size of the file:", fileSize)

	# Opening the file as binary
	cdpFile = open(fileToRead, "rb")

	# Checking it is a CDP file
	if (cdpFile.read(3*__CHAR_SIZE__).decode() != "Cdp"):
		print ("Not a CDP file!")
		continue

	# Reading the data type. Codes are the following:
	dataTypeList = {0: 'SHIFTX', 1: 'SHIFTX_CA', 2: 'SHIFTX_CB', 3: 'SHIFTX_C', 4: 'SHIFTX_N', 5: 'SHIFTX_HA', 6: 'SHIFTX_H',\
	                7: 'SAXS', 8: 'MRDC', 9: 'J', 10: 'NOE', 11: 'PRE', 12: 'R2', 13: 'HYDRO', 14: 'RPRE',\
					15: 'ACCESS', 16: 'ACCESS_0', 17: 'ACCESS_1', 18: 'ACCESS_2'}
	dataType = dataTypeList[struct.unpack('=H', cdpFile.read(__SHORT_SIZE__))[0]]
	print("Data type:", dataType)

	# Number of conformers
	nbConf = struct.unpack('=L', cdpFile.read(__LONG_SIZE__))[0]
	print("NbConf:", nbConf)

	# Size of a data block
	dataBlockSize = struct.unpack('=L', cdpFile.read(__LONG_SIZE__))[0]
	print("dataBlockSize:", dataBlockSize)

	# Number of data per block (each piece of data is a double-precision float)
	if (dataType =="R2"):
		s = __INT_SIZE__
	else:
		s = __DOUBLE_SIZE__
	nbData = int(dataBlockSize / s)
	print("nbData:", nbData)

	# Verifying the relevance of the file (based on its size)
	if (fileSize != 3 * __CHAR_SIZE__ + __SHORT_SIZE__ + 2 * __LONG_SIZE__ + nbConf * dataBlockSize):
		print ("Corrupted file!")
		continue
	
	# Writing down the data in human readable format
	hrdFile = open(outFile, 'w')
	hrdFile.write("Data of type %s for %d conformers (%d data per conformer)%c"%(dataType, nbConf, nbData, __CARETURN__))
	for confNb in range(nbConf):
		hrdFile.write("%cConformer #%d%c%s%c"%(__CARETURN__, confNb, __CARETURN__, "-"*(11+len(str(confNb))), __CARETURN__))
		for dataNb in range(nbData):
			if (dataType =="R2"):
				hrdFile.write("%d%c"%(struct.unpack('=i', cdpFile.read(__INT_SIZE__))[0], __CARETURN__))
			else:
				hrdFile.write("%.50f%c"%(struct.unpack('=d', cdpFile.read(__DOUBLE_SIZE__))[0], __CARETURN__))

	hrdFile.close()
	cdpFile.close()

	print("DONE. Results in %s"%outFile)

print ("%s"%__CARETURN__*4)
```
