## Supplementary material for "IDPConformerGenerator: A Flexible Software Suite for Sampling Conformational Space of Disordered Protein States": Analysis scripts: Format_CSCON_delta2D.py

```
#!/usr/bin/env python3
# -*- coding: utf-8 -*-
"""
Created on Fri Jun 18 00:31:16 2021
Reads ENSEMBLE ready chemical shift .CON files and converts them into delta2D
readable shifty format.

@author: Ashley Namini, Nemo (Zi Hao) Liu @menoliu
"""
import numpy as np

#Path to CON file used for ENSEMBLE
CONPATH = '/home/nemo/Documents/CON_data/asyn_cs.con'
OUTPATH = '/home/nemo/Documents/d2D_data/asyn_d2D_cs.txt'

con = np.loadtxt(CONPATH, dtype=str)

#FASTA sequence of protein of interest
seq_txt = 'MDVFMKGLSKAKEGVVAAAEKTKQGVAEAAGKTKEGVLYVGSKTKEGVVHGVATVAEKTKEQVTNVGGAVVTGVTAVAQKTVEGAGSIAAATGFVKKDQLGKNEEGAPQEGILEDMPVDPDNEAYEMPSEEGYQDYEPEA'
seq = np.array(list(seq_txt))

res = np.array(list(range(1,len(seq)+1)))

C = np.zeros(len(seq)) #C=CO
CA=np.zeros(len(seq))
CB=np.zeros(len(seq))
H=np.zeros(len(seq)) #H=HN
HA=np.zeros(len(seq))
N=np.zeros(len(seq))


for i in range(len(con)):
    r = int(con[i,0])
    if con[i,1] == 'C':
        C[r-1] = con[i,2] 
    elif con[i,1] == 'CA':
        CA[r-1] = con[i,2]
    elif con[i,1] == 'CB':
        CB[r-1] = con[i,2]
    elif con[i,1] == 'H':
        H[r-1] = con[i,2]
    elif con[i,1] == 'HA':
        HA[r-1]=con[i,2]
    elif con[i,1] == 'N':
        N[r-1]=con[i,2]


#CO = np.zeros(len(seq))
#N = np.zeros(len(seq))


        
d2D = np.column_stack([res, seq, HA,CA,CB,C,N,H])

#Saves the newly formatted delta2D input file for chemical shifts
np.savetxt(OUTPATH, d2D, delimiter=" ", newline = "\n", fmt="%s")
       


        


   
```
