## Supplementary material for "IDPConformerGenerator: A Flexible Software Suite for Sampling Conformational Space of Disordered Protein States": Analysis scripts: Get_RMSDs.py

```
#!/usr/bin/env python3
# -*- coding: utf-8 -*-
"""
Created on Tue Feb 15 23:11:02 2022
Variety of functions to calculate the RMSD between back-calculated experimental
data from ENSEMBLE and NMR experimental data.

@author: Ashley Namini
"""
import numpy as np

#This function is used for calculating the RMSD
def calc_RMSD(bc, exp):
    RMSD_num = abs(exp - bc)
    RMSD = np.sqrt(np.sum(RMSD_num**2)/len(exp))
    
    return RMSD

'''
This function prints the RMSD between the back-calculated SAXS data from ENSEMLE
and the experimental SAXS data.
'''
def SAXS_RMSD(bc_data_path, exp_data_path, n_conf):
    data = bc_data_path
    I_exp = np.loadtxt(exp_data_path,usecols = 1)
    nb_points = len(I_exp)
     
    file = open(data,'r')
    lines = file.readlines()
    
    SAXS_Data_arr = np.zeros(nb_points)
    
    
    for idx in range(2,len(lines)):
        if '------------' in lines[idx]:
            counter = 0
            for i in range(idx+2,idx+nb_points+1): 
                SAXS_Data_arr[counter] += float(lines[i])
                counter +=1  
    file.close()
    
    I_exp_sum = np.sum(I_exp)
    I_exp_norm = I_exp/I_exp_sum
    I_bc = SAXS_Data_arr/n_conf
    I_bc_norm = I_bc/np.sum(I_bc)
    RMSD_num = abs(I_exp_norm - I_bc_norm)
    RMSD = np.sqrt(np.sum(RMSD_num**2)/len(I_exp_norm))
    print(f'SAXS RMSD: {RMSD}')

    return RMSD

'''
This function prints the RMSDs between the back-calculated chemical shifts data from ENSEMBLE
and the experimental chemical shifts data.
'''
def CS_RMSD(bc_data_path,exp_data_path,n_conf):
    data = bc_data_path
    CS_exp = np.loadtxt(exp_data_path,dtype = str)
    nb_points = len(CS_exp)
    
    file = open(data,'r')
    lines = file.readlines()
    
    CS_Data_arr = np.zeros(nb_points)
    for idx in range(2,len(lines)):
        if '------------' in lines[idx]:
            counter = 0
            for i in range(idx+1,idx+nb_points+1):
                CS_Data_arr[counter] += float(lines[i])
                counter +=1
     
    file.close()

    CS_bc = CS_Data_arr/n_conf
    CS_bc_C = []
    CS_bc_CA = []
    CS_bc_CB = []
    CS_bc_H = []
    CS_bc_HA = []
    CS_exp_C = []
    CS_exp_CA = []
    CS_exp_CB = []
    CS_exp_H = []
    CS_exp_HA = []
    
    
    for j in range(len(CS_bc)):
        if CS_exp[j,1] == 'C':
            CS_bc_C.append(CS_bc[j])
            CS_exp_C.append(float(CS_exp[j,2]))
            
        elif CS_exp[j,1] =='CA':
             CS_bc_CA.append(CS_bc[j])
             CS_exp_CA.append(float(CS_exp[j,2]))

        elif CS_exp[j,1] == 'CB':
             CS_bc_CB.append(CS_bc[j])
             CS_exp_CB.append(float(CS_exp[j,2]))
             
        elif CS_exp[j,1] == 'H':
             CS_bc_H.append(CS_bc[j])
             CS_exp_H.append(float(CS_exp[j,2]))
             
        elif CS_exp[j,1] == 'HA':
              CS_bc_HA.append(CS_bc[j])
              CS_exp_HA.append(float(CS_exp[j,2]))
              
    print(f'C RMSD: {calc_RMSD(np.array(CS_bc_C),np.array(CS_exp_C))}')
    print(f'CA RMSD: {calc_RMSD(np.array(CS_bc_CA),np.array(CS_exp_CA))}')
    print(f'CB RMSD: {calc_RMSD(np.array(CS_bc_CB),np.array(CS_exp_CB))}')
    print(f'H RMSD: {calc_RMSD(np.array(CS_bc_H),np.array(CS_exp_H))}')
    print(f'HA RMSD: {calc_RMSD(np.array(CS_bc_HA),np.array(CS_exp_HA))}')

'''
This function prints the RMSD between the back-calculated PRE data from ENSEMBLE
and the experimental PRE data.
'''
def PRE_RMSD(bc_data_path,exp_data_path,n_conf):
    data = bc_data_path
    PRE_exp = np.loadtxt(exp_data_path,usecols = 4)
    nb_points = len(PRE_exp)
    
    file = open(data,'r')
    lines = file.readlines()
    
    PRE_Data_arr = np.zeros(nb_points)
    
    for idx in range(2,len(lines)): 
        if '------------' in lines[idx]:
            counter = 0
            for i in range(idx+1,idx+nb_points+1):
                PRE_Data_arr[counter] += float(lines[i])
                counter +=1
    
    file.close()
    
    PRE_exp = np.loadtxt(exp_data_path,usecols = 4)
    PRE_bc = PRE_Data_arr/n_conf
    PRE_bc = PRE_bc**(-1/6)
    
    print(f'PRE RMSD: {calc_RMSD(np.array(PRE_bc),PRE_exp)}')

'''
This function prints the RMSD between the back-calculated J-couplingdata from ENSEMBLE
and the experimental J-coupling data.
'''
def JC_RMSD(bc_data_path, exp_data_path,n_conf):
    data = bc_data_path
    J_exp = np.loadtxt(exp_data_path,usecols = 8)
    nb_points = len(J_exp)
    
    file = open(data,'r')
    lines = file.readlines()
   
    J_Data_arr = np.zeros(nb_points)
    
    for idx in range(2,len(lines)):
        if '------------' in lines[idx]:
            counter = 0
            for i in range(idx+1,idx+nb_points+1):
                J_Data_arr[counter] += float(lines[i])
                counter +=1
     
    file.close()
    
    J_bc = J_Data_arr/n_conf
    print(f'J {calc_RMSD(np.array(J_bc),J_exp)}')
    return
    
'''
This function printss the RMSD between the back-calculated RDC data from ENSEMBLE
and the experimental RDC data.
'''
def RDC_RMSD(bc_data_path,exp_data_path,n_conf):
    

    data = bc_data_path
    RDC_exp = np.loadtxt(exp_data_path,usecols = 4)
    nb_points = len(RDC_exp)
    
    file = open(data,'r')
    lines = file.readlines()
    
    RDC_Data_arr = np.zeros(nb_points)
    
    for idx in range(2,len(lines)):
        if '------------' in lines[idx]:
            counter = 0
            for i in range(idx+1,idx+nb_points+1):
                RDC_Data_arr[counter] += float(lines[i])
                counter +=1
      
    file.close()
    RDC_bc = RDC_Data_arr/n_conf
    
    print(f'RDC RMSD: {calc_RMSD(np.array(RDC_bc),RDC_exp)}')
    return
    
'''
This function prints the RMSD between the back-calculated NOE data from ENSEMBLE
and the experimental NOE data.
'''
def NOE_RMSD(bc_data_path,exp_data_path,n_conf):
    data = bc_data_path
    NOE_exp = np.loadtxt(exp_data_path,dtype = str,delimiter = '\t')
    nb_points = len(NOE_exp)
    
    file = open(data,'r')
    lines = file.readlines()

    NOE_Data_arr = np.zeros(nb_points)

    for idx in range(2,len(lines)):
        if '------------' in lines[idx]:
            counter = 0
            for i in range(idx+1,idx+nb_points+1):
               NOE_Data_arr[counter] += float(lines[i])
               counter +=1

    file.close()

    NOE_bc = NOE_Data_arr/n_conf
    NOE_bc = NOE_bc**(-1/6)
    nb_OR = 0
    counter_arr = []
    idx = 0
    counter = 0
    finished = False
    tolerance = len(NOE_exp)
    iteration = 0

    while not finished:
        atom1 = int(NOE_exp[idx][0])
        atom2 = int(NOE_exp[idx][2])
       
        while atom1 == int(NOE_exp[idx][0]) and atom2 == int(NOE_exp[idx][2]):
            idx +=1
            if idx >= len(NOE_exp):
                finished = True
                counter += 1
                break
            
            counter +=1
        counter_arr.append(counter)
       
        counter = 0
        iteration += 1
        
        if iteration >= tolerance:
            break
      
    idx = 0 
    NOE_min_dists = []         
    for nb in counter_arr:
        distances = [] 
        for i in range(nb):
            distances.append(NOE_bc[i+idx])
        NOE_min_dists.append(min(distances))
        idx += nb
        
    NOE_exp_dists = []
    for exp in NOE_exp:
        if not('OR' in exp):
            NOE_exp_dists.append(exp[4])
    NOE_exp_dists = np.array(NOE_exp_dists).astype(float)  
    NOE_min_dists = np.array(NOE_min_dists)

        
    print(f'NOE RMSD: {calc_RMSD(NOE_min_dists,NOE_exp_dists)}')
    return
```
