## Supplementary material for "IDPConformerGenerator: A Flexible Software Suite for Sampling Conformational Space of Disordered Protein States": Analysis scripts: Plot_Frac_Sec_Str_DSSP.py

```
# -*- coding: utf-8 -*-
"""
Created on Wed Jun 24 23:39:03 2020

@author: Gregory-Neal Gomes, Nemo (Zi Hao) Liu @menoliu
"""

import numpy as np
import glob
import matplotlib.pyplot as plt
import mdtraj

#reading conformer pools
POOL_NAME="drk_chunk112555_csss_nosub_faspr"
mypath0=f'/home/nemoliu/Documents/Conformers/drk_ANYvsCSSS_faspr/{POOL_NAME}/'
OUTPATH = f'/home/nemoliu/Pictures/fraction_SS/{POOL_NAME}_FSS.png'
files0=glob.glob(mypath0+"*.pdb")
traj0 = mdtraj.load(files0)

'''
Function below is the work of Gregory-Neal Gomes in Dr. Claudiu Gradinaru's lab at UofT

MDTraj DSSP implementation outputs letter codes, but we want to know fraction
of frames each residue is in each type of secondary structure.
Right now, I iterating through each frame, and each residue in each frame.
This is reasonably fast for 100-5000 conformations - but I am sure there is a
better way to do this.
Note if each conformation has equal weight (like the output of ENSEMBLE) then
weight=1/Nconf
For BME, I pass vector of w_opt.
'''

def quant_dssp_more(traj,w=np.array([])):
    #For an ensemble or trajectory (traj) find the probablity of each residue
    #being in coil, helix, extended.
    #See DSSP assignment code
    #http://mdtraj.org/1.9.3/api/generated/mdtraj.compute_dssp.html#mdtraj.compute_dssp
    helix_alpha=np.zeros(traj.topology.n_residues)
    extended=np.zeros(traj.topology.n_residues)
    beta=np.zeros(traj.topology.n_residues)
    helix_3_10=np.zeros(traj.topology.n_residues)
    helix_pi=np.zeros(traj.topology.n_residues)
    hbt=np.zeros(traj.topology.n_residues)
    bend=np.zeros(traj.topology.n_residues)
    coil=np.zeros(traj.topology.n_residues)
    
    if not w.any():
        #Allows for unequal weights (i.e., when I do BME reweighting)
        w=np.ones(len(traj))
        w=w/len(w)

    for i in range(len(traj)):
        dssp=mdtraj.compute_dssp(traj[i],simplified=False)[0]
        for j in range(len(dssp)):
            if 'H' in dssp[j]:
                helix_alpha[j]+=w[i]
            elif 'B' in dssp[j]:
                beta[j]+=w[i]
            elif 'E' in dssp[j]:
                extended[j]+=w[i]
            elif 'G' in dssp[j]:
                helix_3_10[j]+=w[i]
            elif 'I' in dssp[j]:
                helix_pi[j]+=w[i]  
            elif 'T' in dssp[j]:
                hbt[j]+=w[i]
            elif 'S' in dssp[j]:
                bend[j]+=w[i]
            else:
                coil[j]+=w[i]
    return helix_alpha, beta, extended, helix_3_10, helix_pi, hbt, bend, coil

#plotting preferences
font = {'family' : 'normal',
        'weight' : 'normal',
        'size'   : 20}
plt.rc('font', **font)
params = {'legend.fontsize': 10,
          'legend.handlelength': 1}
plt.rcParams.update(params)

#Obtaining and combining DSSP codes
H, B, E, G, I, T, S, C = quant_dssp_more(traj0)
L = C+S+T+I+B
He = H+G

#get residue numbers as they appear in PDB (eg., Sic1 starts from -1 in MD, but 1 in TraDES)
aa=[x.resSeq for x in traj0.topology.residues] 

#plotting
plt.figure()
plt.plot(aa,He,label='Helix',color = 'k',alpha = 0.5)
plt.plot(aa,E,label='Strand', linestyle = 'dashed',color = 'm')
plt.plot(aa,L,label='Loop', linestyle = 'dashed',color = 'r')
plt.ylabel('Fraction SS')
plt.xlabel('Residue')
plt.legend()
ax = plt.gca()

plt.savefig(OUTPATH, dpi=300, transparent=False, bbox_inches="tight")
```
