## Supplementary material for "IDPConformerGenerator: A Flexible Software Suite for Sampling Conformational Space of Disordered Protein States": Analysis scripts: Plot_Omega_distributions.py

```
"""
Last updated: May 23, 2022
Calculate omega distributions from IDPCG PDB database.

@author Joao M. C. Teixeira @joaomcteixeira
"""

import json
import sys
from collections import Counter
from math import degrees
#
from matplotlib import pyplot as plt


def center_at_zero(angle):
    """
    Center angle at 0.

    Facilitate representation.
    """
    return angle % 360 - 180


def analyse_angles(db, ss_considered, fig_name, fig_title):
    omega_angles = []  # all omega angles in the database
    for pdb, values in db.items():
        for v, ss in zip(values['omega'], values['dssp']):
            if ss in ss_considered:
                omega_angles.append(v)

    loa = len(omega_angles)
    print('Total number of OMEGA values is:', loa)


    omega_angles_no = [
        center_at_zero(degrees(i))
        for i in omega_angles
        ]


    limit = 50

    omega_centered_d5 = Counter(round(v, 5) for v in omega_angles_no)

    loc = len(omega_centered_d5)

    fig, ax = plt.subplots(nrows=1, ncols=1)
    h2 = ax.hist(
        omega_centered_d5.keys(),
        bins=160,
        range=(-limit, limit),
        weights=omega_centered_d5.values(),
        color="black",
        lw=0,
        )
    t2 = ax.set_title(fig_title)
    x2 = ax.set_xlabel('Degrees')
    y2 = ax.set_ylabel('Counts of angles')
    ax.set_xticks([-40, -30, -20, -10, 0, 10, 20, 30, 40])
    ax.grid(axis='x', ls='dotted', color='darkgrey')
    ax.set_xlim((-45, 45))
    fig.savefig(fig_name, dpi=300, bbox_inches='tight')
    plt.close()


if __name__ == "__main__":
    # get database name
    data_base_json = sys.argv[1]

    # open database to dictionary
    with open(data_base_json, 'r') as fin:
        db = json.load(fin)

    # edit the following file names and plot titles
    # according to the input database
    analyse_angles(
        db,
        ('L',),
        "all_omegas_res_18_L.png",
        "Omega angle distribution for PDBs $\leq$ 1.8$\AA$ - loops",
        )

    analyse_angles(
        db,
        ('H',),
        "all_omegas_res_18_H.png",
        r"Omega angle distribution for PDBs $\leq$ 1.8$\AA$ - helices",
        )

    analyse_angles(
        db,
        ('E',),
        "all_omegas_res_18_E.png",
        r"Omega angle distribution for PDBs $\leq$ 1.8$\AA$ - strands",
        )

    analyse_angles(
        db,
        ('L', 'H', 'E'),
        "all_omegas_res_18_all.png",
        r"Omega angle distribution for PDBs $\leq$ 1.8$\AA$",
        )
```
