## Supplementary material for "IDPConformerGenerator: A Flexible Software Suite for Sampling Conformational Space of Disordered Protein States": Analysis scripts: Plot_pwRMSD.py

```
#!/usr/bin/env python3
# -*- coding: utf-8 -*-
"""
Accepts PDB ensembles and calculates the pairwise-RMSD using MDAnalysis.
Ability to save matrix, plot as a heat-map, and histogram.

Reference: https://userguide.mdanalysis.org/stable/examples/analysis/alignment_and_rms/pairwise_rmsd.html
Author: Nemo (Zi Hao) Liu @menoliu
"""
import MDAnalysis as md
from MDAnalysis.analysis import diffusionmap, align
import glob
import matplotlib.pyplot as plt
import numpy as np

#If you have multiple ensembles ordered in folders of "NAME_SUBGROUP"
#Enter the NAME as `PROTEIN` and SUBGROUP as `GROUPS`
PROTEIN = 'I-2'
GROUPS = [
        'csss_CheSPI_nosub',
        'ANY_sub532',
        'ANY_nosub',
        'csss_d2D_sub532',
        'csss_d2D_nosub'
        ]
PATH2PICS = '/home/nemoliu/Pictures/'

# Loops through all the sub-folders and calculates/plots
for gr in GROUPS:
    #Path to folders of ensembles
    ens_path = f'/home/nemoliu/Documents/Conformers/{PROTEIN}_{gr}/'
    files = glob.glob(ens_path+'*.pdb')
    u = md.Universe(files[0],files)

    aligner = align.AlignTraj(u, u, select='name CA', in_memory=True).run()

    matrix = diffusionmap.DistanceMatrix(u, select='name CA').run()

    #Saves the matrix of pwRMSDs in a CSV
    np.savetxt(f"./{PROTEIN}_{gr}_output.csv", matrix.dist_matrix)
    
    #Plots and saves pwRMSD information as a histogram
    plt.hist(matrix.dist_matrix.flatten(), bins=np.arange(0, 60, 1), density=True,  histtype='step', color='blue')
    plt.xlabel(r'Pairwise RMSD ($\AA$)')
    plt.ylabel("1/2 Frequency")
    plt.savefig(f'{PATH2PICS}{PROTEIN}_{gr}_histogram.png', dpi=300, transparent=False)
    plt.show()
    
    #Settings for the distance-matrix heat map
    plt.imshow(matrix.dist_matrix, cmap='viridis')
    plt.xticks(fontsize=16)
    plt.yticks(fontsize=16)
    plt.colorbar(label=r'RMSD ($\AA$)')
    
    #Sets the size of the plot to save
    width = 6
    height = width / 1.618
    plt.rcParams["figure.figsize"]=(width, height)
    
    #Saves the distance matrix heat-map
    plt.savefig(f'{PATH2PICS}{PROTEIN}/{PROTEIN}_{gr}_matrix.png', dpi=300, transparent=False)
    plt.show()
    #Clears the plot for the next batch of conformers
    plt.clf()
```
