## Supplementary material for "IDPConformerGenerator: A Flexible Software Suite for Sampling Conformational Space of Disordered Protein States": Analysis scripts: Plot_Ramachandran.py

```
# -*- coding: utf-8 -*-
"""
Created on Mon Mar 14 16:31:25 2022

Script to plot torsion angles on the Ramachandran space using MDAnalysis.

Reference: https://docs.mdanalysis.org/1.0.0/documentation_pages/analysis/dihedrals.html

@author: Nemo (Zi Hao) Liu @menoliu
"""

import MDAnalysis as md
from MDAnalysis.analysis.dihedrals import Ramachandran
import matplotlib.pyplot as plt
import glob
import numpy as np

#Change the following variables to specify your input path and number of residues
POOL_NAME = 'drk_chunkD_csss_nosub_mcsce'
POOL_PATH = f'/home/nemoliu/Documents/Conformers/drk_ANYvsCSSS/{POOL_NAME}/N3000/'
NUMBER_RESIDUES = 59

files = glob.glob(POOL_PATH+'*.pdb')
u = md.Universe(files[0],files)

r = u.select_atoms(f"resid 1-{NUMBER_RESIDUES}")

#plotting preferences
font = {'family' : 'normal',
        'weight' : 'normal',
        'size'   : 12}
plt.rc('font', **font)
params = {'legend.fontsize': 10,
          'legend.handlelength': 1}
plt.rcParams.update(params)


#Scatterplot of the ramachandran space based on torsion angles in the ensemble
R = Ramachandran(r).run()
fig, ax = plt.subplots(figsize=plt.figaspect(1))
R.plot(ax=ax, color='k', marker='.', s=2, alpha=0.4)

'''
The following ranges for phi and psi angles for the alpha and beta regions of
the Ramachandran plot is as defined as the same in delta2D:
    
Determination of Secondary Structure Populations in Disordered States of 
Proteins Using Nuclear Magnetic Resonance Chemical Shifts
Carlo Camilloni, Alfonso De Simone, Wim F. Vranken, and Michele Vendruscolo
Biochemistry 2012 51 (11), 2224-2231
DOI: 10.1021/bi3001825
'''
#for alpha : ymin = 120/360 = 0.333, ymax = 225/360 = 0.625 (psi between -120 and 45 deg)
#for alpha : xmin = -180, xmax = 10 (phi between -180 and 10 deg)
plt.axvspan(xmin=-180, xmax=10, ymin=0.333, ymax=0.625, color='red', alpha=0.2, clip_on = True)

#for beta : psi between -180 and -120 or 45 and 180 (ymin=0, ymax = 0.333 
#           and ymin=0.625 and ymax = 1 respectively)
#for beta : phi between -180 and 0
plt.axvspan(xmin=-180, xmax=0, ymax=0.333, color='blue', alpha=0.2, clip_on = True)
plt.axvspan(xmin=-180, xmax=0, ymin=0.625, color='blue', alpha=0.2, clip_on = True)

rama_3D = R.angles
aa = [x+1 for x in range(u.atoms.n_residues)]
alpha = np.zeros(u.atoms.n_residues)
beta = np.zeros(u.atoms.n_residues)
coil=np.zeros(u.atoms.n_residues)
p=np.ones(len(u.trajectory))
p=p/len(p)

#Filters the ramachandran plots and notes each torsion
for f in range(len(rama_3D)):
    for r in range(len(rama_3D[f])):
        phi = rama_3D[f][r][0]
        psi = rama_3D[f][r][1]
        if (-180.0 < phi and phi < 10.0) and (-120.0 < psi and psi < 45.0):
            alpha[r] += p[f]
        elif (-180.0 < phi and phi < 0.0) and ((-180 < psi and psi < -120) or (45 < psi and psi < 180)):
            beta[r] += p[f]
        else:
            coil[r] += p[f]

#Plotting the fractional information based on DSSP defined cutoffs for the
#ramachandran space
plt.figure()
plt.plot(aa,alpha,label='Alpha', linestyle = 'dashed', color = 'r')
plt.plot(aa,beta,label='Beta', linestyle = 'dashed',color = 'b')
plt.plot(aa,coil,label='Other', linestyle = 'dashed',color = 'k', alpha=0.4)
plt.ylabel('Fractional Secondary Structure')
plt.xlabel('Residue')
plt.legend()
ax = plt.gca()
ax.set_xlim([1,NUMBER_RESIDUES])
plt.xticks(np.arange(1, NUMBER_RESIDUES, 10))
```
