## Supplementary material for "IDPConformerGenerator: A Flexible Software Suite for Sampling Conformational Space of Disordered Protein States": Analysis scripts: Plot_Sec_Str_Prop_delta2D.py

```
# -*- coding: utf-8 -*-
"""
Created on Mon Mar 14 15:08:03 2022
Plots fractional secondary structure information as seen in delta2D predictions
after processing with `idpconfgen csssconv -d2D`

@author: Nemo (Zi Hao) Liu @menoliu
"""

import numpy as np
import matplotlib.pyplot as plt
import json


#Paths to input and output
CSSS_NAME = 'csss_drk.json'
INPATH=f'/home/nemoliu/Documents/database/{CSSS_NAME}'
OUTPATH='/home/nemoliu/Pictures/fraction_SS/{CSSS_NAME}_FSS.png'

with open(INPATH) as json_file:
    csss_dict = json.load(json_file)
    aa = int(len(csss_dict))
    H = np.zeros(aa)
    L = np.zeros(aa)
    E = np.zeros(aa)
    for residues in csss_dict:
        for ss in csss_dict[f'{residues}']:
            ires = int(residues)
            if ss == 'L':
                L[ires-1] = csss_dict[f'{residues}'][ss]
            elif ss == 'H':
                H[ires-1] = csss_dict[f'{residues}'][ss]
            elif ss == 'E':
                E[ires-1] = csss_dict[f'{residues}'][ss]
                
    aa = [numAA+1 for numAA in range(aa)]
    
    #plotting fractional secondary structure information
    plt.figure()
    plt.plot(aa,H,label='Helix',color = 'k',alpha = 0.5)
    plt.plot(aa,E,label='Strand', linestyle = 'dashed',color = 'm')
    plt.plot(aa,L,label='Loop', linestyle = 'dashed',color = 'r')
    plt.ylabel('Fraction SS')
    plt.xlabel('Residue')
    plt.legend()
    ax = plt.gca()
    
    #saves the plot
    plt.savefig(OUTPATH, dpi=300, transparent=False, bbox_inches="tight")
```
