## Supplementary material for "IDPConformerGenerator: A Flexible Software Suite for Sampling Conformational Space of Disordered Protein States": Analysis scripts: Plot_Tertiary_Contacts.py

```
#!/usr/bin/env python3
# -*- coding: utf-8 -*-
"""
Created on Tue Oct 19 14:50:19 2021
Script to plot the Ca-Ca tertiary contacts using modules from MDAnalysis.
Also has the ability to plot the significant differences of Ca-Ca tertiary
contacts between two ensembles.

References:
https://userguide.mdanalysis.org/stable/examples/analysis/distances_and_contacts/distances_within_selection.html
https://doi.org/10.1016/j.bpj.2020.05.015

@author: Ashley Namini,Gregory-Neal Gomes, Nemo (Zi Hao @menoliu) Liu
"""

import MDAnalysis as md
import numpy as np
from MDAnalysis.analysis import distances
import matplotlib.pyplot as plt
import glob
import scipy.stats as stat
import copy

#Change the following parameters for input and output
INPATH='/home/nemoliu/Documents/Conformers/'
PROTEIN='Sic1'
drA='_chunk1_L+_sub532'
drB='_chunk5_L+_sub532'
OUTPATH='/home/nemoliu/Pictures/'

#mypath0 corresponds to ensemble A
mypath0=f'{INPATH}{PROTEIN}{drA}'
#mypath1 corresponds to ensemble B
mypath1=f'{INPATH}{PROTEIN}{drB}'

def get_dist_std_arr(mypath):

    files=glob.glob(mypath+"/*.pdb")
    
    #create universe 
    u_ens = md.Universe(files[0],files)
    
    ca=u_ens.select_atoms('name CA')  # a selection (AtomGroup)
    n_ca=len(ca)
    
    ind=0
    
    distr = []
    
    #iterate through all frames
    for ts in u_ens.trajectory:     
        #calc. all possible distancess for ca atoms
        distr += [distances.self_distance_array(ca.positions)] 
        ind += 1
    #array contains the distrubtion of distances for each i,j pair
    distr = np.array(distr) 
    distr = distr.T
    
    #compute std of each row of distr
    std_distr = np.std(distr, axis = 1) 
    #we want std_distr to now be an array in the same form as ens_sq 
    sq_std_distr_arr = np.zeros((n_ca,n_ca))
    triu_std_distr = np.triu_indices_from(sq_std_distr_arr, k=1)
    sq_std_distr_arr[triu_std_distr] = std_distr #upper triangle
    sq_std_distr_arr.T[triu_std_distr] = std_distr #lower triangle
    
    ens_sq_std = sq_std_distr_arr
    
    #get median distance for each i,j 
    median_distr = np.median(distr, axis = 1)
    #create array of dimensions (number residues,number residues)
    sq_dist_arr = np.zeros((n_ca, n_ca))
    #array of indices (all poss i,j ie. (0,1) (0,2) (0,3) ...)
    triu = np.triu_indices_from(sq_dist_arr, k=1) 
    sq_dist_arr[triu] = median_distr
    #transpose array 
    sq_dist_arr.T[triu] = median_distr 
    
    ens_sq_dist=sq_dist_arr
    return ens_sq_std, ens_sq_dist, distr, triu

def get_residue_number(mypath):
      files=glob.glob(mypath+"/*.pdb")
      u_ens = md.Universe(files[0],files)
    
      ca=u_ens.select_atoms('name CA')
      residues = ca.resids
      return residues

def plot_tertiary_contacts(residues, array,xticks,yticks, protein):
    fig, ax = plt.subplots()
    fig.subplots_adjust(left=0, bottom=.15, right=.99, top=.97,hspace = 0, wspace = 0)
    
    im = ax.pcolor(residues, residues, array, cmap='CMRmap_r')
    
    # plt.pcolor gives a rectangular grid by default
    # so we need to make our heatmap square
    ax.set_aspect('equal')
    ax.set_xticks(xticks)
    ax.set_yticks(xticks)
    ax.set_yticklabels(yticks)
    
    ax.tick_params(axis='x', labelsize=14)
    ax.tick_params(axis='y', labelsize=14)
    
    fig.colorbar(im)

    plt.xlabel('Residues',fontsize = '12')
    plt.ylabel('Residues', fontsize = '12')

    width = 6
    height = width / 1.618
    fig.set_size_inches(width, height)
    plt.savefig(f'{OUTPATH}{PROTEIN}_hm.png',dpi = 300,transparent=False, bbox_inches='tight')

    
std_A, dist_A, distr_A, triu_A = get_dist_std_arr(mypath0) 

std_B, dist_B, distr_B, triu_B = get_dist_std_arr(mypath1) 

diff_dist = abs(dist_A - dist_B)

diff_std = abs(std_A - std_B)

#filter distance and std arrays for differences that are not significant
diff_dist_filt = np.array(diff_dist)
diff_std_filt = np.array(diff_std)


for i in range(len(distr_A)):
    u,p = stat.mannwhitneyu(distr_A[i,:], distr_B[i,:])
    
    if p > 0.05:
        res_i = triu_A[0][i]
        res_j = triu_A[1][i]
        diff_dist_filt[res_i,res_j] = 0
        diff_dist_filt[res_j,res_i] = 0
        diff_std_filt[res_i,res_j] = 0
        diff_std_filt[res_j,res_i] = 0
        
upper_dist_A = np.triu(dist_A,k=1)
lower_std_A = np.tril(std_A,k=0)


upper_dist_B = np.triu(dist_B,k=1)
lower_std_B = np.tril(std_B,k=0)

upper_diff_dist = np.triu(diff_dist, k=1)
lower_diff_std = np.tril(diff_std,k=0)

upper_diff_dist_filt = np.triu(diff_dist_filt, k=1)
lower_diff_std_filt = np.tril(diff_std_filt,k=0)

#The following 4 arrays are the array that will be used for plotting
dist_std_A = upper_dist_A + lower_std_A

dist_std_B = upper_dist_B + lower_std_B

diff_dist_std = upper_diff_dist + lower_diff_std

diff_dist_std_filt = upper_diff_dist_filt + lower_diff_std_filt

#call function get_residue_number to get the residue numbers in the protein
residues = get_residue_number(mypath0)

#change value of increment to control the labels on the figures
xticks = [1, 10, 20, 30, 40, 50, 60, 70, 80, 90]
yticks = copy.deepcopy(xticks)

#Plots distance map of ensemble A
plot_tertiary_contacts(residues, dist_std_A, xticks,yticks, PROTEIN)
#Plots distance map of ensemble B
plot_tertiary_contacts(residues, dist_std_B, xticks,yticks, PROTEIN)
#Plots the differences between A and B
plot_tertiary_contacts(residues, diff_dist_std, xticks,yticks, PROTEIN)
#Plots the significant differences between A and B
plot_tertiary_contacts(residues, diff_dist_std_filt, xticks,yticks, PROTEIN)
```
