## Supplementary material for "IDPConformerGenerator: A Flexible Software Suite for Sampling Conformational Space of Disordered Protein States": Analysis scripts: Plot_Torsion_Angles.py

```
#!/usr/bin/env python3
# -*- coding: utf-8 -*-
"""
Created on Wed Sep 22 21:05:42 2021
Script to plot the Phi and Psi (optional Omega) torsion angle distributions
of ensembles of PDBs using MDAnalysis packages

References:
https://userguide.mdanalysis.org/stable/examples/analysis/structure/dihedrals.html
https://docs.mdanalysis.org/1.0.0//documentation_pages/analysis/dihedrals.html
https://userguide.mdanalysis.org/stable/examples/analysis/structure/dihedrals.html
    
@author: Ashley Namini, Nemo (Zi Hao) Liu @menoliu
"""
import MDAnalysis as md
from MDAnalysis.analysis import dihedrals
import glob
import numpy as np
import matplotlib.pyplot as plt

#specify protein so plots will be saved in folder for that specific protein
PROTEIN = 'drk'
#specify the parent folder of protein ensembles
INPATH='/home/nemoliu/Documents/Conformers/'
#specify the output folder for the plots
OUTPATH='/home/nemoliu/Pictures/torsions/'
#list of subfolders for multiple ensemble analysis of torsion angles
dr_list = ['E+_nosub',
           'H+_nosub',
           'H+E+_nosub',
           'L+H+E+_nosub',
           'L+_nosub',
           'L+E+_nosub',
           'L+H+_nosub',
           ]

#number of conformers in ensemble. This variable is used for creating the scatter plots
n_conf = 1000

for dr in dr_list:
    #path to folder of pdbs
    mypath =f'{INPATH}{PROTEIN}_nosub_LHE/{PROTEIN}_{dr}'
    
    files = glob.glob(mypath+'/*.pdb')
    u = md.Universe(files[0],files)
    
    protein = u.select_atoms('protein')
    print('There are {} residues in the protein'.format(len(protein.residues)))
    
    phis = []
    psis = []
    omegas = []
    chi1s = []
    
    #can't start at 0 because needs four atoms in the atomgroups
    for res in u.residues[1:len(protein.residues)]: 
        phi = res.phi_selection()
        phis.append(phi)

    for res in u.residues[0:len(protein.residues)-1]:
        psi = res.psi_selection()
        psis.append(psi) 
        omega = res.omega_selection()
        omegas.append(omega)
    
    dihs_phi = dihedrals.Dihedral(phis).run()
    phi_ang = dihs_phi.angles
    dihs_psi = dihedrals.Dihedral(psis).run()
    psi_ang = dihs_psi.angles
    dihs_omega = dihedrals.Dihedral(omegas).run()
    omega_ang = dihs_omega.angles
    
    res_phi = list(range(2,len(protein.residues)+1))
    res_psi = list(range(1,len(protein.residues)))
    res_omega = list(range(1,len(protein.residues)))
    
    #plotting preferences
    font = {'family' : 'normal',
            'weight' : 'normal',
            'size'   : 12}
    plt.rc('font', **font)
    
    #This can be changed based on user preference
    xticklabels = [1, 10, 20, 30, 40, 50]
    
    #create scatter plot of psi angle distribution
    plt.figure(figsize = (10,6))
    for i in range(1,len(protein.residues)):
        res_psi = [i]*n_conf
        plt.scatter(res_psi,psi_ang[:,i-1],s=10, facecolors='none', edgecolors='k')
    plt.xticks(ticks=np.arange(0,len(protein.residues), 10), labels=xticklabels, fontsize=20)
    plt.yticks(fontsize=20)
    plt.ylabel('Psi',fontsize = 20)
    plt.xlabel('Residue')
    plt.title(f'{PROTEIN} {dr}')
    plt.savefig(f'{OUTPATH}psi_{PROTEIN}_{dr}.png',dpi = 300,transparent=False, bbox_inches='tight')
    
    #create scatter plot of phi angle distribution
    plt.figure(figsize = (10,6))
    for i in range(1,len(protein.residues)):
        res_phi = [i+1]*n_conf
        plt.scatter(res_phi,phi_ang[:,i-1],s=10, facecolors='none', edgecolors='k')
    plt.xticks(ticks=np.arange(0,len(protein.residues), 10), labels=xticklabels, fontsize=20)
    plt.yticks(fontsize=20)
    plt.ylabel('Phi',fontsize = 20)
    plt.xlabel('Residue')
    plt.title(f'{PROTEIN} {dr}')
    plt.savefig(f'{OUTPATH}phi_{PROTEIN}_{dr}.png',dpi = 300,transparent=False, bbox_inches='tight')

    
```
