## Supplementary material for "IDPConformerGenerator: A Flexible Software Suite for Sampling Conformational Space of Disordered Protein States": Analysis scripts: PRE_Ratio_Converter.py

```
#Calculator that takes user-defined variables and Iox/Ired ratios and converts them to PRE distances in cm
#Requires wolframalpha to be installed to anaconda distribution
#Author: Nemo (Zi Hao) Liu @menoliu

import wolframalpha
import numpy as np

#populate with to your own wolframalpha AppID after making an account
app_id = ""
client = wolframalpha.Client(app_id)

#Change paths and names of Iox/Ired ratio input files
PATH_TO_PRE_RATIOS = 'C:/Users/nemol/Desktop/asyn_PRE_ratios/'
NAME_OF_CSV = 'Exp_asyn_PRE_20'

output = open(PATH_TO_PRE_RATIOS+NAME_OF_CSV+"_converted.txt", "w")
ratios = np.loadtxt(PATH_TO_PRE_RATIOS+NAME_OF_CSV+'.csv', delimiter=',')

#Populate constants with experimental data/parameters
K = 1.23e-32 # units are cm^6 s^-2
tc = 4e-9 #units are s
w = 700 #units are MHz
R2red = 4 #units are s^-1
#t = 0.01 #units are s
t = 0.004 #units are s

alpha = K*((4*tc) + ((3*tc)/(1+((w**2)*(tc**2)))))

#beta value represents Iox/Ired
#script can be iterable through a loop to read .txt files or CSV files
#'r' function is doing the heavy lifting
beta = 0.026
r = client.query('solve ((%s*%s)/%s) = r^6 * (exp(-(%s*%s)/r^6) - %s) for r over the reals' %(beta, alpha, R2red, alpha, t, beta))
print(((next(r.results).text).strip("r = ± ")).replace("×10^", "E"))
```
