## Supporting Information for "IDPConformerGenerator: A Flexible Software Suite for Sampling Conformational Space of Disordered Protein States"

### ADDITIONAL METHODS

*Technical specifications of hardware.* For all calculations unless otherwise specified, we used one node of the Graham resource of Compute Canada (now Digital Research Alliance of Canada) with 2x Intel E5-2683 v4 Broadwell @ 2.1GHz CPUs and with 125GB of RAM per node.

*Default parameters.* We used a number of default parameters when building our test ensembles. We employed a fixed non-redundant database of PDB structures from the Dunbrack PISCES database[1] (October 15th, 2020), including high-resolution structures with resolution better than or equal to 2.0 Å and R-factor of 0.25 or lower with a maximum mutual sequence identity of 90%. This is a compromise to increase the number of high-quality backbone torsion angles without adding redundant information that will slow down the process. This Dunbrack PISCES database included a total of 24,003 PDB IDs, with some IDs referring to multiple chains from the same PDB file, leading to 22,599 PDB files ("cullpdb_pc90_res2.0_R0.25_d201015_chains24003"). These files contain 33,230 distinct chains from which we extract torsion angles and secondary structures. When not explicitly testing the impact of fragment sizes, we used the default of 10%, 10%, 30%, 30% and 20% probabilities for fragment sizes of 1, 2, 3, 4 and 5 residues, respectively. For MC-SCE, we used the simple mode with 128 trials per backbone structure.

*Creating the torsion angle database:* IDPConformerGenerator's initial database contains the primary sequence, secondary structure, and torsion angles (omega-phi-psi) from individual continuous protein chains derived from protein coordinates. During the preparation of the torsion angle database, the first and last residues of each protein are discarded to reduce artifacts in torsion angles. To create this database, IDPConformerGenerator has a series of commands that can be used in a modular fashion. First, the PDB downloader command (`pdbdl`) downloads a list of PDB IDs from the RCSB databank. Both PDB and mmCIF format files are accepted. Listed PDB IDs have the form of ABCDX where ABCD is the PDB ID and X is the optional chain id. If given, only that chain is stored in disk. We recommend using a culled list of PDB IDs from the Dunbrack PISCES database (http://dunbrack.fccc.edu/lab/pisces)[1]. The PDB downloader client further processes each PDB file by removing known solvent and ligands, considering only elements composed of amino acids, considering only the alternative location 'A' or ' ' and selecting only the first model for multi-model structures. If no chain ID is provided, IDPConformerGenerator will save each chain of the PDB file separately, and treat them as individual structures in subsequent steps. In this step, the user can easily add or remove PDBs to feed the sampling process. PDB files can also be added manually to the set of downloaded PDBs.

After the PDB files have been downloaded, the secondary structure calculator client (`sscalc`) annotates the secondary structure information from the PDBs using third-party software. We have implemented parsers for DSSP[2,3], but IDPConformerGenerator code structure allows for facile implementation of additional parsers. Because DSSP v2 and v3 do not consider polyproline type II helices, we adapted the DSSP-PPII Perl script[4] to work with DSSP v2 or v3 annotate poly-proline type II helices as "P". The output of the `sscalc` client produces a JSON file containing aligned primary sequences, residue numbers and DSSP annotations. Additionally, the input PDB files are split according to chain gaps, if existent, and resulting chains treated as individual PDBs. Here, segments shorter than 3 (configurable) are discarded. We suggest using the "reduce" flag (`-rd`), which directs IDPConformerGenerator to group DSSP codes into either Loops (L), Helices (H), or Extended strands (E). IDPConformerGenerator's definition of a loop incorporates DSSP's hydrogen-bonded turn (T), bend (S), beta-bridge (B), loops ('space'), poly-proline type II helix (P), and pi-helix (I); helices include DSSP's alpha-helix (H), and 3-10 helix (G); extended strands (E) are identical to DSSP's extended strands that participate in beta-ladder structures[5].

Finally, the torsion angles for each PDB are calculated using the `torsions` command in IDPConformerGenerator. The torsion angle calculator client calculates phi, psi, and omega torsion angles from the coordinates. On top of all of IDPConformerGenerator's filters in previous steps, the integrity of the PDB files is further confirmed at this stage, most importantly, by ensuring input structures have all required backbone atoms and no gaps. The torsion angle calculator client stores all the successfully processed PDB IDs aligned with their secondary structure code, primary sequence, and per-residue torsion angles in a JSON file. This is the initial IDPConformerGenerator database that will be used during the building algorithm. We chose a JSON structure because the amount of data used in this step, despite being large, is still workable within a JSON file, and because we advocate for using text-editable files as much as possible in research software pipelines for the sake of traceability, editing, and easy reviewing.

Once the database file has been created, it can be used to build conformers of any protein sequence, and can be shared among collaborators. The database only needs to be recreated if different sets of structures are desired (e.g., including new structures or using different resolution cutoffs). Otherwise, users only need the primary sequence of the disordered protein of interest. If users want to bias secondary structure propensities by NMR chemical shifts, they need an NMR STAR v2 file[6] for CheSPI[7] or SHIFTY[8] format for $\delta$2D[9]. The outputs of secondary structure propensity predictions from CheSPI or δ2D can be directly fed into the custom secondary structure sampling conversion `cssconv` client which will read the fractional secondary structure information and standardize them for each secondary structure code on a per-residue basis for the conformer building module. The standardized CSSS input file can also be edited in the `makecsss` module. If no chemical shift-based secondary structure propensity prediction exists, however, the user can specify secondary structure biases using `makecsss`, as well. The fact that the database preparation process is separated into three commands allows advanced users to manually alter PDB files, add or remove them from the JSON files, or otherwise intervene in the different steps, if needed.

*Building conformers with IDPConformerGenerator:* IDPConformerGenerator builds coordinates by sampling torsion angles from the database file. Therefore, the user has complete control over which PDBs will be used to drive conformational sampling. IDPConformerGenerator's `build` command-line interface requires the database file (created in the previous steps) and the IDP target sequence. Several details of the building process can be configured through the client's parameter flags. The most important will be explained here, and the complete list is available in the documentation pages, or via the `IDPConformerGenerator build -h` menu. Before starting building the actual coordinates, IDPConformerGenerator maps the input sequence to the respective positions in the input database. For this, the input protein sequence is split into fragments of different sizes (described below). Identified fragments allow overlap, i.e., abcde has three fragments of 3 residues: abc, bcd, and cde. Then, IDPConformerGenerator searches the input database for positions matching the fragments generated from the input protein. IDPConformerGenerator considers fragments preceding proline differently, i.e., abcP is treated differently than abcX, where X is not P[10,11]. Users can also specify a list of residues allowing substitutions to increase the likelihood of matching. For example, A → AV allows the input fragment GAG to match GAG or GVG sequences in the database. This process happens internally and no additional files are saved to disk.

Users specify the fragment sizes utilized to build conformers. Different from other tools to build disordered chains, IDPConformerGenerator allows the user to select whatever sizes for the fragments and the probabilities by which they are sampled during the building process. By default, IDPConformerGenerator uses fragment sizes 1, 2, 3, 4, and 5 with relative probabilities of 10%, 10%, 30%, 30%, and 20%, respectively.
Importantly, the user can sample fragment torsion angles specifically for user-defined secondary structure annotations. For example, fragments can be searched for residues in the database annotated as "loop" and/or "sheet" and/or "helix" regions depending on the user choice, using the `-rd` flag in the `sscalc` client. With these options, fragments consist only of residues matching a single secondary structure annotation. Alternatively, the user can specify "--dany" to allow fragment matching to disregard the secondary structure annotation. Moreover, users can specify sampling specific secondary structure annotations for distinct parts of the input protein and in certain probabilities. This option allows, for example, conformers to directly sample alpha-helices for certain residue ranges of the sequence, matching experimental data. To benefit from this feature, users need to prepare the required CSSS files as noted in the previous section. These options allow a rich combination of sampling possibilities adaptable to the project requirements.

Generation of structures requires transformation from internal to Cartesian coordinates. The default approach used for this work is based on an internal database built from structures with resolutions better than 1.6Å created with the bgeo command line interface. The database is hierarchically organized with bond angles separated by type, by residue type and residue neighbors, placed in 10° bins. While building backbones, the angles are statistically sampled from these values stored in the bgeo database. The bond lengths are taken as fixed values for each backbone atom pair and residue type, and are calculated from the same structures.

IDPConformerGenerator also ships with the recently developed Int2Cart[12] recurrent neural network algorithm for transformation from internal to Cartesian coordinates, currently integrated as a flag `--bgeo-strategy int2cart` within IDPConformerGenerator. This deep learning model provides improved prediction of bond lengths and bond angles from backbone ω, ϕ, and ψ torsion angles and residue types of a protein, relative to using fixed bond lengths and bond angles, or the original IDPConformerGenerator statistical approach used for most ensemble calculation in this work. Running Int2Cart within IDPConformerGenerator requires CUDA compatible GPUs, with at least two good GPUs needed for optimal performance if it is utilized at each fragment incorporation step. More comprehensive benchmarks and optimization of Int2Cart are forthcoming, with the goal to exploit the improved geometry that Int2Cart provides with no speed compromises. In addition to being implemented within IDPConformerGenerator, the Int2Cart algorithm is also available as an individual Python package at https://github.com/THGLab/int2cart.

IDPConformerGenerator can create backbone-only or full sidechain-containing conformers using the recommended Monte Carlo Side Chain Entropy (MC-SCE) algorithm[13] or FASPR[14]. The rationale during the process is however the same: conformers are accepted if the final energy is below the threshold. Accepted conformers are saved to disk. The conformer building process is an embarrassingly parallel process, for which users can give a number of cores across which the conformers will be distributed: 1000 conformers in 10 CPUs results in each CPU building 100 conformers, simultaneously. Also, IDPConformerGenerator is deterministic, and the building process can be reproduced by giving the same input and the same seed number.

*Utilities available and technical information:* As IDPConformerGenerator has been designed with modularity in mind, certain functions could be used as Python libraries for scripting or integration into other programs. One example of this is the parser for PDB files. Since IDPConformerGenerator parses only the labels and coordinate data from PDB and PDBx/mmCIF files, its parser module is much faster at processing structure files than other Python tools, including the highly utilized BioPython, or alternatives such as pdb2sql. IDPConfGen is approximately 2.5x faster than BioPython and 1.4x faster than pdb2sql for processing PDB files and approximately 847x faster than BioPython for mmCIF files, testing the same 22,599 structures with variable protein lengths and complexities in both PDB and mmCIF formats (without mulit-processing). BioPython had 48 errors due to these PDB IDs having only mmCIF file formats, while IDPConformerGenerator had no errors. Note that pdb2sql does not handle the mmCIF file format. Therefore, IDPConformerGenerator's PDB parsing algorithm could be imported as a library for custom scripts and computational processing of structure files.

Other examples are tools for analysis of torsion angles and fractional secondary structure of ensembles and for analysis of the database to provide histograms of matching fragment sequences and to search PDB headers for keywords of interest. For a basic analysis of conformer ensembles, there are three plotting functions within the “torsions” and “sscalc” sub-clients. Within “torsions”, given a folder/tarball of generated PDBs, a torsion angle distribution scatter plot can be plotted as well as fractional secondary structure information based on Ramachandran ranges for Phi/Psi angles. Fractional secondary structure based on DSSP codes can be plotted in the "sscalc" sub-client.

IDPConformerGenerator is written in Python, its source is open and can be found at Julie Forman-Kay's GitHub repository (https://github.com/julie-forman-kay-lab/IDPConformerGenerator). While IDPConformerGenerator is written all in Python, we incorporated a modification of the C++ code for the FASPR sidechain building tool in our repository to facilitate its usage (license compatible). For further details, please refer to the usage examples on the ReadTheDocs webpage (https://IDPConformerGenerator.readthedocs.io), as well as within the help documentation in IDPConformerGenerator, providing a guide to users of all levels from installation on multiple operating systems, to updating, and to using IDPConformerGenerator along with all of its functions.

*Monte Carlo Side Chain Entropy (MC-SCE).* The original code for MC-SCE[13] was re-written and implemented in Python, incorporating elements of the IDPConformerGenerator code. For each side chain other than glycine and alanine, the candidate torsion angle rotamers and their prior probabilities are obtained from the Dunbrack library[1] with the specific backbone $\phi/\psi$ values rounded to the closest tenth. For the N-terminal residue that does not have $\phi$ and the C-terminal residue that does not have $\psi$, $-180°$ was used as the undefined backbone torsion angle. The torsion angles $\chi_1$ and $\chi_2$ were extended by a standard deviation, i.e. $\chi_i \pm \sigma$ were also considered as rotamer candidates in addition to $\chi_i$ itself. Accordingly, probabilities for all rotamers from residues that only have $\chi_1$were set to 1/3 of their original probabilities in the Dunbrack library, and those from residues having and $\chi_2$ were set to 1/9 of the original probabilities. Torsion angles beyond $\chi_2$ were not expanded.

At each side chain growing position, let $\{\nu_m\}$ denote all candidate rotamers, the probability for each rotamer state to be selected was defined by:

$$P(\nu_k) = \frac{p^{(prior)}(\nu_k)e^{-\beta E(\nu_k)}}{\sum_{\{\nu_m\}} p^{(prior)}(\nu_m)e^{-\beta E(\nu_m)}}$$

where $p^{(prior)}(\nu_m)$ was the prior probability for the rotamer $\nu_m$ from the Dunbrack library scaled by 1/3 or 1/9, $E(\nu_m)$was the energy of the peptide structure with partially completed side chains up until the current building position, and $\beta = \frac{1}{k_B T}$ where $k_B$ was Boltzmann's constant and $T$ was selected as 300K. In this study only the 12-6 Lennard Jones term and a pairwise distance-based clash term were used for energy evaluation. The Lennard Jones term followed the implementation of Amber14[15] force field, and the clash term set the energy to ∞ if the distance for any pair of atoms in the structure was smaller than 0.6 times the sum of the van der Waals radii of the two atoms. We intentionally kept minimal terms in the energy function to avoid computational burden. If the clashes were inevitable, the algorithm selects the lowest energy conformation. Further details can be found in the MCSCE project (https://github.com/THGLab/MCSCE).

While it is possible to run MC-SCE through IDPConformerGenerator as a side chain method (`-scm`), for longer protein sequences we would advise running MC-SCE separately after the backbones have been successfully generated by IDPConformerGenerator. This is due to the current integration of MC-SCE within IDPConformerGenerator requiring MC-SCE to run immediately after each successful generation of a backbone. Although both programs are capable of Pythonic multiprocessing, running MC-SCE with multiple trials within IDPConformerGenerator will significantly stress the single worker responsible for calculating that conformer. For example, when requesting the typical 128 trials for the simple mode, if we specified 32 workers in IDPConformerGenerator, each of those workers would be independently generating backbones which then require 128 sidechain trials, leading to 4096 possible concurrent processes. In contrast, running MC-SCE with the same settings once all backbones have been generated would distribute the 128 trials among 32 workers, each worker running 4 calculations as opposed to 128.

*Calculating Ensembles with FastFloppyTail.* In order to use FastFloppyTail, input files are required to generate fragment libraries. We used the input files for Sic1 and alpha-synuclein provided by the authors of FastFloppyTail[16]. For the drkN SH3 domain, the sequence used by the authors of FastFloppyTail was different (one residue substitution) so we regenerated all the input files. We also created our own for inhibitor-2 and Tau since those were not modelled in the FastFloppyTail study. For proteins for which we needed to create input files, we used the suggested "best-reweighting scheme" to reweight the secondary structure predictions generated with RaptorX[17] and PSIPRED[18]. The disorder prediction was done using RaptorX, since this is the only predictor the authors found to predict alpha-synuclein as disordered. We created fragment libraries for all sequences using these input files and the Quota protocol, with the times required to create these libraries provided in **Supplementary Table 2C**. For calculating the structures, we used the 3-mer fragment libraries, ignoring the 9-mer fragment libraries. We generated ensembles with FastFloppyTail for all sequences other than the drkN SH3 domain using the suggested approach using the sequence and the fragment library as inputs, without supplying a disorder prediction file so that the sequence is treated as fully disordered, since RaptorX predicted all sequences other than the drkN SH3 domain to be largely disordered. For the unfolded state of the drkN SH3 domain, we ran two calculations, one treating it as partially ordered and disordered as RaptorX predicts, using the default protocol for a sequence with more than 20% predicted order as 42% of residues were predicted to be ordered. This entails first running AbInitioVO and providing the lowest energy output to use in FastFloppyTail along with the disordered probability prediction from RaptorX so that any ordered regions are maintained. For the second set of calculations for the unfolded state of the drkN SH3 domain, we treated it as fully disordered. For these, we edited the disorder probability prediction file from the RaptorX server, changing all residues predicted to be ordered to disordered with probability 1.0. This edited disorder file was used to reweight the secondary structure predictions with the best-reweighting scheme and to then generate the fragment libraries. AbInitioVO was not utilized and we did not supply FastFloppyTail with the disorder prediction file so the sequence was treated as fully disordered.

*Analysis of Conformational Ensembles and Experimental Data Sources.* To analyze the generated ensembles for fractional secondary structure information, torsion angle distribution, and tertiary contacts, well established Python libraries were used from MDAnalysis[19,20] and MDTraj[21]. For the analysis of pairwise RMSDs, the `diffusionmap` and `align` functions were used from the MDAnalysis `analysis` library. To calculate secondary structure information as DSSP codes, we used the `compute_dssp` function in MDTraj. Alpha, beta, and coil regions on the Ramachandran plot were determined using the `Ramachandran` function in the MDAnalysis `dihedrals` library. Tertiary contacts

(Cα-Cα) analysis was done with the `distances` function in the `analysis` library of MDAnalysis. Statistics performed for tertiary contacts analysis used the `stat` function from the `stats` library in SciPy.

RMSD values to experimental restraints and scores were calculated using ENSEMBLE[22] and X-EISD[23,24]. ENSEMBLE was modified to generate human-readable back-calculated data and to enable calculation of ENSEMBLE scores without optimizing. NMR and SAXS experimental data for comparison to ensembles were taken from supplemental information of papers or from our previous publications, and, for some chemical shifts, from the Biological Magnetic Resonance Data Bank (BMRB[25]). For the drkN SH3 domain unfolded state and I-2, data were from Marsh & Forman-Kay (2012)[26], with I-2 chemical shifts also in BMRB (ID 15179). For Sic1, data were from Gomes et al (2020)[27], with chemical shifts also in the BMRB (ID 16657). For alpha-synuclein, RDC, $^{3}J$-coupling, chemical shifts, and PRE restraints were from Ferrie et al (2020)[16] and SAXS restraints were from Pesce & Lindorff-Larsen (2021)[28]. For Tau chemical shifts, BMRB IDs: 17920 and 50701 were used and for SAXS, data were also from Pesce & Lindorff-Larsen (2021)[28]. Chemical shifts for the drkN SH3 domain unfolded state and I-2 were also used for δ2D[9] and CheSPI[7] calculations.

**SUPPLEMENTARY FIGURES.** All plots, in the main text and Supplement, were generated using Matplotlib[29] unless otherwise specified.

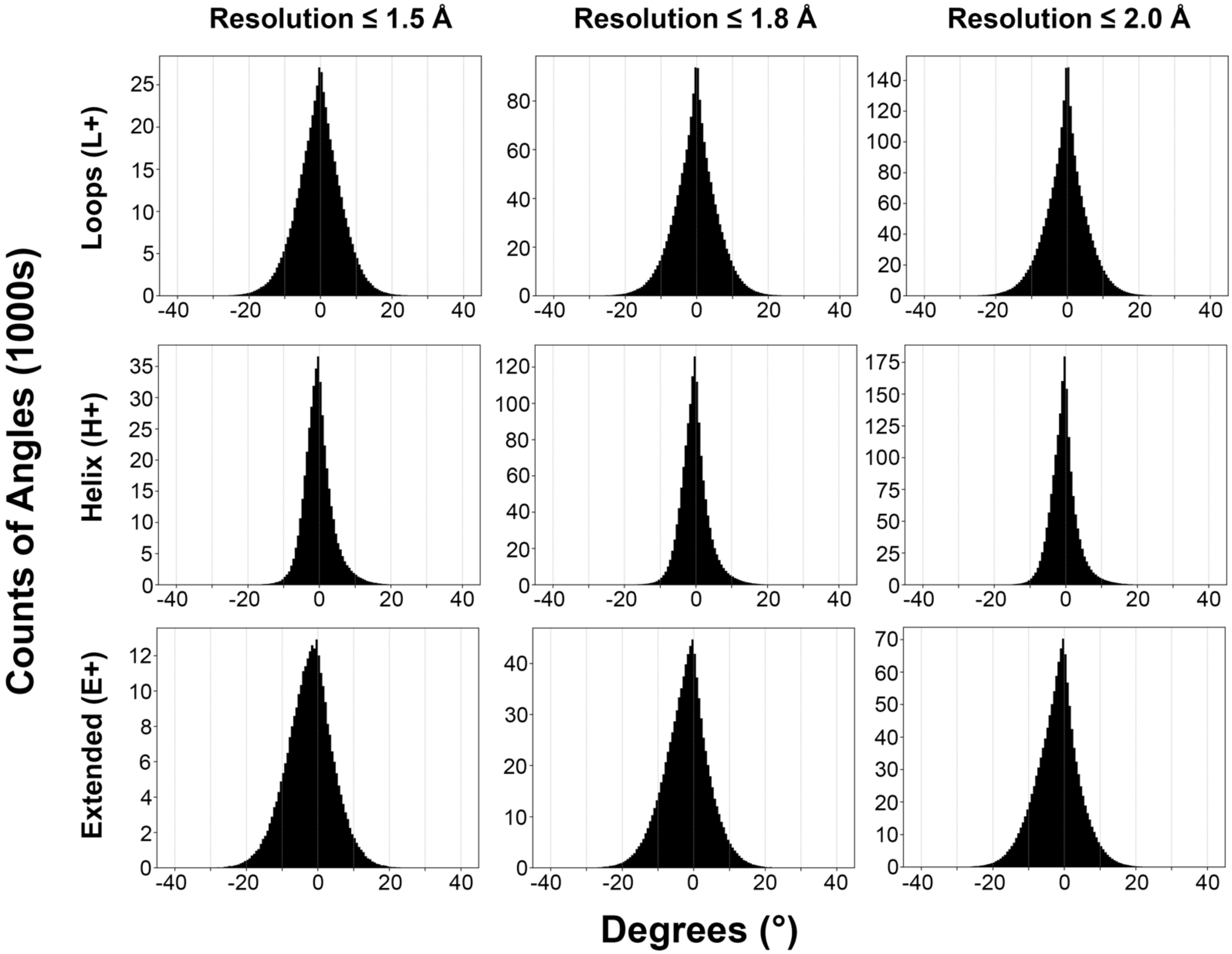


**Supplemental Figure S1.** Histogram of omega dihedral angle distributions for structures found within the IDPConformerGenerator database. PDBs from the 24,003 PDBID database were used, with sets of this with resolutions better than 1.5Å (~5,000 structures), better than 1.8Å (~16,000 structures) and better than 2.0Å (full set). The deviations from an omega torsion angle of 180° (trans) is plotted. values are centered around 0o to facilitate visualization of the distribution. Cis peptide bonds are ignored. Distributions for omega are shown for various secondary structures as defined by the reduced DSSP codes (loops, L+; helices, H+; and extended, E+) as well as for every protein in the database (whole database).

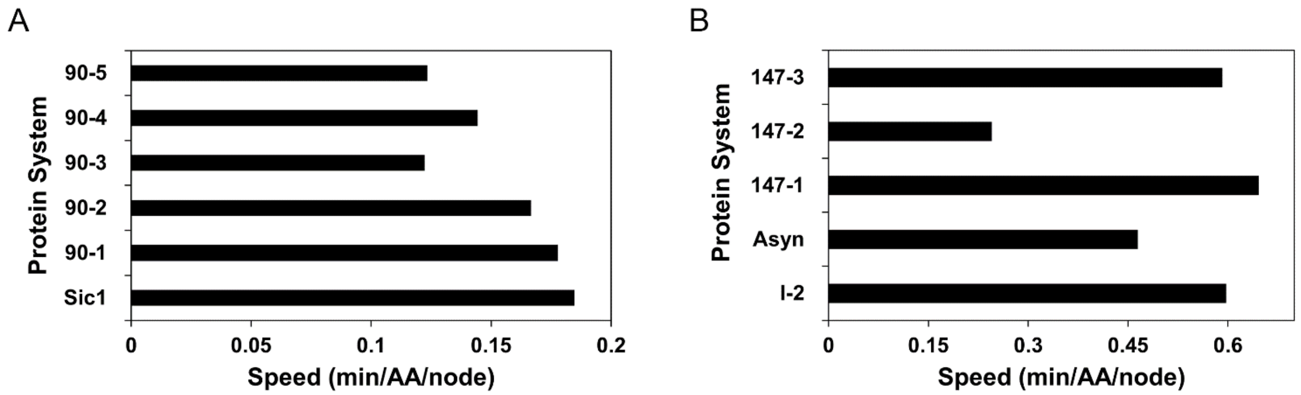


**Supplementary Figure S2. Dependence of IDPConfGen speed on sequence complexity.** Calculations of 1000 conformers of each protein were done using FASPR. **(A)** Tau (441 residue fragment) was broken up into 5 smaller fragments of 90 residue peptides sequentially (90-1: residues 1-90, 90-2: residues 91-180, 90-3: residues 181-270, 90-4: residues 271-360, 90-5*: residues 361-441). Note that 90-5* is only 80 residues. **(B)** The 441-residue fragment of Tau was broken up into 3 smaller fragments of 147 residue peptides sequentially (147-1: residues 1-147, 147-2: residues 148-294, 147-3: residues 295-441) to compare with alpha-synuclein (140 aa) and I-2 (159 aa).

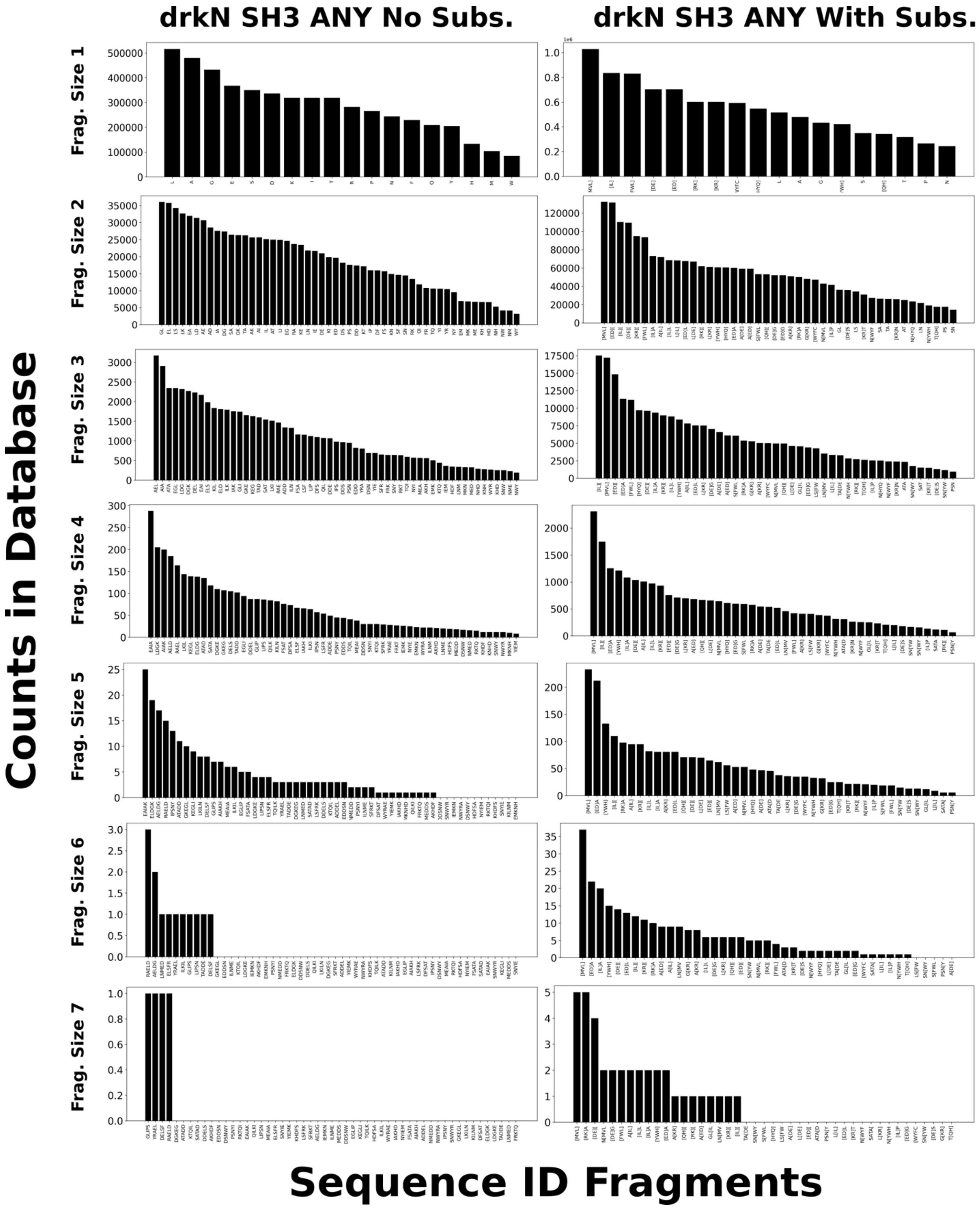


**Supplementary Figure S3. Histograms of number of segments of the drkN SH3 domain sequence for various fragment sizes present in the database.** This is based on sampling how many matches for the ANY flag (no secondary structure preference) with exact sequence or substitutions from the EDSSMat50 matrix, up to fragment sizes of 7.

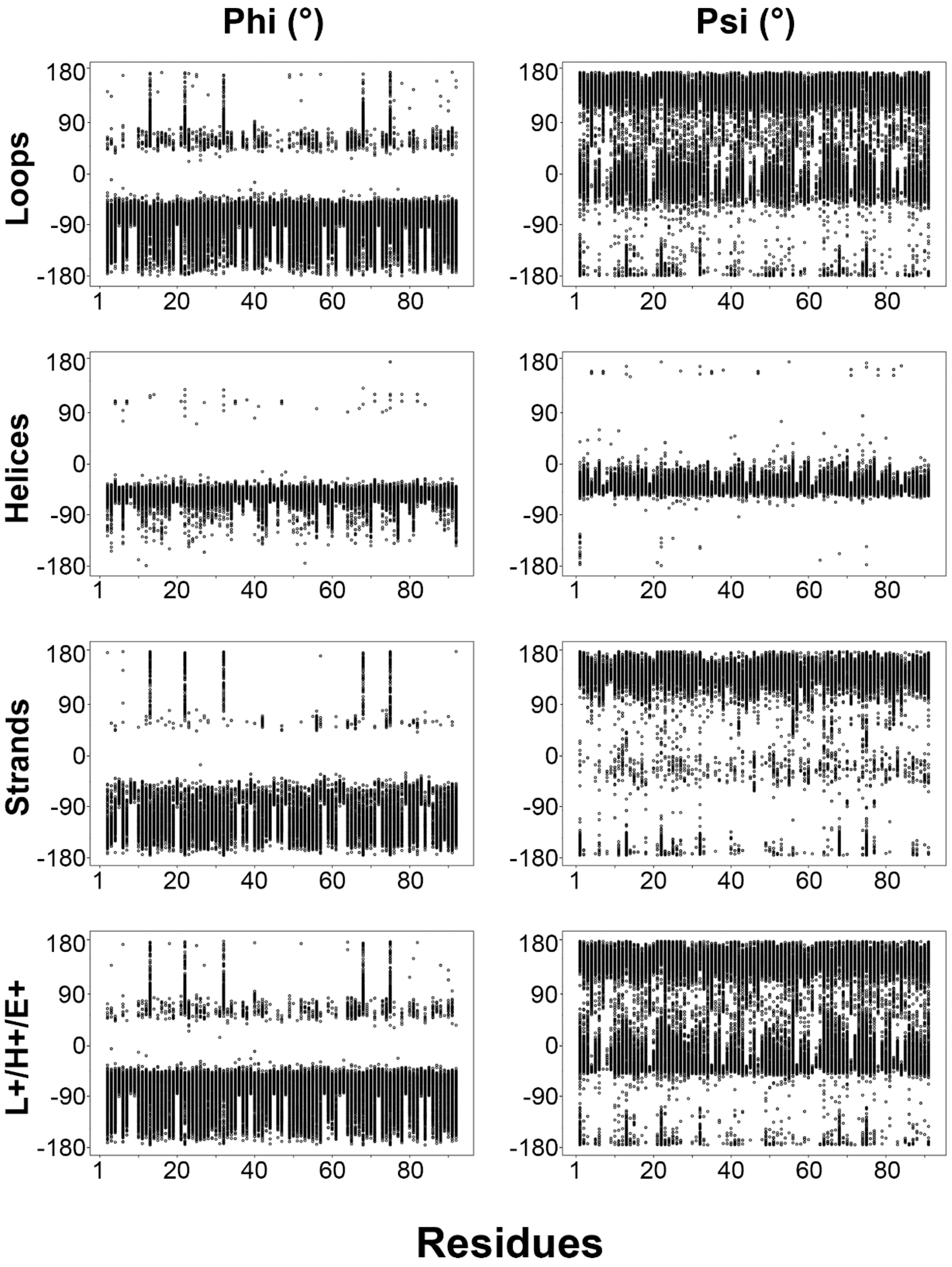


**Supplementary Figure S4. Torsion angle analysis for Sic1 conformers sampling various secondary structures.** Phi and psi torsion angle distributions of 1200-conformer ensembles of Sic1 generated with different combinations of secondary structure sampling, including loops (L), helix (H), and extended strands (E) or the combination. The column on the left **(i)** indicates Phi angles in degrees while the column on the right **(ii)** denotes Psi angles in degrees. **(A)** Generated with L. **(B)** Generated with H. **(C)** Generated with only extended/strands. **(D)** Generated with L, H, and E.

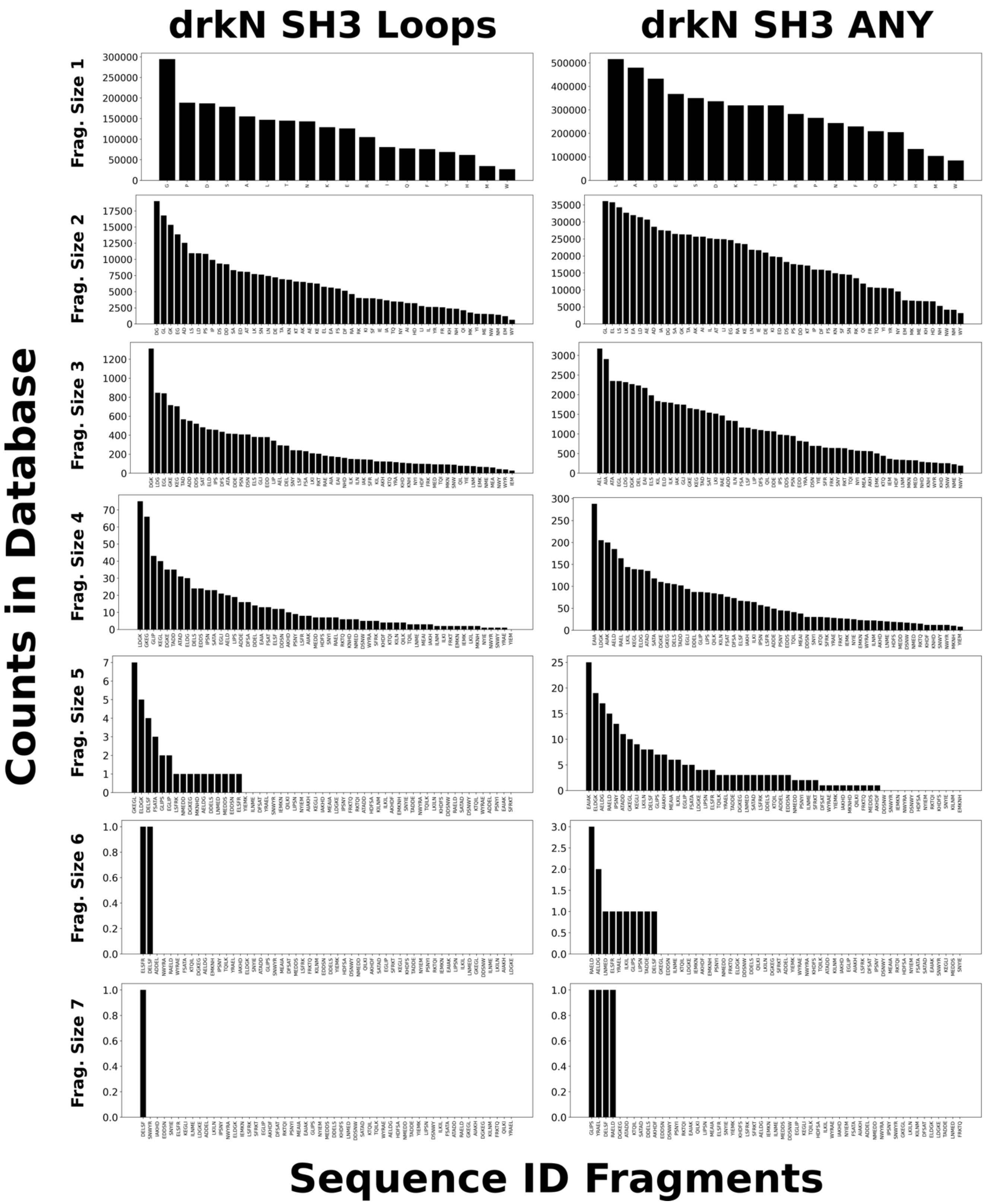


**Supplementary Figure S5. Histograms of number of segments of the drkN SH3 domain sequence for various fragment sizes present in the database, comparing sampling of loops to no preference (ANY).** This is based on sampling requiring exact sequence matches, up to fragment sizes of 7.

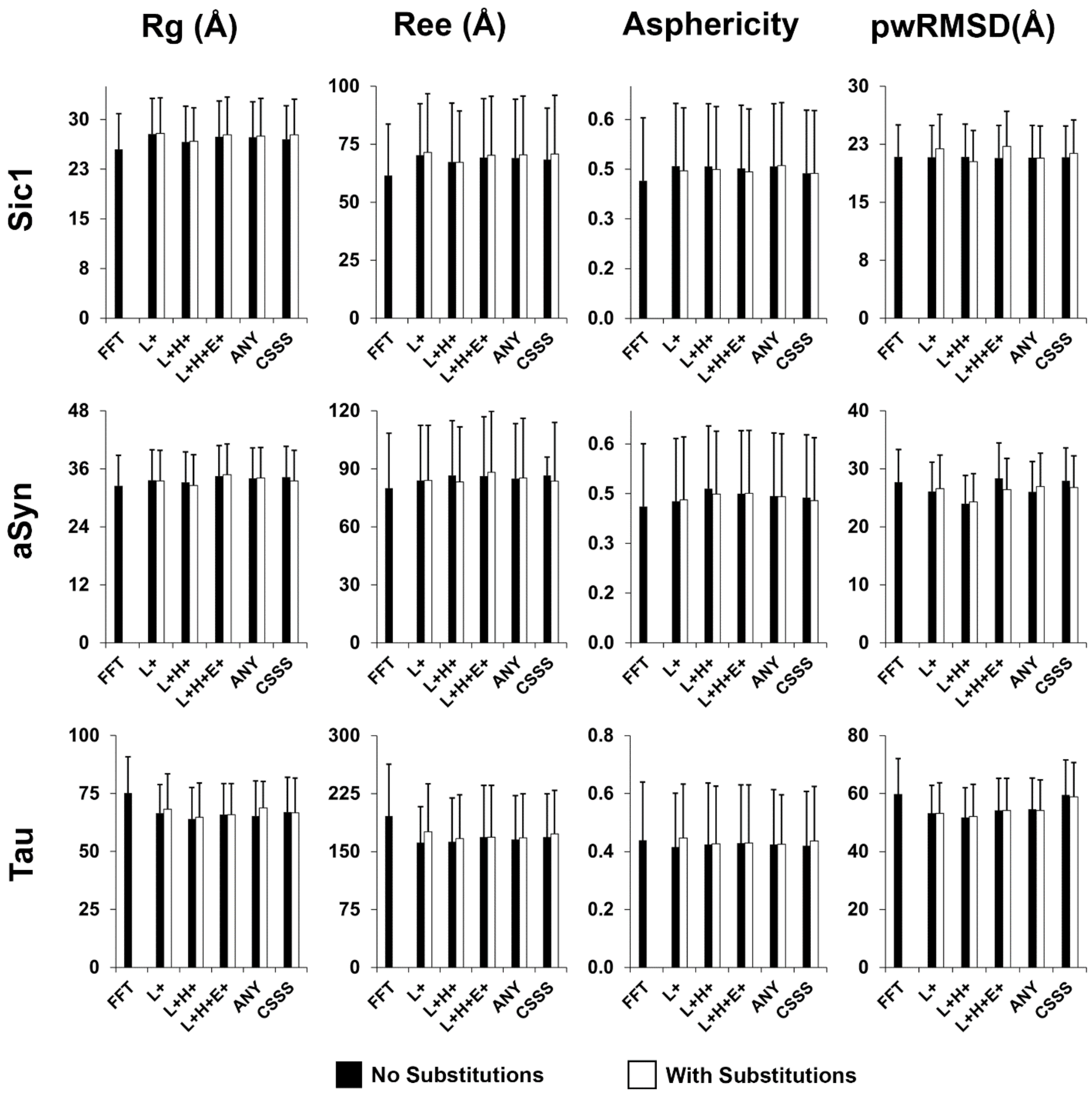


**Supplementary Figure S6**. **Diversity analysis of conformational ensembles of Sic1, aSyn and Tau**. Radius of gyration ($R_g$), end-to-end distance ($R_{ee}$), asphericity (A) and pairwise root-mean-squared-deviations of atomic positions (pwRMSDs), are shown as a function of secondary structure sampling parameters for ensembles generated with different secondary structure sampling, including loops (L+), loops and helices (L+H+), loops, helices and extended strands (L+H+E+), all torsion angles agnostic to secondary structure (ANY) and biased by $\delta$2D chemical shifts (CSSS), or with FastFloppyTail (FFT), for Sic1 (row 1), alpha-synuclein (row 2), and Tau (row 3). The error bars represent the standard deviations. For all but Tau, 1000-conformer ensembles were generated and analyzed. For Tau, variable number of conformers were generated and analyzed, with FFT, L+, L+H+, L+H+E+, ANY, CSSS having 500, 240, 200, 500, 350, 480 conformers, respectively.

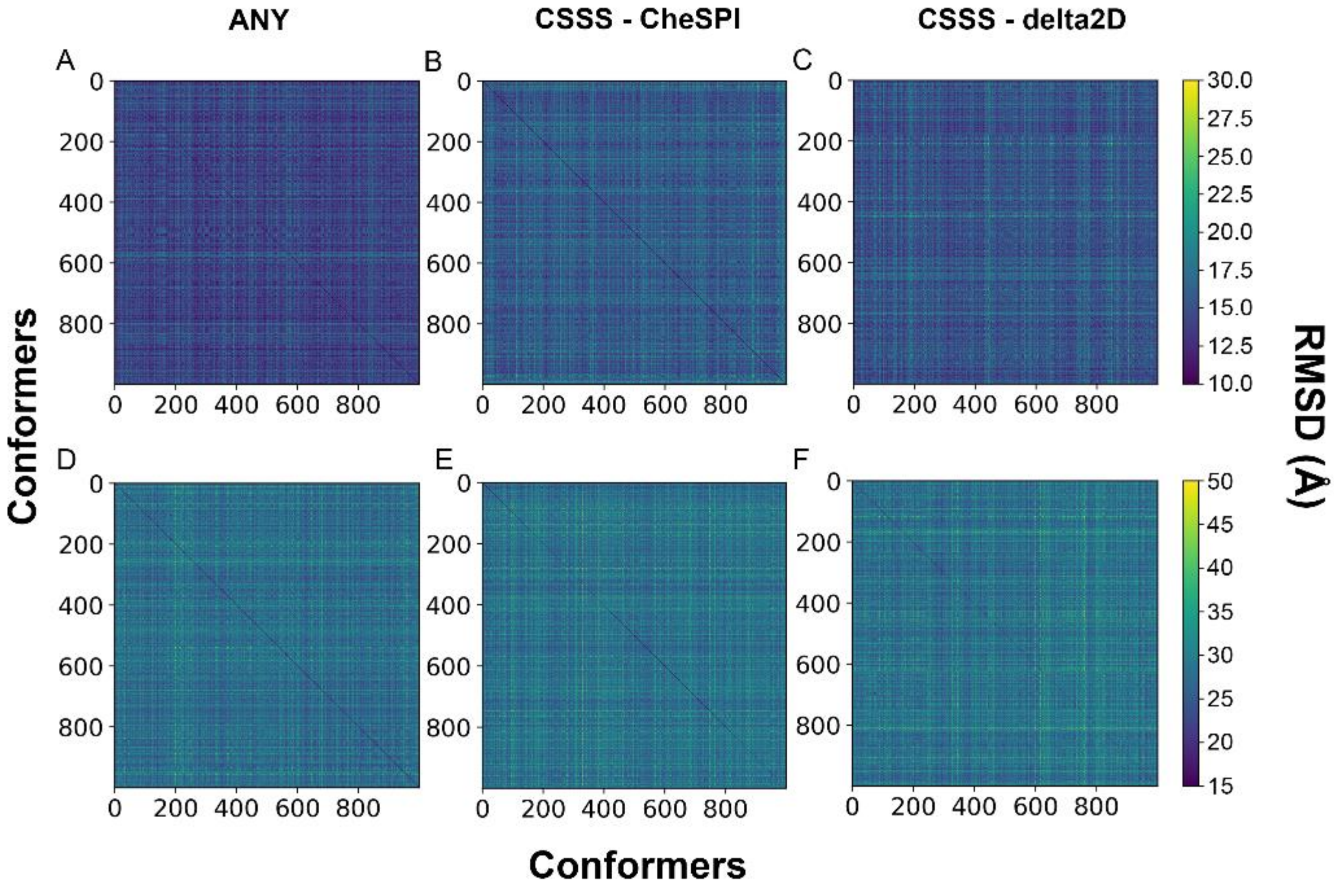


**Supplementary Figure S7. Pairwise RMSD distance matrices**. Ensembles of the drkN SH3 domain unfolded state (A, B, C) and I-2 (D, E, F) with 1000 conformers each, generated using different secondary structure sampling methods, ANY and CSSS with either CheSPI or δ2D to bias sampling. differences between $C_{alpha}$-$C_{alpha}$ distance matrixes ($P < 0.05$ from a Mann-Whitney U test).

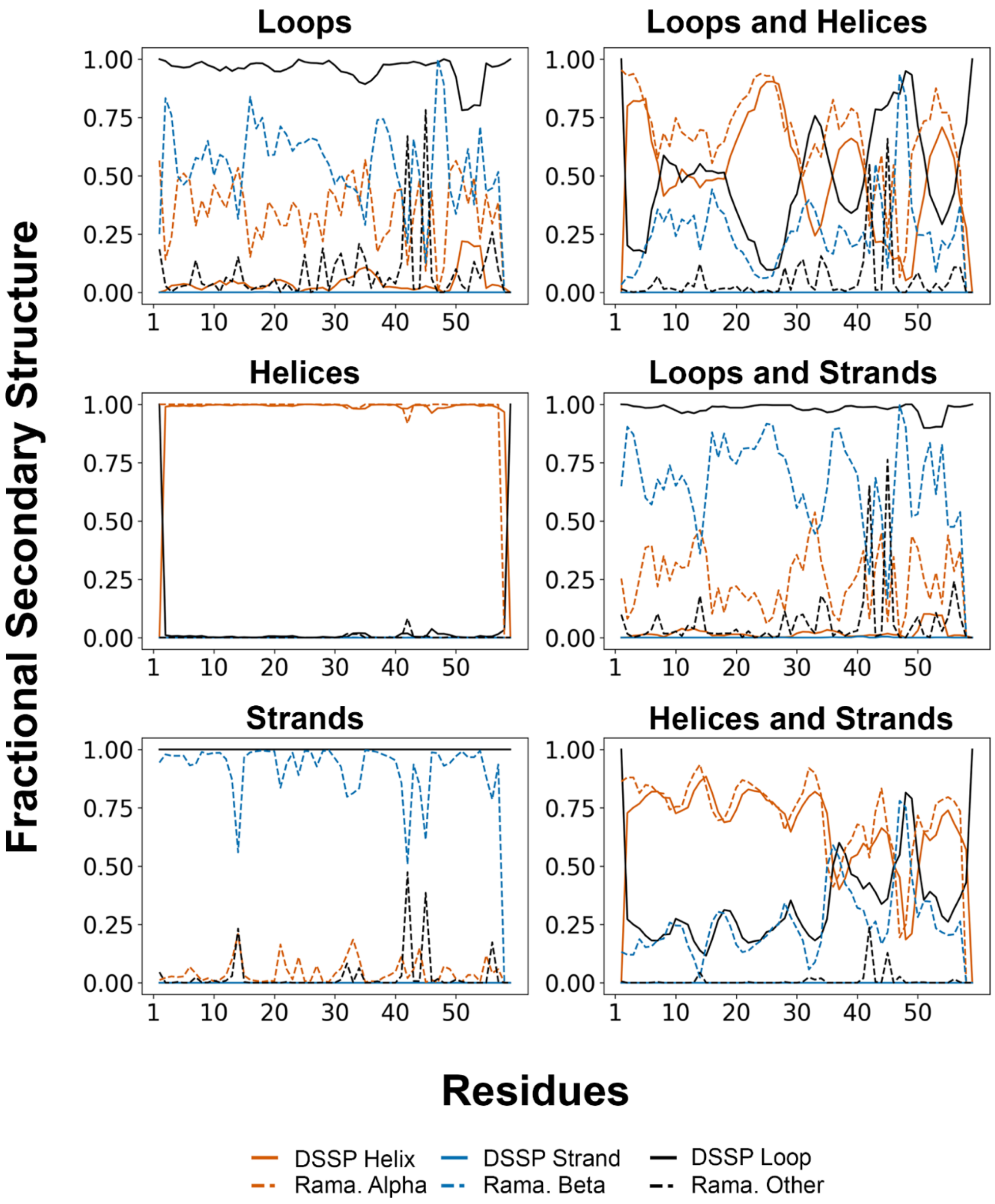


**Supplementary Figure S8. Fractional secondary structure in ensembles of the drkN SH3 domain unfolded state.** Analyses were performed on sets of 1000-conformer pools of the drkN SH3 domain unfolded state generated with different combinations of secondary structure sampling consisting of loops, helix, and extended strands. Orange indicates alpha-helix for DSSP (solid) and the alpha-region on the Ramachandran diagram (dashed). Blue indicates extended strand for DSSP (solid) and beta-region on the Ramachandran diagram (dashed). Black indicates coil/loop for DSSP (solid) and other regions on the Ramachandran diagram (dashed).

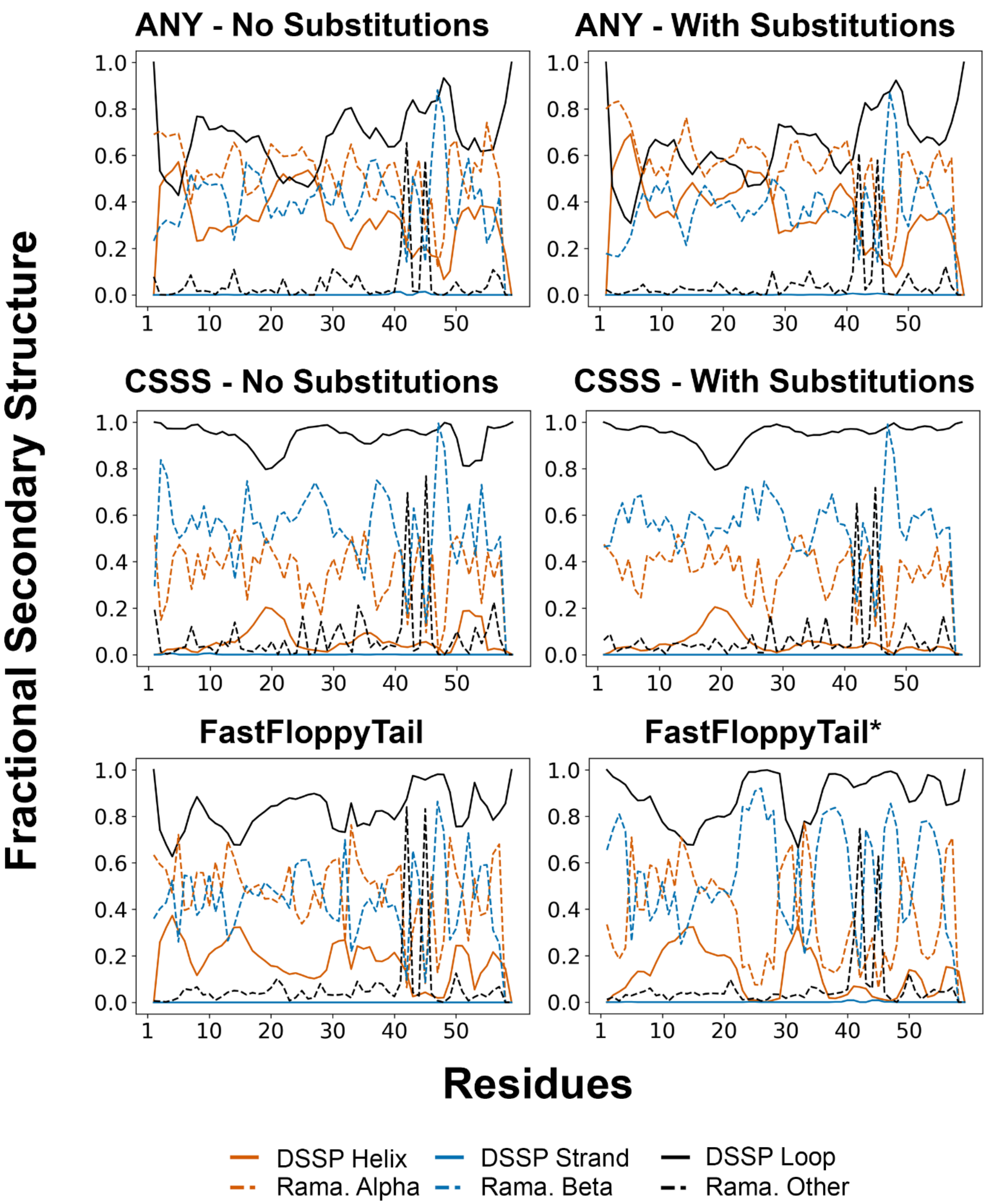


**Supplementary Figure S9. Fractional secondary structure in calculated ensembles of the drkN SH3 domain unfolded state generated with IDPConformerGenerator using ANY and CSSS and FastFloppyTail**. Analyses were with ensembles of 1000 conformers, a backbone energy threshold of 100 kJ, and side chains added on with MC-SCE with 128 trials per conformer in simple mode. Sequence substitutions were from the EDSSMat50 substitution matrix rows 5, 3, and 2. * is for the standard protocol which for this case treats the protein as a mixture of ordered and disordered, while the other is for a modified protocol in which the protein is considered to be fully disordered.

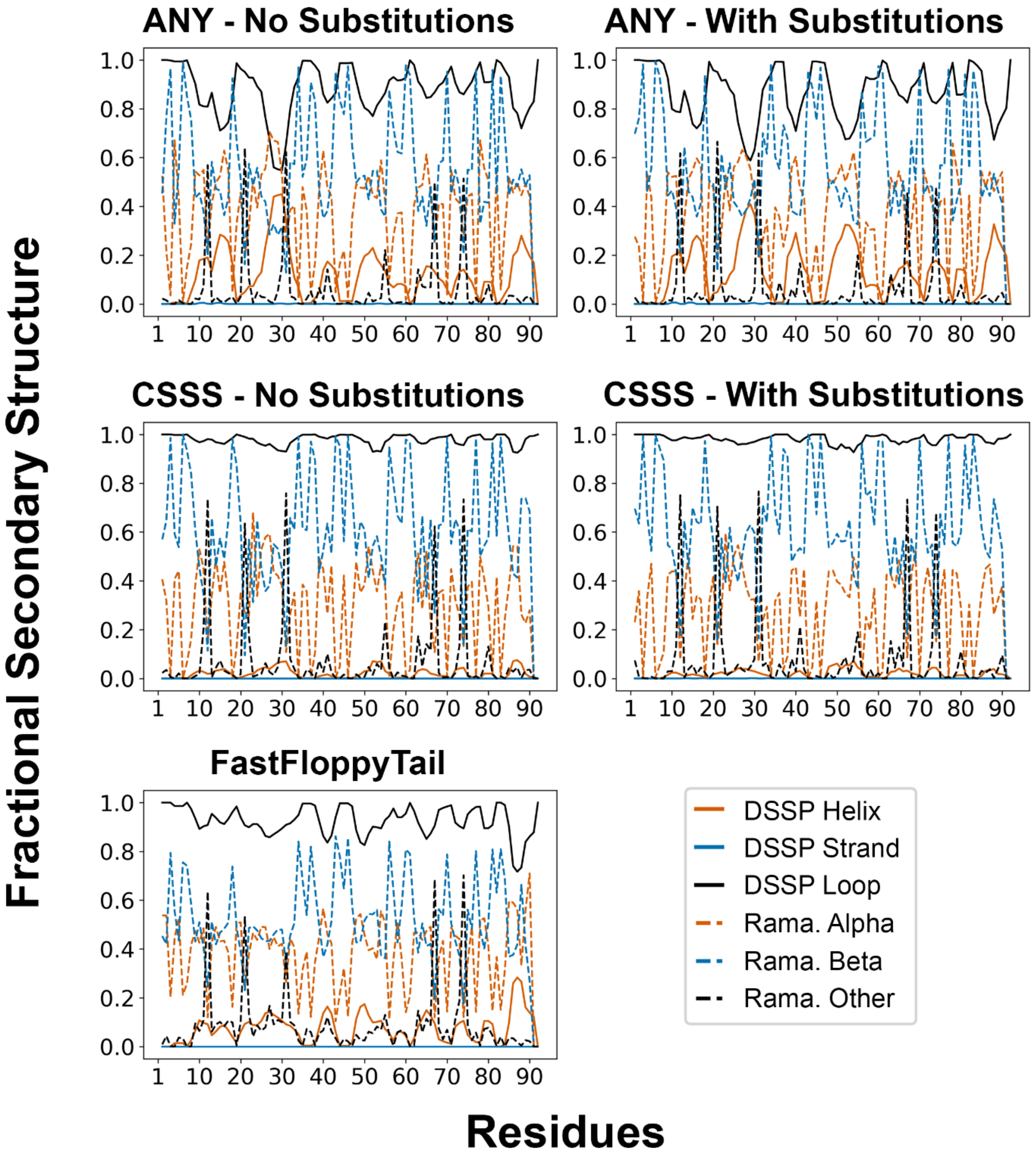


**Supplementary Figure S10. Fractional secondary structure in calculated ensembles of Sic1 generated with IDPConformerGenerator using ANY and CSSS and FastFloppyTail.** Analyses and plots are as for Supplementary Figure S9.

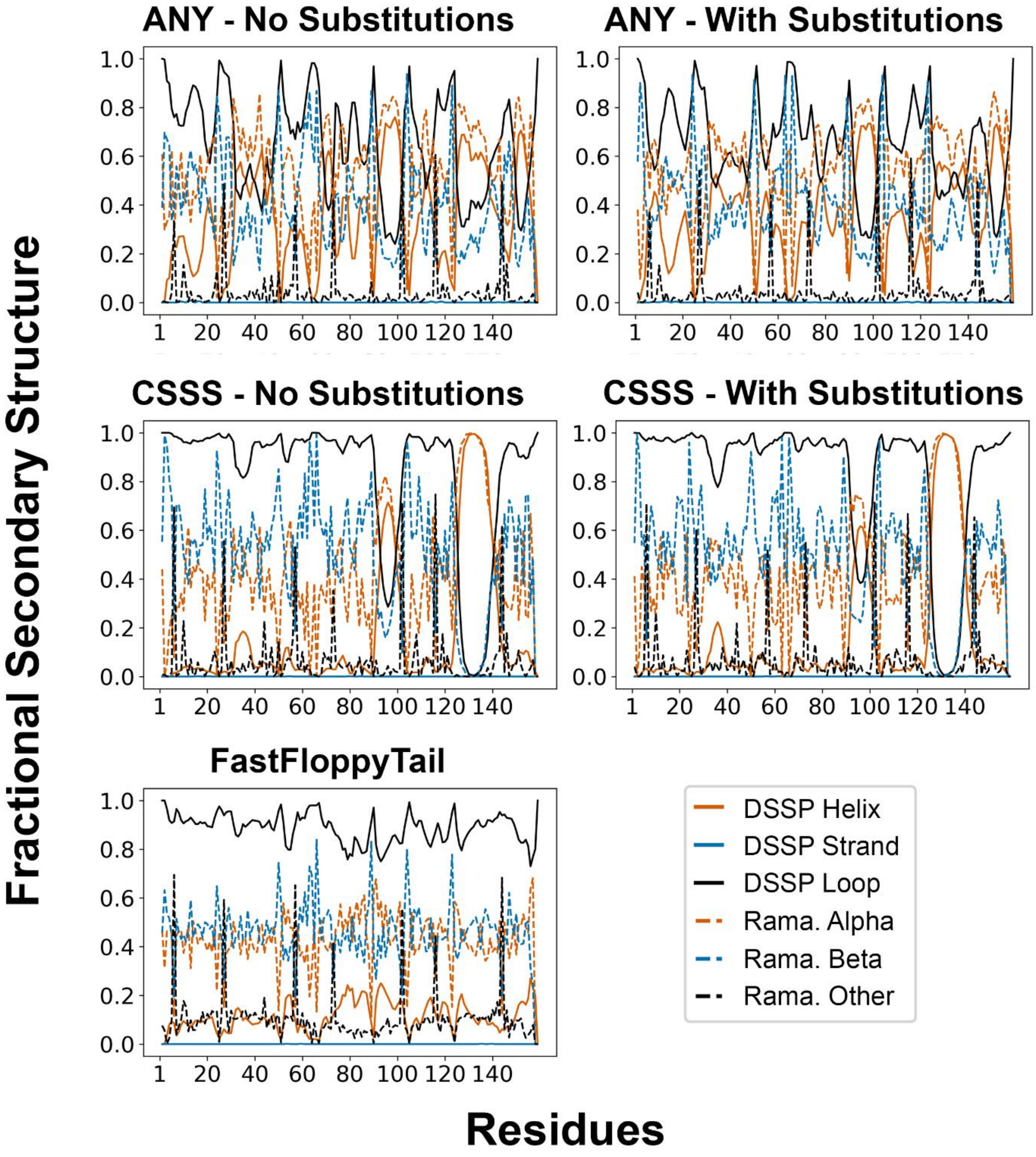


**Supplementary Figure S11. Fractional secondary structure in calculated ensembles of alpha-synuclein generated with IDPConformerGenerator using ANY and CSSS and FastFloppyTail.** Analyses and plots are as for Supplementary Figure S9.

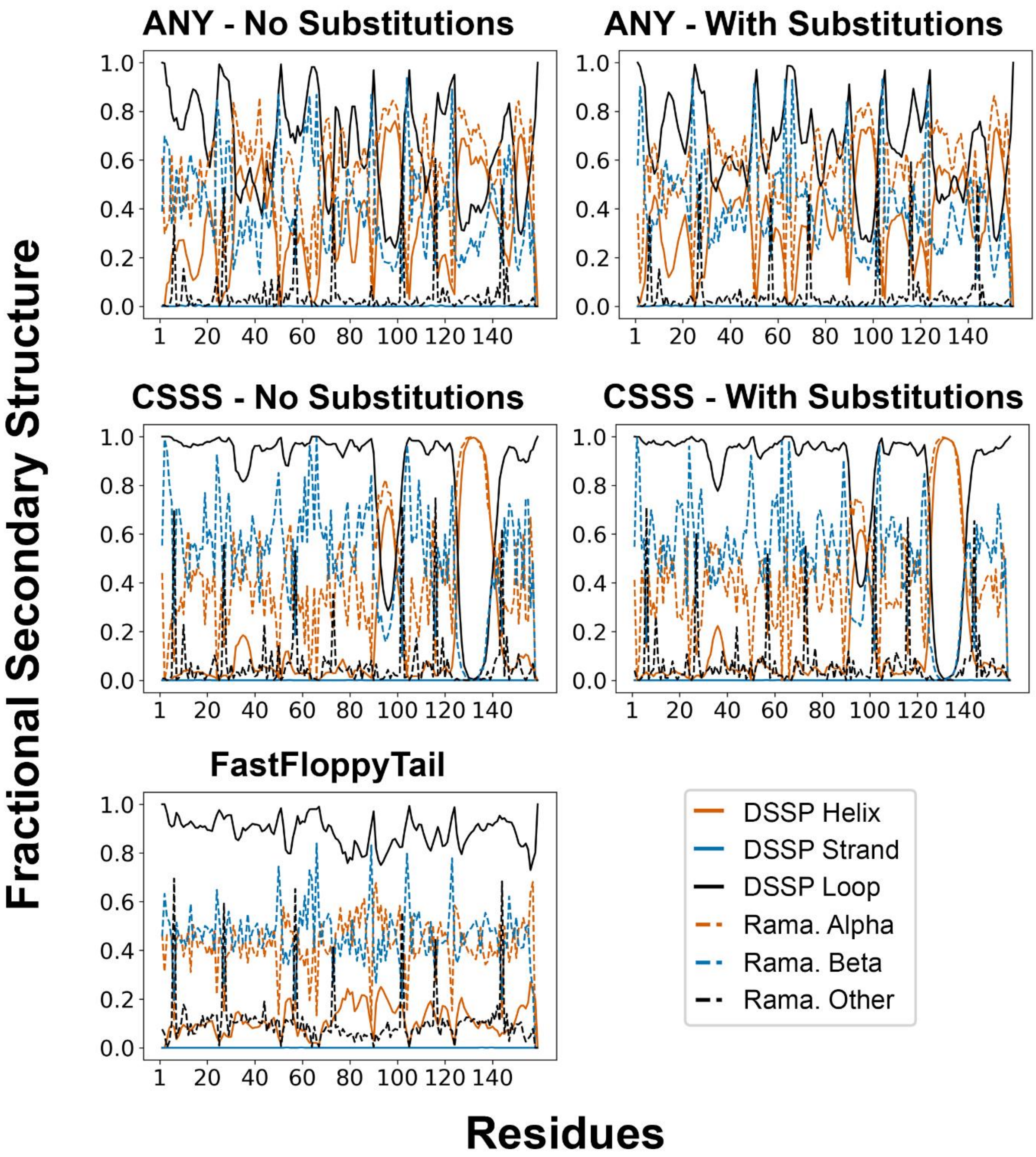


**Supplementary Figure S12. Fractional secondary structure in calculated ensembles of Inhibitor-2 generated with IDPConformerGenerator using ANY and CSSS and FastFloppyTail.** Analyses and plots are as for Supplementary Figure S9.

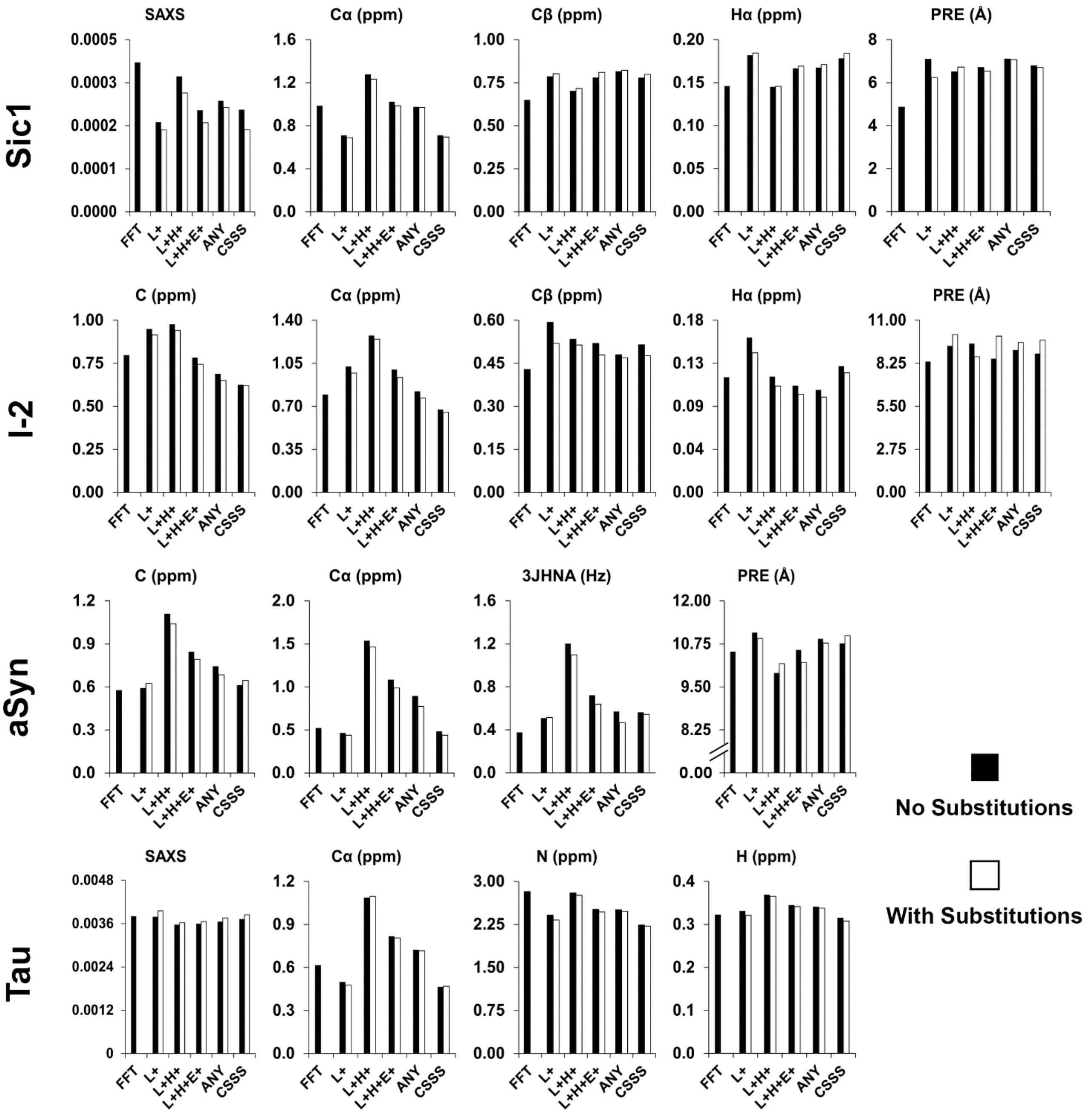


**Supplementary Figure S13. Root-mean squared deviations (RMSDs) of back-calculated values from conformational ensembles to experimental data.** Analyses of ensembles generated using various secondary structure sampling and using FastFloppyTail (FFT) for Sic1, I-2, alpha-synuclein and Tau. RMSDs are given for SAXS, chemical shifts (Carbonyl, Calpha, Cbeta, Amide Nitrogen, Halpha), PRE, $^3J_{HN\text{-}HA}$, and NOE if available. For all but Tau, 1000-conformer ensembles were generated and analyzed. For Tau, variable number of conformers were generated and analyzed, with FFT, L+, L+H+, L+H+E+, ANY, CSSS having 500, 240, 200, 500, 350, 480 conformers, respectively.

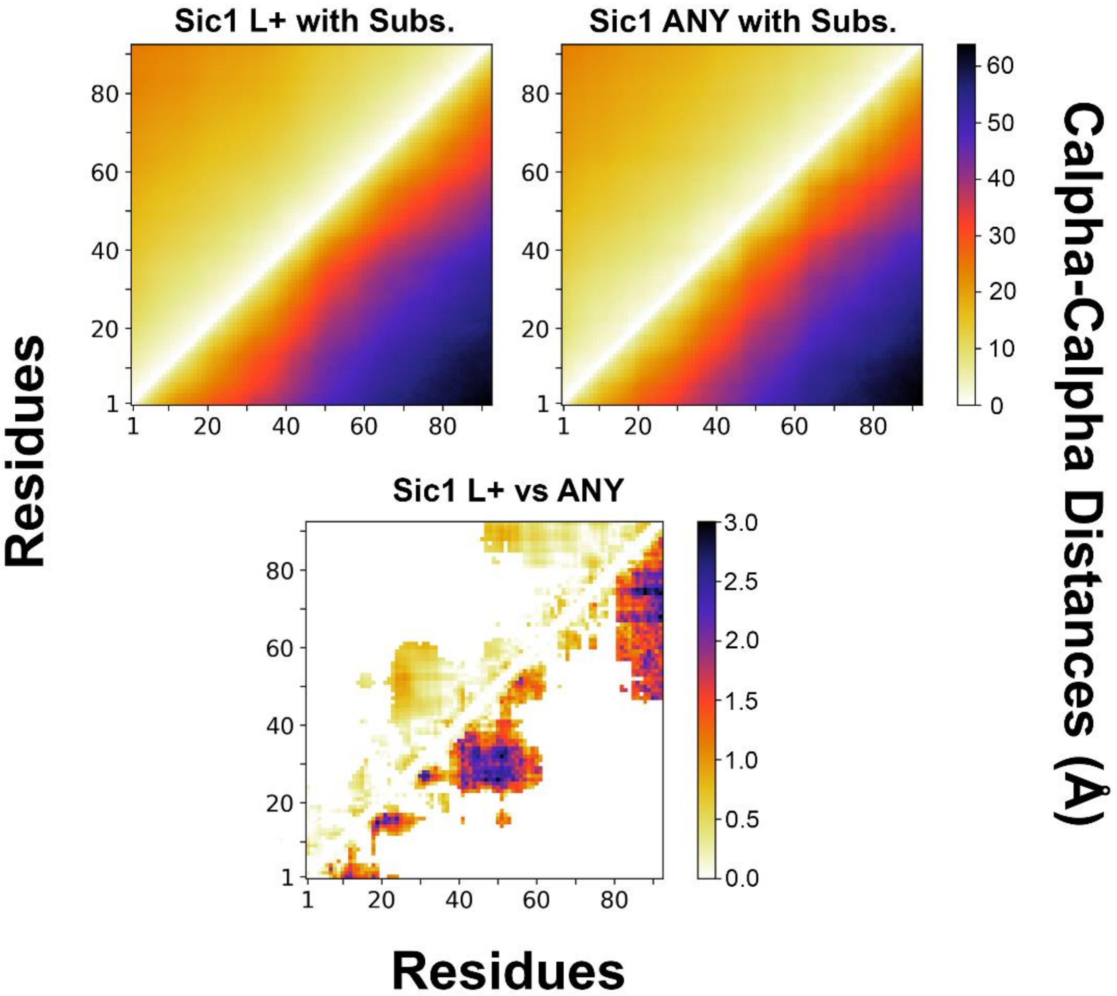


**Supplementary Figure S14. Analysis of tertiary contacts for Sic1 ensembles with varying secondary structure sampling preferences.** (top row) $C_{alpha}$-$C_{alpha}$ distance matrixes (lower) with deviations (upper) for 1000-conformer ensembles of Sic1 generated with the loops only and ANY flag for secondary structure, with substitutions from columns 5, 3, and 2 of the EDSSMat50 amino acid substitution matrix and with variable fragment lengths. (bottom row) Significant differences between the two $C_{alpha}$-$C_{alpha}$ distance matrixes (lower) and deviations (upper), ($P < 0.05$ from a Mann-Whitney U test).

**SUPPLEMENTARY TABLES**

Supplementary Table S1A: EDSSMat50 matrix for amino acid substitutions.

Supplementary Table S1B: Primary sequences of all protein systems tested.

Supplementary Table S1C: PDB IDs that contain fragments from our test protein systems.

Supplementary Table S1D: Counts of sequence ID matches for different fragment sizes of the drkN SH3 domain in our database.

Supplementary Table S2A: Speeds for calculated ensembles.

Supplementary Table S2B: Benchmarking RAM Size and MC-SCE Trials.

Supplementary Table S2C: Comparison of percentage of clash free conformers for alpha-synuclein and inhibitor-2 at different backbone energy thresholds.

Supplementary Table S2C: Total time required to prepare input files for IDPConformerGenerator and FastFloppyTail.

Supplementary Table S3: Radii of gyration, end-to-end distances, asphericity values and pairwise RMSDs for calculated ensembles.

Supplementary Table S4A: Deviation from experimental restraints and ENSEMBLE scores for calculated ensembles.

Supplementary Table S4B: Deviation from experimental restraints and X-EISD scores for calculated ensembles.
